# Tracking single cell evolution via clock-like chromatin accessibility

**DOI:** 10.1101/2022.05.12.491736

**Authors:** Yu Xiao, Wan Jin, Lingao Ju, Jie Fu, Gang Wang, Mengxue Yu, Fangjin Chen, Kaiyu Qian, Xinghuan Wang, Yi Zhang

## Abstract

Single cell chromatin accessibility sequencing (scATAC) reconstructs developmental trajectory by phenotypic similarity. However, inferring the exact developmental trajectory is challenging. Here, we show a simple, accurate and phenotypic-neutral measure of cell developmental hierarchy – the fraction of accessible clock-like loci. As cells undergo mitosis, the heterogeneity of chromatin accessibility on clock-like loci is reduced, providing a measure of mitotic age. We developed a method, EpiTrace, that counts the fraction of opened clock-like loci from scATAC data to determine cell age and perform lineage tracing. EpiTrace works in various cell lineages and animal species, shows concordance with known developmental hierarchies, correlates well with DNA methylation-based clocks, and is complementary with mutation-based lineage tracing, RNA velocity, and stemness predictions. Applying EpiTrace to scATAC data revealed a multitude of novel biological insights with clinically relevant implications, ranging from hematopoiesis, organ development, tumor biology and immunity to cortical gyrification. Our work discovered a universal epigenomic hallmark during cellular development, which facilitates the study of cellular hierarchies and organismal aging.

## Introduction

Single cell chromatin accessibility sequencing (scATAC) is a powerful technique for interrogating the epigenomic landscape at single cell resolution^1^. However, inferring the exact developmental trajectory from scATAC data is challenging. While excellent tools such as RNA velocity, stemness prediction and metabolic labeling^2–5^ exist for determining the cell evolution trajectory on the manifold of phenotypes – usually described as Waddington’s landscape – for single cell RNA-seq datasets, no comparable methods exist for scATAC. State-of-the-art, similarity-based lineage deduction methods^6^ would be limited when phenotypes are fluidic, such as in dedifferentiation or oncogenesis. On the other hand, mutation-based lineage tracing methods, for example, using mitochondrial SNPs^7–9^, which track the phylogeny of cells over divisions, are highly accurate, yet their temporal resolution is restrained by the low natural mutation rate.

The concept of mitotic age refers to the accumulative counts of mitosis a cell undergoes after fertilization – or the ground state of cell division. While the first proposed mitotic age biomarker – telomere length – was genetic^10, 11^, the concept quickly extended to epigenetic replication errors on DNA methylation (DNAm)^12, 13^. During DNA replication, epigenetic covalent modifications are not faithfully replicated to the daughter strand, resulting in stochastic DNAm changes. Stochastic DNAm fluctuation has been applied to infer the mean mitotic count of the cell population^14–16^. On the population scale, irreversible, stochastic DNAm changes were thought to be underlying age-associated DNAm changes. A vast number of studies have documented age-associated DNAm changes^17^, including hypomethylation and hypermethylation, in specific genomic regions. We term such genomic regions clock-like differential methylation loci (ClockDML) because their DNA methylation exhibited timekeeper-like behavior.

The DNAm-based regression model predicts the age of biological samples with extremely high precision in many organisms^18–24^ and correlates with rejuvenation or accelerated aging in various scenarios^25^. A similar age association of DNA methylation was conserved across mammalian species in homologous genomic regions^26^, suggesting that it is controlled by a defined, possibly functional molecular mechanism^27^. Interestingly, a predictor model has been built to estimate the mitotic age of samples from the DNAm state of a defined set of CpG loci^28, 29^, indicating that mitosis is associated with clock-like DNA methylation changes at specific genomic loci. Introducing population statistics into the DNAm-based age prediction model enabled single cell age prediction from single cell methylation sequencing data^30, 31^, suggesting that clock-like DNAm changes are not merely a statistical phenomenon at the population scale but also occur at the single cell level.

Based on the intrinsic link between chromatin accessibility and DNA methylation^32–37^, we hypothesized that age-dependent DNA methylation could either result from or result in chromatin accessibility changes at clock-like differentially methylated loci. If so, the derived mitotic age of single cells from scATAC data would serve as a powerful tool to delineate developmental trajectory. In theory, mitotic age is a “timekeeper” tracker: the mitotic age of an ancestor cell is lower than that of its progeny, and cells originating earlier in time should show lower mitotic age than those originating later. Such a measure of cell age, if exists, would provide a precise temporal reference of the cell birth sequence to help delineate the developmental trajectories in a complex organism.

To develop a mitotic age estimator for scATAC data, we determined ClockDML across the human genome and characterized the chromatin accessibility changes in these loci associated with cell mitotic age. The heterogeneity of chromatin accessibility at these loci reduces across cell division. Through genomic synteny mapping, we showed that the age-dependent chromatin accessibility is conserved on these loci across evolution. Such clock-like chromatin accessibility is independent from DNA methylation. Even in species without active DNA methylation, clock-like chromatin accessibility exists on these loci. Hence, we term these region clock-like loci. We leveraged this phenomenon to develop a computational framework, *EpiTrace,* which infers cell mitotic age from scATAC data by counting the opened fraction of clock-like loci in single cells.

## Results

### Chromatin accessibility on ClockDML enables cell age estimation

Although the molecular mechanism that generates age-associated DNAm changes is unclear, it is possible that the methylation state of ClockDML might be affected by chromatin accessibility. Alternatively, the methylation state of ClockDML might reversely regulate chromatin accessibility. In either case, the chromatin accessibility on ClockDML could be used to deduce cell age (Figure 1a). However, the dynamics of chromatin accessibility on ClockDML during aging are currently unknown.

**Figure 1.**
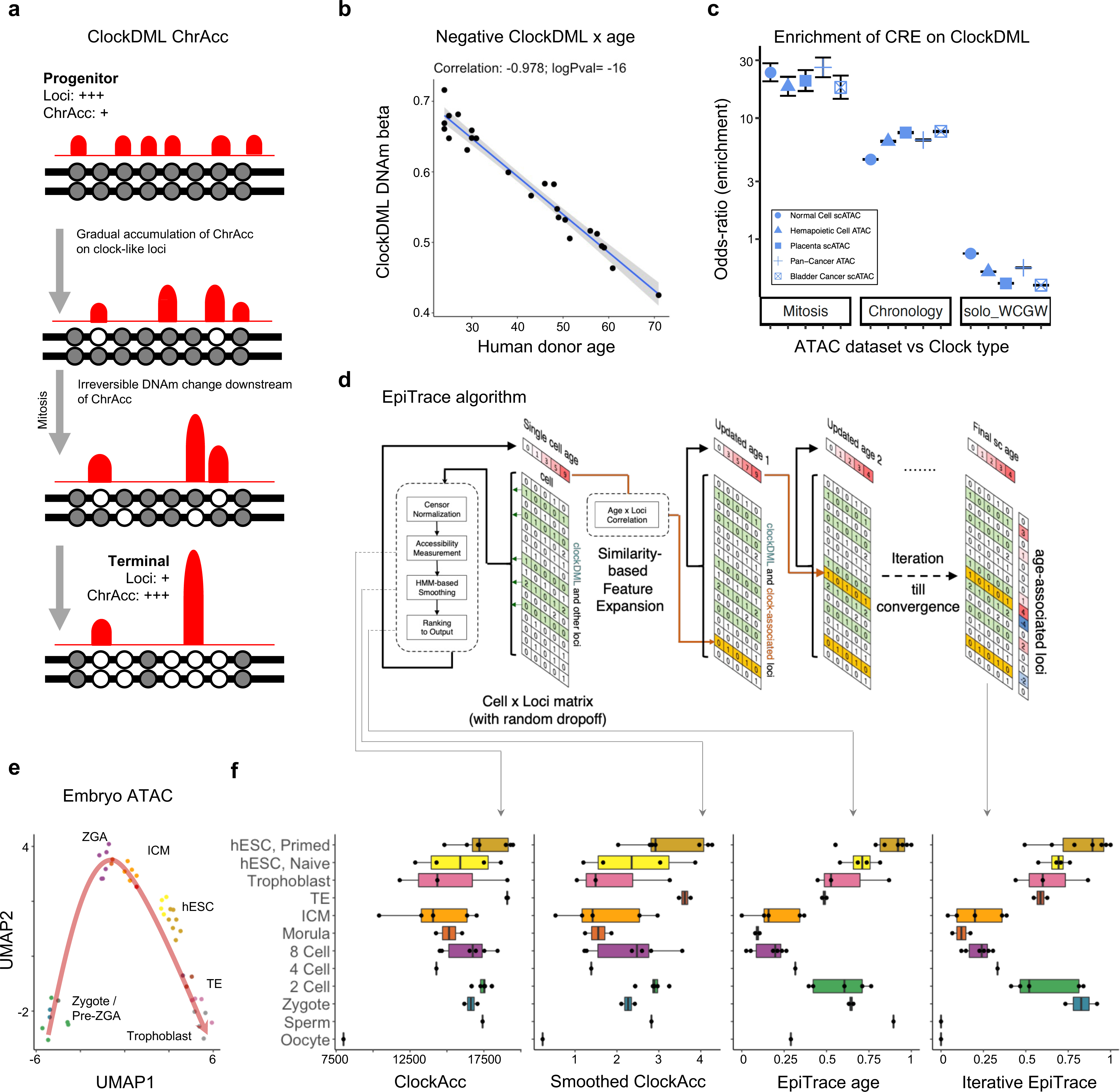
ChrAcc change associated with irreversible DNA methylation drift on ClockDML enables cell age estimation. **(a)** Schematic diagram of the underlying epigenetic mechanism of cell mitotic age tracing using chromatin accessibility (ChrAcc) on Clock-like differentially methylated loci (ClockDML). **(b)** Correlation between the DNA methylation level on G8-group and sample age in human PBMC. **(c)** Enrichment of each class of ATAC peaks in mitosis-associated ClockDML (Mitosis), actual age-associated ClockDML (Chronology), and solo-WCGW loci. **(d)** Overview of the *EpiTrace* algorithm. **(e)** UMAP projection of the human early embryonic development scATAC dataset. **(f)** The total chromatin accessibility on ClockDML (ClockAcc), HMM-smoothened ClockAcc, initial and iterative EpiTrace ranking result (EpiTrace Age) corresponding to the embryonic dataset. Abbreviations: ZGA: zygotic genome activation; ICM: inner cell mass; ESC: embryonic stem cell; TE: trophectoderm.

Literature-documented ClockDML^18, 25, 26, 38^ were found mainly with methylation-specific microarrays and thus represent only a tiny fraction of possible genome-wide DNAm variation during aging. However, ATAC -- and more so scATAC -- data are too sparse to fully cover these loci. To enable accurate tracking of cell mitotic age by ATAC signal, we determined 126,420 ClockDML in the human genome by bisulfite capture sequencing the CpG island regions of peripheral blood mononuclear cells from a panel of donors of different ages and determined the correlation coefficient between age and the methylation level (beta) on each locus (Supplementary Dataset 1: ClockDML). The DNAm status of these ClockDML showed excellent correlation with age in the training cohort (Figure 1b). Both the general linear model (GLM) and probability model (TimeSeq)^31^ built upon beta values of our ClockDML predict donor age with good precision in an additional validation cohort of samples (R = 0.85 (GLM)/0.7998 (TimeSeq), Supplementary Figure 1a-c), indicating that the DNAm status of these loci stably drifts over age. Functional annotation suggests that ClockDML are enriched in the open, accessible chromatin region of the genome across different cell types and organs^38–41^ (Figure 1c). In correlation with DNAm status, the fraction of opened ClockDML (hereinafter in short: ClockAcc) shifts in correlation with cell aging (Supplementary Notes and Supplementary Figures 2-5, GSE74912^42^, GSE89895^43^, GSE179606^44^). We established an algorithm, EpiTrace, that predicts sample age by counting the fraction of opened ClockDML in bulk ATAC-seq datasets (Supplementary Notes). Validation experiments using bulk ATAC-seq of FACS-sorted blood cells, iPSC induction experiments, and native immune cells showed that EpiTrace accurately predicted sample age in concordance with known developmental trajectories (Supplementary Notes and Supplementary Figures 2-5). We then adopted the algorithm for single cell ATAC-seq data. In brief, a reference ClockDML set was provided to the algorithm. ClockAcc, the total chromatin accessibility on this reference ClockDML set, is measured for each cell. The measurement was performed using a HMM-mediated diffusion-smoothing approach, borrowing information from similar single cells to reduce noise in the single cell measurement. Cells were clustered via correlation of top variable ATAC peaks to form a cellLcell similarity matrix. The matrix was then used for diffusion-regression iterations of ClockAcc until convergence. The regularized and smoothened ClockAcc were then ranked. Such rank denotes the relative mitotic (replicational) cell age. To overcome sampling sparseness in scATAC-seq, we reasoned that it might be unnecessary for age-dependent chromatin accessibility to always be accompanied by DNAm changes (Supplementary Notes). Thus, we perform stepwise iterations by extracting additional open regions with a high correlation coefficient with estimated single cell age and then include them together with the reference loci to form a new set of clock-like loci for the next round of analysis until the age prediction converges (Figure 1d). Importantly, during the computation of EpiTrace algorithm, known sample age information is not required. The algorithm simply leverages the fact that heterogeneity of given reference ClockDML reduces during cell replication, then uses such information as an intermediate tool variable to infer cell age.

We first validated the EpiTrace algorithm on *in vitro* models. Chromatin accessibility in single mouse cells was profiled with the SHARE-seq assay (Supplementary Figure 6), and DNAm age (in batch) was determined by DNAm sequencing (Supplementary Figure 1d-g). In asynchronized immortal mouse embryonic fibroblast (MEF) cells, progression in the cell cycle results in a reduction in EpiTrace predicted age (Supplementary Figure 7), suggesting that EpiTrace tracks an epigenomic modification that dilutes genome replication. In primary MEF (pMEF) cells, this phenomenon persists (Supplementary Figure 8a-b, d, h). However, as the cells were passaged *in vitro*, the EpiTrace age stably increased (Supplementary Figure 8b, i). Such a mitosis age-dependent increase in EpiTrace age overwhelms the genome replication-mediated dilution effect (Supplementary Figure 8) and correlates well with DNAm-based age prediction of the same batch of samples (Supplementary Figure 9). Finally, for cells that are pharmacologically blocked in a specific cell cycle (GSE65360^1^), EpiTrace age increases from G1 to S and G2/M phase (Supplementary Figure 10), suggesting that accumulation of error during copying of epigenomic modification results in an increase in EpiTrace age prediction over mitosis. In large *in vivo* single cell datasets without cell phase synchronization, the cell cycle had little effect on EpiTrace age prediction (Supplementary Figure 11, GSE163579). Together, these data indicate that EpiTrace reports mitosis age.

As a proof-of-concept, we gathered ATAC data from various studies of early human embryonic development from gametes to blastula^45, 46^ (Figure 1e and Supplementary Figure 12, PRJNA494280, PRJNA394846), which were generated from only a few cells each, and subjected them to EpiTrace analysis without batch correction (Figure 1f). The total ClockAcc in sample positively correlates with known cell mitotic age. While the initial EpiTrace age prediction is noisy, iterative optimization improved the signal-to-noise ratio to draw a biologically plausible trajectory of age resetting during early embryonic development: starting from zygote, cell mitotic age gradually reduces to near ground state at the time of zygotic genome activation at morula, before its rebound in inner cell mass (ICM) and trophectoderm (TE).

### Extending the EpiTrace algorithm application across cell types and animal species

For many cell types and animal species, ClockDML has not been experimentally determined. The fact that ClockDML derived from human PBMCs could not only be used to predict the sample age of human blood cells but also cells of the non-hematopoietic lineage (Figure 1 and Supplementary Figure 5) suggests that clock-like chromatin accessibility on the ClockDML genomic region might be universal across cell lineages. To test whether we could extend known ClockDML to other species or cell types for EpiTrace prediction, we mapped human ClockDML to the mouse genome using genomic synteny and computed EpiTrace age for the mouse scATAC dataset using mouse ClockDML or “human-guided” clock-like loci (Supplementary Figure 13a). We found that the EpiTrace prediction results using the reference “human-guided” clock-like loci closely approximated the prediction results using the reference mouse ClockDML (R = 0.81, Supplementary Figure 13b, GSE137115^47^).

To further validate the concordance between EpiTrace prediction starting from different reference loci, we tested a mouse scATAC dataset of T cells under chronic or acute virus infection (Supplementary Figure 14a, GSE164978^48^). EpiTrace age prediction using the mouse reference ClockDML agrees with the known developmental trajectory of these immune cells (Supplementary Figure 14b). In concordance from its tissue-of-origin, clock-like loci inferred from genomic synteny of human PBMC ClockDML overlaps with known immune cell exhaustion genes such as Pdcd1, Havcr2, Tox and Eomes, whilst mouse ClockDML^49^ (derived from pan-body DNAm interrogation) does not (Supplementary Figure 14c). However, single cell age inferred by EpiTrace with mouse ClockDML as the reference loci correlates well with that inferred with “human-guided” clock-like loci as reference (Supplementary Figure 14d). The association of ATAC peak chromatin accessibility and single cell ages from the two predictions shows extremely high concordance (R = 0.92, Supplementary Figure 14e), with the identification of many known immune exhaustion genes being positively correlated with cell age (Supplementary Figure 14e). Such correlation is not dependent on whether the loci are previously overlapping with a reference clock-like loci (Supplementary Figure 14f). Furthermore, peaks overlapping with both “human-guided” clock-like loci and mouse ClockDML showed the greatest age-dependent chromatin accessibility shift (Supplementary Figure 14g). These results indicate that EpiTrace can use ClockDML from different tissues of origin to predict single cell age, even in a cross-species scenario.

We then mapped human ClockDML to the zebrafish genome using a similar synteny-guided approach (Figure 2a) and tested EpiTrace prediction on a zebrafish scATAC dataset spanning from fertilization to the adult stage (GSE178969^50^) using this “human-guided” clock-like loci as reference. The mean EpiTrace age prediction from each stage closely approximated the known sample age (R = 0.97, Figure 2b). For each single cell type, the EpiTrace prediction closely assembles their time-of-emergence (Figure 2c). Similar results were obtained for “mouse-guided” clock-like loci (Supplementary Figure 13c, GSE152423^51^).

**Figure 2.**
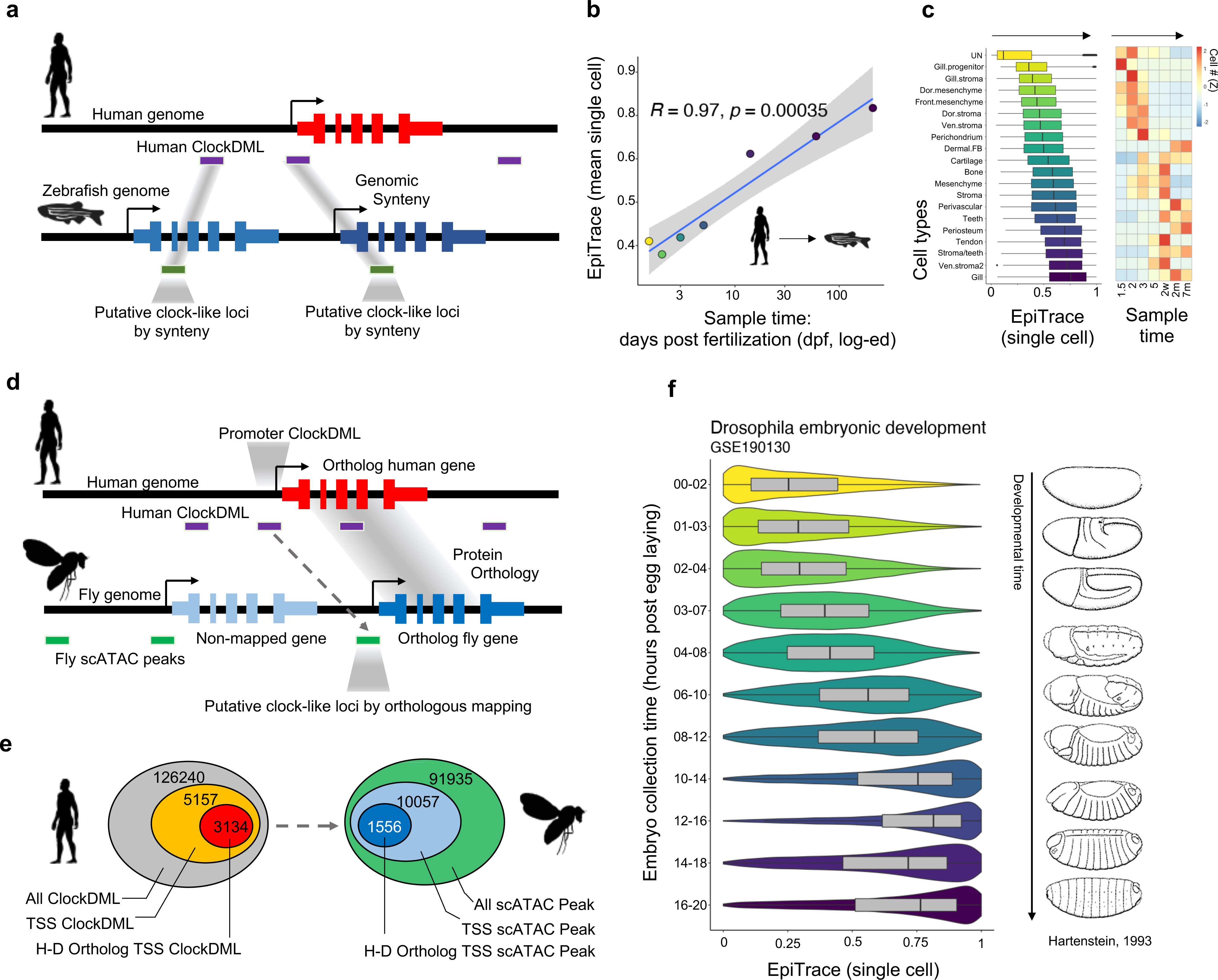
Mapping ClockDML orthologous genomic regions across species facilitates single cell age estimation using chromatin accessibility. **(a)** Schematic of the experiment. Human ClockDML are mapped to the zebrafish genome by homology to produce “human reference clock-like loci” and then used to infer zebrafish neural crest cell mitotic age. Since the data were provided as a one-hot matrix, we adopted the bulk-ATAC-like algorithm output. **(b)** Linear regression of predicted mean mitotic age (y) against days post fertilization (dpf) of the sample. **(c)** Single cell mitotic age of each cell type (left) and cell type-specific prevalence in samples of different ages (right). **(d)** Schematic of defining putative counterparts of human clock genomic loci in the *Drosophila* genome. Human ClockDML falling within +-100 bp of the gene transcription start site were defined as “Promoter ClockDML”. For human genes that simultaneously have a promoter ClockDML and one or more *Drosophila* ortholog gene(s), we define any *Drosophila* scATAC peaks falling within +-100 bp of transcription start sites of these *Drosophila* orthologs as putative clock-like genomic loci. These loci were subsequently used for *EpiTrace* analysis in the *Drosophila* dataset. **(e)** Diagram showing the number of ClockDML and scATAC falling in each category. **(f)** EpiTrace age of *Drosophila* embryonic development time series samples taken every 2 hours after egg laying (GSE190130^54^). Corresponding embryo sketches are shown on the right (adapted from Hartenstein, 1993).

For many animal species, active DNA methylation is not present in the genome; for example, the Drosophila melanogaster genome has <1% of CpG being methylated^52, 53^. We hypothesized that if clock-like chromatin accessibility is a universal phenomenon in ClockDML-homologous genomic regions, then identification of ClockDML-homologous genomic regions in such species might be sufficient for age prediction using EpiTrace.

Since these animal species are evolutionarily too distant from humans, only gene-level orthologous relationships could be reliably identified between their genomes and humans. To overcome this problem, we used an orthology-guided approach to first identify human-animal orthologous genes whose promoters encompassed ClockDML in the human genome and then identify the corresponding promoter genomic loci in the distant animal genome (Figure 2d). For the Drosophila melanogaster genome, we identified 1,556 such loci (Figure 2e). We then used this as “human-guided” clock-like loci in the Drosophila genome for reference in EpiTrace age prediction in a Drosophila embryonic development scATAC dataset. The prediction result showed high concordance with the known sampling time (Figure 2f, GSE190130^54^).

### ClockAcc change occurs upstream of the DNAm shift on ClockDML to report cell mitotic age

Notably, Drosophila is an invertebrate species that lacks canonical DNA methyltransferase and does not show CpG methylation. Only 0.4% (1hpf) - 0.1% (12hpf) of cytosine in the Drosophila genome was methylated^53^ compared to 6%-8% in humans, and most Drosophila methylated C was CpT/CpA. The fact that EpiTrace can work on the Drosophila genome suggests that clock-like chromatin accessibility might be independent of clock-like DNA methylation. To validate this model, we first tracked chromatin accessibility and DNAm on the same DNA molecule using the human embryonic development scCOOL-seq dataset (Supplementary Figure 15, GSE100272^55^) and a long-read nanoNOME dataset (Supplementary Figure 16, GSE183760^56^). In both datasets, chromatin accessibility shifts prior to ClockDML DNAm changes on the same molecule, indicating that clock-like DNA methylation is not necessary for clock-like chromatin accessibility.

We then performed forced transcriptional activation around ClockDML to test whether changes in chromatin accessibility would influence DNA methylation in these regions. We transfected sgRNA lentivirus targeting human G8 ClockDML group loci (shown in Figure 1b) into HEK293 cells stably expressing the dCas9-p300 transactivator (Supplementary Figure 17a). Gain of chromatin accessibility around these loci results in DNA hypomethylation on neighboring ClockDML (Supplementary Figure 17b-c), indicating that clock-like DNA methylation could be driven by chromatin accessibility shift.

In another conceptually similar scenario, we measured the DNAm change under forced transcription activation around mouse Sox1 loci and found them to linearly correlate with the age-dependent DNAm shift coefficient of corresponding loci in the human genome (Supplementary Figure 18, PRJNA490128^57^). Hence, changes of chromatin accessibility on ClockDML are sufficient to drive clock-like differential DNA methylation.

To validate whether changes in DNAm would affect chromatin accessibility on ClockDML, we tracked chromatin accessibility in a dataset where forced DNA methylation was performed with a ZNF-DNMT3A artificial methylator (Supplementary Figure 19a, GSE102395^58^). While ZNF-DNMT3A induction results in irreversible DNA methylation on ClockDML around its binding site (Supplementary Figure 19b, e), it does not change the overall chromatin accessibility on these loci (Supplementary Figure 19c-d, GSE102395^58^, GSE103590^59^), nor does it change the EpiTrace age on these cells (Supplementary Figure 19f-g, GSE102395^58^, GSE103590^59^).

Together, these data indicate that clock-like chromatin accessibility occurs upstream of the DNAm shift on ClockDML. In animals without active DNA methylation, genomic region exhibiting clock-like chromatin accessibility could also be identified. In other words, clock-like chromatin accessibility is an innate property on the clock-like loci, which usually harbor ClockDML. Furthermore, clock-like chromatin accessibility is independent from DNA methylation.

### EpiTrace tracks the reversal of cell age during iPSC induction

We tested EpiTrace on a single cell multiome (scMulti) sequencing dataset (CNP0001454^60^) of primed human embryonic stem cell (“Primed” hESC) cultures undergoing chemical reprogramming through a “4CL naïve PSC” state toward an 8-cell-like (“8CL”) state (Figure 3a), measured EpiTrace age in single cells, and compared the EpiTrace prediction with biological age predicted by whole-genome bisulfite sequencing (WGBS) of the same cultures^60^. Both DNAm-based prediction of sample age (Figure 3b) and single cell age predicted by EpiTrace (Figure 3c) suggest that mitotic age increases as cells undergo transformation, with single cell age gradually increasing across the evolutionary trajectory toward the 8CL state (Supplementary Figure 20). Furthermore, the biological age estimation of DNAm and EpiTrace were precisely correlated (correlation coefficient of mean single cell (sc) EpiTrace age x mean DNAm age: 0.998 (P = 0.04); scEpiTrace age x mean DNAm age: 0.526 (P = 1.9e-38)) (Figure 3d). While RNA velocity projections on these cells showed erroneous evolution trajectories rooted at single cells of a differentiated state (Supplementary Figure 21), combining RNA velocity and EpiTrace age of the same cell results in more biologically plausible evolution trajectories with the primed hESC as the root of all other cells (Figure 3e and Supplementary Figure 21). These results suggest that chromatin accessibility on ClockDML predicts single cell biological age at least as well as the DNAm-based age estimator, even in an age-reverse scenario.

**Figure 3.**
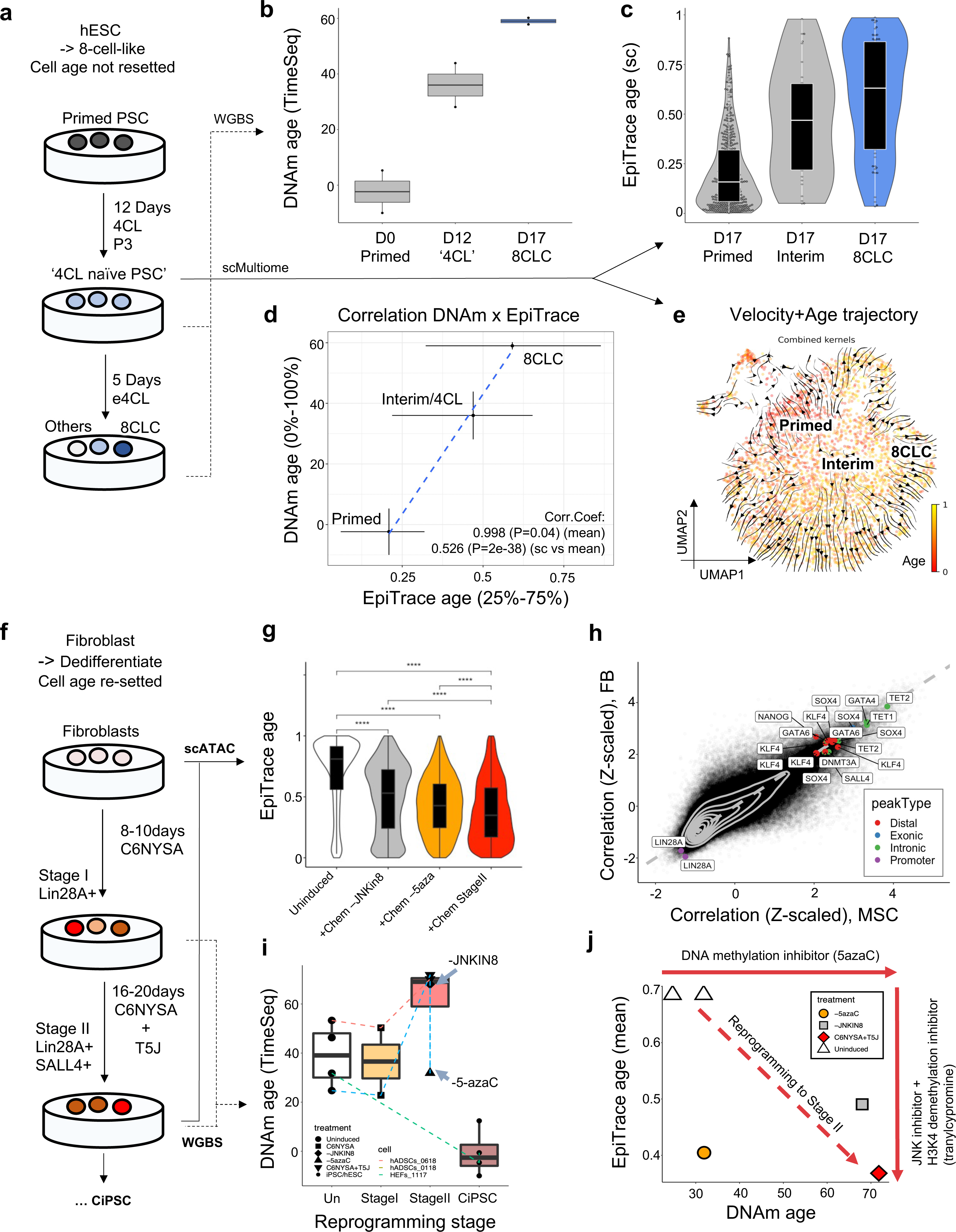
Inferring single cell age reversal in iPSC induction with EpiTrace. **(a)** Schematic overview of the *in vitro* chemical induction of human pluripotent stem cells (“Primed”) back to naïve (“4CL”) or 8-cell-like (“8CL”) states. **(b)** DNAm age of D0 (Primed) and D12 (4CL) cultures and sorted 8-cell-like cells (8CLC) from D17 culture, from WGBS data. **(c)** Single cell age estimated with EpiTrace from the D17 scMultiome dataset. **(d)** Correlation of inferred age from DNAm (min-mean-max, y-axis) or single cell EpiTrace age (25%-50%-75%, x-axis) from the same set of cells. **(e)** UMAP projection of scMultiome-sequenced D17 culture with single cell evolution trajectories built with kernels combining EpiTrace age and RNA velocity information. **(f)** Schematic overview of the *in vitro* chemical induction of adult fibroblasts toward human pluripotent stem cells. Both the uninduced and intermediate stage II cultures are sequenced by scATAC. **(g)** Single cell age estimated with EpiTrace from **(f**). The induced cultures were either subjected to the full induction paradigm (+Chem: C6NYSA+T5J) or with 5-azaC/JNKin8 removed (−5aza, −JNKin8). Statistical comparisons are shown between groups (***: P<0.001; Wilcox test). **(h)** Correlation coefficient between the chromatin accessibility on each ATAC peaks and EpiTrace age estimated from the MSC experiment (x-axis) or FB experiment (y-axis). Peaks of interest were labeled, colored by their genomic location class. **(i)** Prediction of sample age by DNAm from WGBS data of chemical induction of iPSCs. Chemical reprogramming induces genome-wide demethylation and an increase in DNAm age, as reported previously^31^, while the addition of 5-azaC globally reduces DNA methylation to increase DNAm age. Removal of 5-azaC blocks DNAm age from increasing. **(j)** Scatter plot of WGBS DNAm age (x-axis) and mean single cell EpiTrace age (y-axis) of the same sample. Abbreviations: HEF: human embryonic fibroblast; hADSC: human adipose stromal cell (mesenchymal stromal cell); C6NYSA: combination of CHIR99021, 616452, TTNPB, Y27632, SAG, and ABT869; JNKIN8: c-Jun N-terminal kinase inhibitor; 5-azaC: 5-azacytidine; T5J: tranylcypromine, JNKIN8 and 5-azaC.

We measured EpiTrace age in an additional scATAC dataset of cells undergoing the early stages of chemical reprogramming from differentiated endodermal cells (fibroblast (FB), or mesenchymal stem cell (MSC)) toward induced pluripotent stem cells (Figure 3f, GSE178324^61^) and compared them with WGBS-predicted ages from the same study. EpiTrace age prediction of single cells significantly decreased at stage II compared to the uninduced state (Figure 3g), indicating that these cells are “rejuvenated” as expected. Compared to uninduced cells, the EpiTrace age of C6NYSA+T5J cells are significantly rejuvenated (decreased). Removal of 5-azaC from the treatment resulted slightly impairs the rejuvenation. On the contrary, removal of the JNK inhibitor from the treatment resulted in more significant impairment of rejuvenation (Figure 3h). The age-peak association from the MSC reprogramming experiment is highly similar to that from the FB reprogramming experiment. (Figure 3g). These results suggest relevance between the observed mitotic age resetting and cell fate reprogramming.

The DNAm-predicted age of cells during the chemical induction procedure shows that the biological age first increases at induction stage II (Figure 3i, GSE178966^61^) before decreasing to near zero in the pluripotent state. Removal of 5-azaC from the induction formula blocks the age increase at stage II, indicating that the apparent DNAm age increase is a result of global DNA demethylation^31^. Similar to our previous observations, comparing the DNAm age and EpiTrace age prediction of the same cell sets suggests that chromatin accessibility on ClockDML is independent from ClockDML DNAm change (Figure 3j).

### EpiTrace in combination with genetic tracing reveals the molecular determinant of Hayflick’s limit at the single cell level

To test EpiTrace age estimation in genetically defined cell lineages, we took advantage of a mitochondria-enhanced scATAC dataset (GSE142745^8^) of cultured CD34 hematopoietic stem cells (HSCs) that underwent *in vitro* expansion for 14 days before being forced into differentiation under LIF/EPO toward myeloid/erythroid lineages for an additional 6 days (Figure 4a). These cells were sequenced at day 8 (D8), 14 (D14), and 20 (D20). Cells were clustered by their transcriptomic (scRNA) phenotype as progenitors (Prog), differentiated (Diff) or terminally differentiated (Terminal) cells, which gradually emerged over days in culture (Figure 4b and Supplementary Figure 22). Furthermore, they were segregated into lineages (clones) arising from the same progenitor by mitochondrial SNV.

**Figure 4.**
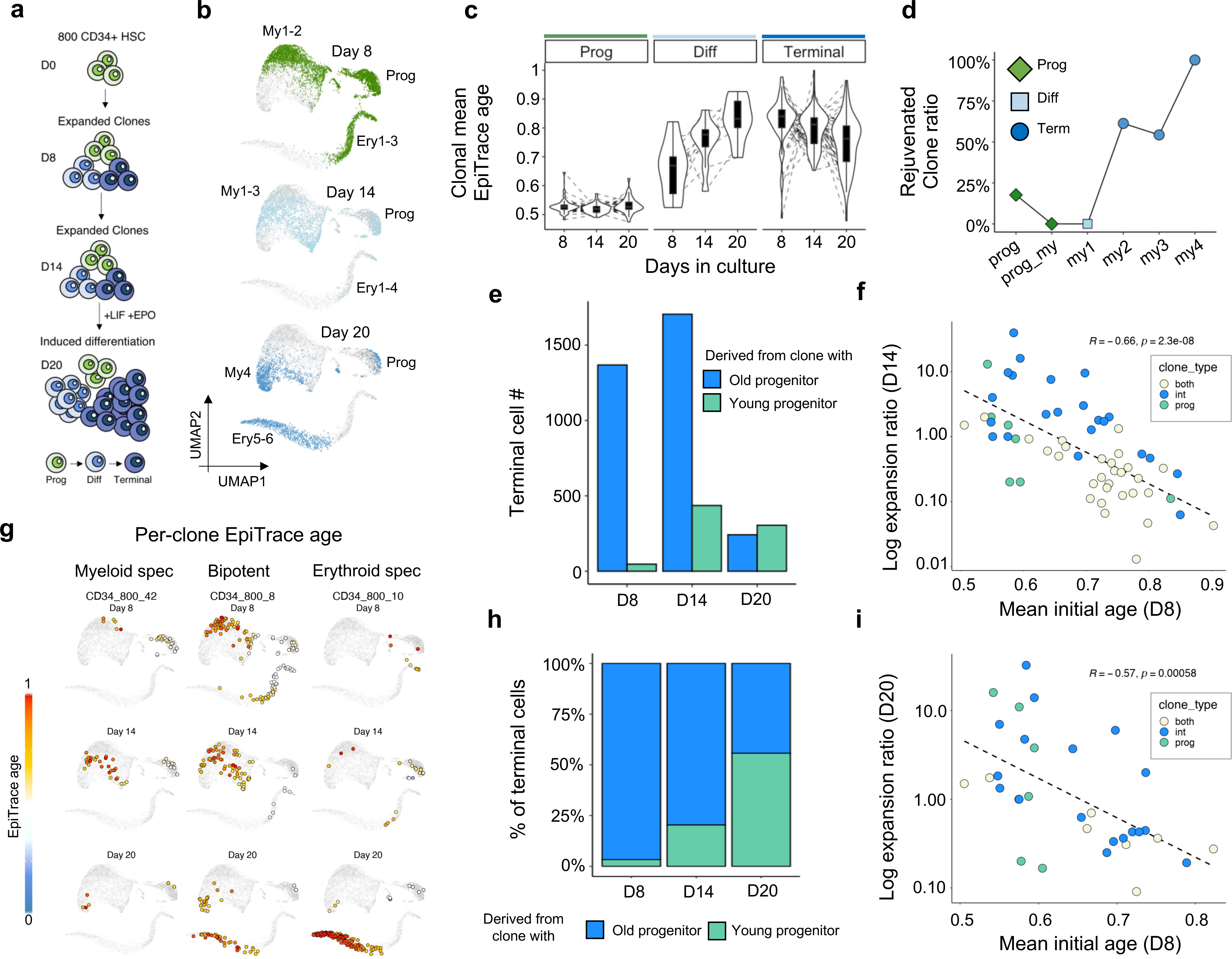
Single cell age estimation revealed that epigenomic age determines clonal expansion potential. **(a)** Schematic of the experiment. CD34+ hematopoietic stem cells were used in the *in vitro* expansion/differentiation experiment. Cells were first expanded to day 8 (D8) (CD34_500) or day 14 (D14) (CD34_800) and then differentiated by LIF and EPO until day 20 (D20). Mitochondrial mutations from the scATAC experiment were used for tracking cells derived from similar clones. Cell phenotypes were determined by the scATAC profile. **(b)** Cells from experiments performed on D8, D14, and D20, showing a gradual transition toward terminally differentiated myeloid (my4) and erythroid (ery6) cells. **(c)** Tracking the mean EpiTrace age of each myeloid cell clone at each timepoint. **(d)** Ratio of rejuvenated (clone age decrease over time) clones in all clones for the myeloid cells. The terminally differentiated cells are dominated by rejuvenated clones. **(e)** Number of terminal myeloid cells derived from young proliferator clones (mean initial clonal EpiTrace age < 0.7) and old proliferator clones (mean initial clonal EpiTrace age >= 0.7) at three time points. **(f)** Scatter plot of the log-clonal expansion ratio on day 14 (Y-axis) compared to the mean initial clonal EpiTrace age of the same clone (X-axis). Clonal types are color-labeled. **(g)** EpiTrace age (color) of the single cells derived from a similar clone. Three clones with different fates are shown for example. The CD34_800_42 clone was a myeloid-specific clone that generated only myeloid cells. The CD34_800_8 clone was a bipotent clone that generated both myeloid and erythroid decedents. The CD34_800_10 clone is an erythroid-specific clone that generates predominantly erythroid cells. **(h)** Relative contribution of young proliferator clones (mean initial clonal EpiTrace < 0.7) and old proliferator clones (mean initial clonal EpiTrace age >= 0.7) in the terminal myeloid cell population at three time points. **(i)** Scatter plot of the log-clonal expansion ratio on D20 (Y-axis) compared to the mean initial clonal EpiTrace age of the same clone (X-axis). Clonal types are color-labeled. Correlation statistics (R and P-value): Pearson’s. Group statistics: t-test.

We used EpiTrace to predict mitosis age in these cells, separately for myeloid lineage and erythroid lineage cells. Since the single cell age prediction by EpiTrace could be affected by highly biased cell composition (Supplementary Figure 23), we selected a relatively balanced CD34_800 dataset for erythroid lineage cell age prediction. Both CD34_500 and CD34_800 datasets were used for myeloid lineage cell age prediction.

The age prediction shows high concordance with known sampling days across cell types and enables tracking the mitosis age of individual cells derived from the same clone (Figure 4g). For the myeloid lineage, the EpiTrace age of cells from the same clone increased from progenitor to terminal myeloid cells (Figure 4c-d). Forced differentiation increased the age of differentiated cells as expected but decreased the age of terminal cells (Figure 4c-d). To explore this phenomenon in depth, we classified clones according to the relative age change between days of culture (Supplementary Figure 24): clones that exhibited an age increase from D8 to D14 in a cluster as “Aged”, and those that exhibited an age decrease from D8 to D14 as “Rejuvenated”. While most progenitor and differentiated cells show clonal aging during induction, clonal rejuvenation dominates the terminally differentiated clusters (Figure 4c-d). The proportion of clones showing a rejuvenation increase in terminal cells (Figure 4d), in correlation with their differentiation state, suggests that these terminally differentiated cells were derived from younger hematopoietic progenitors instead of existing intermediate differentiated cells.

To validate this hypothesis, we analyzed the expansion capability of different cell clones, which processes different types of proliferating cells, including progenitor (Prog) and intermediate differentiated cells (Int), from the CD34_800 experiment (which was sequenced on all three timepoints). We first classified the cell clones according to their cell composition on D8 (the first time point): at this time, clones with only Prog but no Int cells were classified as “Prog-only”; clones with only Int but no Prog cells were classified as “Int-only”; and clones with both Int and Prog cells were classified as “Both” (Supplementary Figure 25a). The mean EpiTrace age of the clones at the initial timepoint was measured as mean EpiTrace age of cells from D8 (Supplementary Figure 25b, d). We then tracked their clonal derivatives at the next timepoints (D14 and D20) to see if a clone was expanded (defined as an increased terminally differentiated cell number at later timepoints compared to D8). For each expanded clone, we calculated the clonal expansion ratio, defined as the increased number of terminally differentiated cells divided by the total cell number on D8.

At both D14 and D20, the log clonal expansion ratio was inversely correlated with the initial EpiTrace age of the clone (Figure 4f, i, and Supplementary Figure 25c, e): the correlation between the log clonal expansion ratio and initial clonal age was R = −0.66 (P = 2.3e-08) for D14 and R = −0.57 (P = 0.00058) for D20. While Int-only clones expanded better than Prog-only clones at earlier timepoints (Supplementary Figure 25c), the Prog-only clones caught up at the latter timepoint and showed improved expansion potential (Supplementary Figure 25e).

We then re-classified the clones by their mean clonal age at D8 into “young progenitor derived clones” (defined as the mean EpiTrace age < 0.7), or “old progenitor derived clones” (defined as the mean EpiTrace age >= 0.7). The number of terminal cells derived from young clones steadily increased during the stimulation time course, outnumbering the terminal cells derived from old clones on day 20 (Figure 4e). As a result, the relative contribution of terminal cells from young clones steadily increased during the stimulation time course (Figure 4h and Supplementary Figure 25f-g), explaining the observed decrease in terminal cell EpiTrace age (Figure 4c-d).

Combining the observations, we conclude that the clonal expansion potential is better explained by clonal epigenetic age instead of the initial phenotype of proliferating cells in the clone. Interestingly, the initial clonal age in clones with both Prog and Int cells was significantly older than that in Prog-only or Int-only clones (Supplementary Figure 25b, d). These clones expanded the least at both timepoints (Figure 4f, i and Supplementary Figure 25c, e). This result indicates that cells in these clones, though phenotypically classified as capable of proliferation, are at the end of their expansion potential.

Together, these results supported the model that during *in vitro* HSC-stimulated expansion, terminally differentiated cells are preferentially derived from younger progenitors. In other words, younger hematopoietic progenitor cells are much more capable of expansion and differentiation. In the seminal paper Hayflick himself determined the *in vitro* passage limit of cultured cell^62^, he cocultured 46,XX and 46,XY cells with different *in vitro* passage number together. By counting the karyotypes of cells in the final passage population, he found that the “younger” cells with less starting passage number always dominated the final passage population. Our current experiment is by design similar to Hayflick’s original experiment by using a genetic marker, mitochondrial mutation, to track each clone. By measuring the “clonal age” of these single cells, EpiTrace derived a quantitative measure of future expansion potential against the current age of the clone. A pioneering study showed that the genome-wide DNA methylation level decreases during cell culture passage^63^. This phenomenon was later used to propose a method to infer Hayflick’s limit for individual cell lines^64^. This result now provided the first experimental evidence for the pioneering theoretical works.

### Single cell age estimation facilitates discovery of molecular markers of T cells underlying the anti-PD1 response

The CD34 dataset demonstrated above is based on an ideal *in vitro* scenario with cells at a similar starting point and only proliferate and die without exchange with the external environment. To test EpiTrace in a more complex cell population in an *in vivo* setting, with possible influx, efflux and proliferation, we applied EpiTrace to a scATAC dataset comprising biopsies from basal cell carcinoma pre- and post-anti-PD1 treatment (Figure 5a, GSE129785^65^). Post-anti-PD1 treatment, cytotoxic T cells with exhaustion markers (Tex) are significantly increased in anti-PD1 responders (R) but not nonresponders (NR). More immature Tex cells were present in nonresponders, and this phenomenon was exaggerated after anti-PD1 treatment. However, overall maturity did not change in responders. Importantly, the EpiTrace age of interim and mature Texs in responders did not change after the anti-PD1 treatment, suggesting that the increased cell number might not be solely due to local proliferation of pre-anti-PD1 mature Texs (Figure 5b).

**Figure 5.**
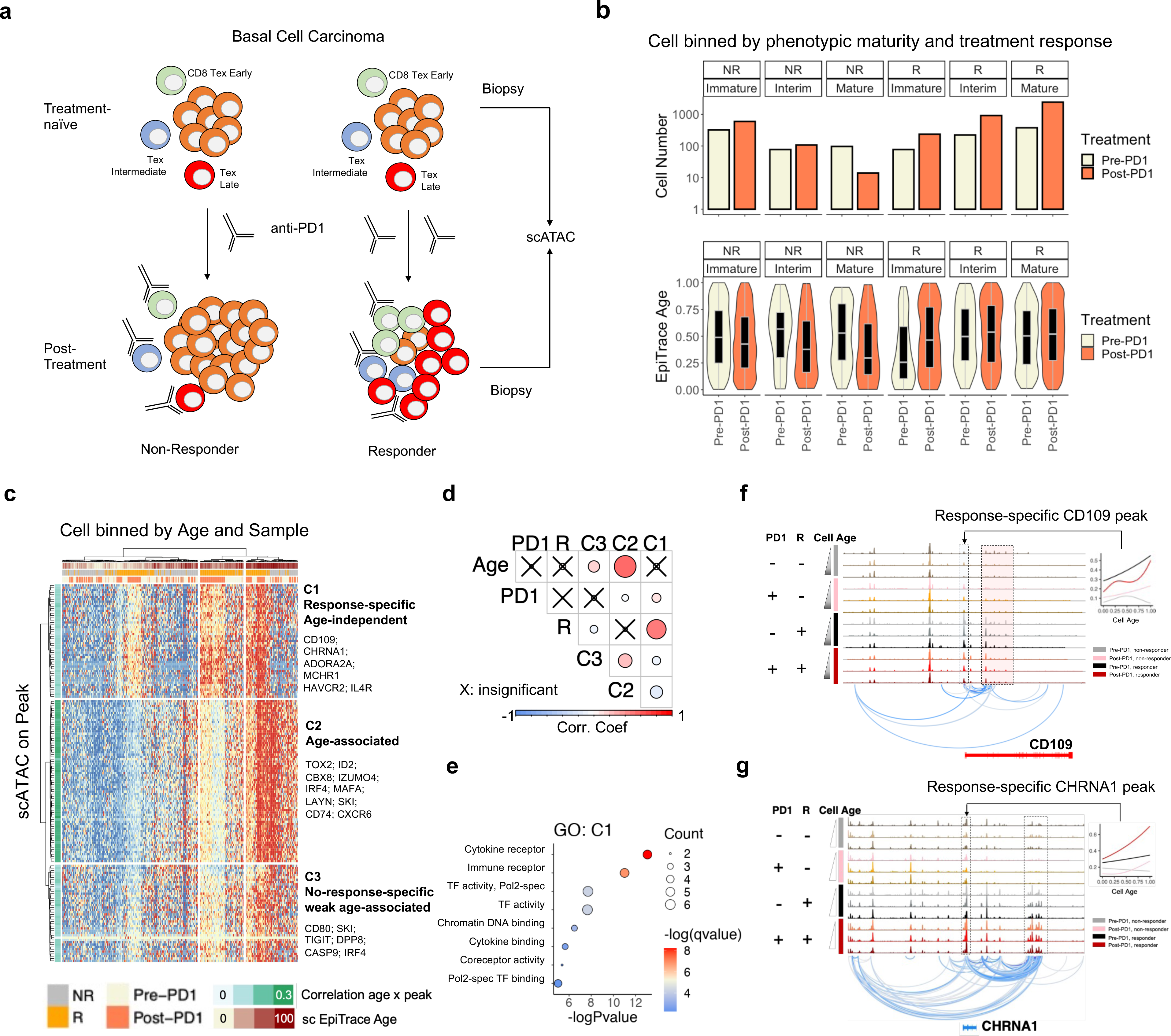
Single cell age estimation facilitates the discovery of molecular markers of peripheral-influx T cells underlying the anti-PD1 response. **(a)** Schematic overview of the experiment. Biopsies were taken from basal cell carcinoma patients before (pre) and after (post) anti-PD1 treatment and subjected to scATAC sequencing. **(b)** Cell number (above) and EpiTrace age of “exhausted T” (Tex) cells, separated by treatment response (R: responder; NR: non-responder) and phenotypic maturity (Immature/Interim/Mature). **(c)** Heatmap showing scATAC peak activity in pseudobulk single cells grouped by phenotype (R/NR), sample (pre- or post-anti-PD1), and EpiTrace age. Correlations between peak activity and EpiTrace age are shown on the left. Peaks were clustered according to their activity profile into response specific, nonresponse specific, and age-associated clusters. **(d)** Correlation coefficient between clusters of peaks and treatment (PD1: pre/post = 0/1), response (R: NR/R = 0/1), and cell age (Age). Nonsignificant correlations are labeled with “X”. **(e)** GO enrichment of the C1 cluster peaks as in (c). **(f)** ChrAcc (top) and cross-correlation between peaks (bottom) of CD109 loci from pseudobulk single cells grouped with phenotype (R/NR), sample (pre- or post-anti-PD1), and EpiTrace age (Young/Interim/Mature). The association of the CD109 promoter ChrAcc across age is shown in the right panel. **(g)** ChrAcc (top) and cross-correlation between peaks (bottom) of CHRNA1 loci from pseudobulk single cells grouped as in (f). The association of the CHRNA1 promoter ChrAcc across age is shown in the right panel.

New post-anti-PD1 mature Tex cells could be either derived from pre-anti-PD1 immature Tex cells or from the influx of peripheral T cells. To test these alternatives, we performed a correlation of chromatin accessibility on Tex differentially expressed peaks and cell age. Hierarchical clustering of the peak openness of pseudobulk cells from similar age and phenotype segregated peaks into three clusters: C1: response specific, age-independent; C2: response-irrelevant and age-associated; and C3: nonresponse-specific peaks that were weakly associated with age (Figure 5c-d). Interestingly, known markers of activated (TIGIT, LAYN, HAVCR2) and tumor-reactive T cells (ENTPD1) were segregated into different clusters. T-cell markers (TOX2, ID2, MAFA) and tissue resident Trm marker (CXCR6) belong to the group of scATAC peaks that are mainly associated with age but are not associated with PD1 response (Figure 5c-d). The anti-PD1 response is not associated with cell age but instead with C1 peak expression. In contrast, cell age is associated with C2/C3 peaks, which are related to no response under anti-PD1. GO enrichment of this C1 cluster, in contrast to C2/C3 cluster genes, showed particular enrichment in the “cytokine receptor” and “immune receptor” pathways (Figure 5e and Supplementary Figure 26), highlighting genes such as IL4R, CD74, IFNGR2 and IFNAR2 which might be implicated in the anti-PD1 response. Finally, we identified that cis-regulatory loci of a coreceptor and negative regulator of TGF-beta, CD109 (Figure 5f), and the nicotinic acetylcholine receptor CHRNA1 (Figure 5g), are specifically activated in response-associated Tex cells, suggesting novel targets for future research.

### EpiTrace reveals the developmental history during human cortical gyrification

To test how mitotic age estimation might complement RNA-based development analysis, we applied EpiTrace to a single cell 10x Genomics multiomic sequencing (scMultiomic) dataset from the pcw21 human fetal brain cortex (GSE162170^66^) to study the trajectory of glutaminergic neuron development (Figure 6a). Glutaminergic neurons develop from radial glia (RG) through the cycling progenitor (Cyc. Prog) cells into neuronal intermediate progenitor cells (nIPC), before it undergoes a cascade of maturation (GluN1->2->3->4->5)^66–68^. We modeled the cell fate transition by CellRank^69^ with kernels built with RNA velocity, CytoTrace, EpiTrace age, or a combined kernel with all three estimators. While RNA velocity and CytoTrace produced inconsistent transition trajectories that pointed toward a group of nIPCs (Figure 6b, i-ii), kernels with EpiTrace age revealed a correct direction of development from the nIPCs toward terminally differentiated neurons (Figure 6b, iii). The combined kernel of all three estimators resulted in a biologically plausible transition trajectory that starts from RG to bifurcate into two different branches, each giving rise to a distinct nIPC population that differentiates into mature neurons (Figure 6a).

**Figure 6.**
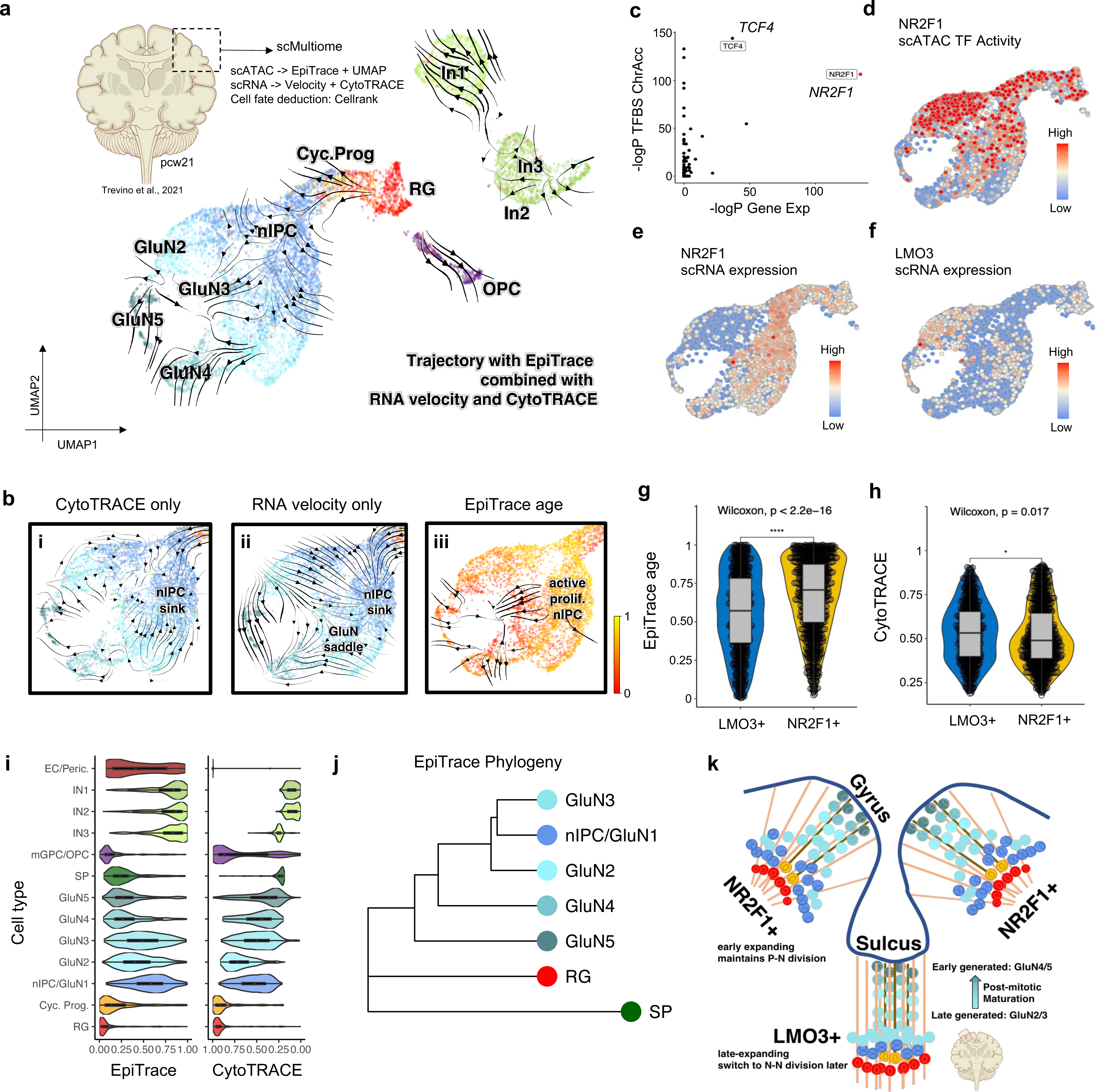
EpiTrace reveals the developmental history during human cortical gyrification. **(a)** UMAP projected cell evolution trajectory built with cellrank by using a hybrid kernel of EpiTrace, CytoTRACE, and RNA velocity of a scMultiome-seq dataset from a pcw21 human brain. Abbreviations: RG: radial glia; Cyc. Prog: cycling progenitor; nIPC: neuronal intermediate progenitor cell; GluN: excitatory glutaminergic neuron; IN: inhibitory GABAergic neuron; EC: endothelial cell; mGPC/OPC: medial ganglionic eminence progenitor/oligodendrocyte precursor cell; SP: subplate neuron. SP and EC are not shown in the figure due to space limitations. **(b)** Trajectories built with only CytoTRACE **(i)** or RNA velocity **(ii)** resulted in unrealistic “sinks” and “saddles” on the map. In contrast, EpiTrace age **(iii)** provided a unidirectional reference of time to reveal that the “sink” nIPC population is mitotically active to resolve the “nIPC stall”. **(c)** Scatter plot of the differential gene expression estimate (−log P-value, x-axis) and differential TFBS-specific ChrAcc estimate (log P-value, y-axis) in the GluN cells. Most significantly differential expressed transcription factors NR2F1 and TCF4 were highlighted in the figure. **(d)** UMAP projection of TFBS-specific ChrAcc of NR2F1. **(e)** Expression of NR2F1 on UMAP. **(f)** Expression of LMO3 on UMAP. **(g)** EpiTrace age of cells belong to the LMO3+ population or NR2F1+ population. **(h)** CytoTRACE of cells belong to the LMO3+ population or NR2F1+ population. **(i)** Mitotic clock (EpiTrace) and differentiation potential (CytoTRACE) of the same cell in scMultiome-seq. The CytoTRACE score is reversed to show differentiation from left to right to facilitate comparison with EpiTrace. **(j)** Excitatory neuron phylogeny built with mitotic clock, showing that GluN4/GluN5 are likely direct, early-born progenies of RG, while GluN2/GluN3 are likely late-born, immature progenies of nIPC. **(k)** Overall model of corticogenesis in the light of EpiTrace. Data source: Trevino *et al.* 2021^66^.

Two transcription factors, TCF4 and NR2F1 (COUP-TFI), were differentially expressed between the branches. They exhibit significant differential binding activities in these neurons (Figure 6c). Interestingly, NR2F1 is mainly expressed in the gyrus of the human cortex, and hereditary NR2F1 loss-of-function mutations are associated with mental retardation and the polymicrogyri phenotype^70–72^. NR2F1 TFBS-associated peaks are open in a branch (Figure 6d) that is NR2F1 negative (Figure 6e) and LMO3 positive (Figure 6f), suggesting that NR2F1 turned into a transcriptional repressor in nIPC. The EpiTrace age of the NR2F1^+^ branch nIPC was significantly higher than that of the LMO3+ nIPC, suggesting accelerated mitotic activity (Figure 6g). In concordance with this, the CytoTRACE score of LMO3^+^ nIPC was higher than that of NR2F1^+^ nIPC (Figure 6h). These results indicate that nIPCs are divided into NR2F1^+^ clones that support earlier neurogenesis and LMO3^+^/NR2F1^-^ clones that expand relatively later, linking the gyrus-specific expression pattern of NR2F1 to its function in cortical gyrification^70^.

We compared the EpiTrace age of the neurons with their CytoTRACE score (Figure 6i). While the CytoTRACE score of glutaminergic neurons correlates with their differentiation, the EpiTrace age of these cells is inversely correlated with their maturity. To explain this inconsistency, we built a “phylogenetic tree” with ClockDML chromatin accessibility (EpiTrace phylogeny). We reasoned that cells traverse on the phenotype manifold on branched trajectories while they undergo mitosis. As they evolve, chromatin accessibility on ClockDML converges into a specific state that should be lineage-dependent because of the irreversible nature of such change. Hence, it is possible to infer cell lineage trees using phylogenetic-like methods. Such analysis revealed a birth sequence of glutaminergic neurons: GluN5 is first divided from RG, followed by GluN4, GluN2, GluN3 and GluN1/nIPC (Figure 6j), indicating that neurons that formed earlier undergo longer postmitotic maturation (Supplementary Figure 27). In concordance with this observation, by analyzing scRNA expression of the same cells, we found that while the late-aged nIPC/GluN1 and GluN2 cells still showed reminiscent RNA expression of the proliferating cells, such as SOX11, SOX4, MALAT1 and NFIB, the earlier-aged, “more mature” GluN5 and GluN4 cells showed significantly increased expression of mature neuron markers, including synaptic proteins, including SYT4, FABP7, APP, GAP43, SYT11, and PCDH17, mature neuron cytoskeleton proteins, such as TUBB2A and NEFL, and post-mitotic functioning transcription factors, such as MEF2C (Supplementary Figure 28). Furthermore, in concordance with the known “inside-out” developmental paradigm of the cortex^73^, the earlier-aged GluN5 specifically expresses the layer V/VI marker genes SCUBE1 and SEMA3E^74^, while the younger GluN4 population expresses comparably higher levels of the layer III/IV marker genes NTNG1 and MME^74^ (Supplementary Figure 29). Hence, the dynamics of postmitotic neurons undergoing continuous differentiation could be captured by combining mitotic age with other modality measurements.

Together, this analysis demonstrated that EpiTrace age analysis complements RNA velocity and stemness prediction in characterizing complex organ development, indicated a long post-mitotic maturation of neurons, and revealed the molecular mechanism of NR2F1 controlling human nIPC proliferation to underlie cortical gyrification (Figure 6k).

### Single cell lineage tracing by mitotic age loci indicates dedifferentiation driven by master transcription factors in translocation renal cell carcinoma

We have already demonstrated that EpiTrace can track development using developing tissues. To test whether EpiTrace can recover epigenomic changes during development from a single, terminally developed, static snapshot from adult tissue, we applied EpiTrace to a scATAC dataset from adult human kidney (Figure 7a, GSE166547^75^). The birth sequence of kidney cells by EpiTrace phylogeny analysis suggests an endothelial origin of kidney tubules and delineates a cell type-specific generation cascade during nephrogenesis (Figure 7b), with correlation to their spatial position (Supplementary Figure 30). The distribution of EpiTrace age for each cell type suggests a distal-to-proximal genesis cascade of nephron tubules with a late expansion of proximal tubules (Figure 7c).

**Figure 7.**
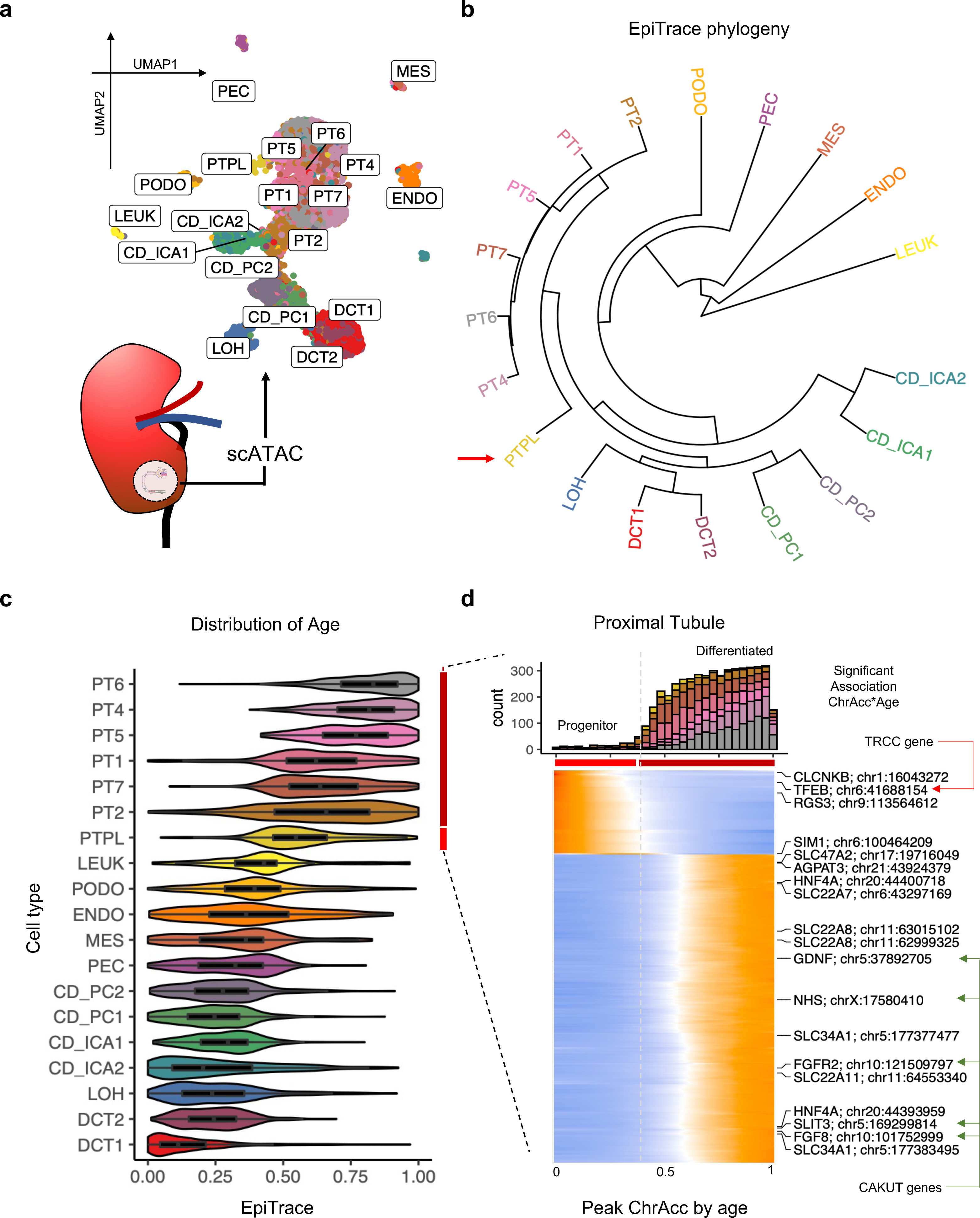
Single cell phylogeny on mitotic age loci indicates dedifferentiation driven by master transcription factors in translocation renal cell carcinoma. **(a)** UMAP projection of scATAC data of normal human kidney samples from biopsies. Abbreviations: LEUK: leukocytes; ENDO: endothelial cells; MES: mesangial cells; PEC: parietal epithelial cells; PODO: podocytes; PT: proximal tubule; PTPL: proximal tubule progenitor-like cell; LOH: loop-of-Henri; DCT: distal collection tubule; CD_PC: collection duct principal cell; CD_ICA: collection duct intercalating cell. **(b)** Single cell phylogeny built with chromatin accessibility on ClockDML (EpiTrace phylogeny). **(c)** Distribution of EpiTrace age for each cell type. **(d)** Cell age distribution of PT cells (colored by type) (top) and ATAC peak activity across cell age (bottom) in PT cells. Genes of interest are labelled on the right. Known translocation renal cell carcinoma (TRCC) driver gene: TFEB; known hereditary renal dysgenesis (CAKUT) genes: FGF8, FGFR2, SLIT3, GDNF and NHS.

In the proximal tubule (PT) lineage, EpiTrace age-derived phylogeny could be orthogonally validated with snRNA-derived phylogeny (Supplementary Figure 31, GSE121862^76^). The correlation between EpiTrace age and peak openness showed clear segregation of peaks opened in progenitor or differentiated proximal tubule (PT) cells (Figure 7d). Importantly, such association is not guided by known cell type information, indicating the power of EpiTrace in positioning single cells along their evolutionary trajectory. Interestingly, the translocation renal cell carcinoma (TRCC) driver gene TFEB is specifically activated in progenitor cells and shows an age-dependent decrease in activity. In contrast, all hereditary renal dysgenesis (CAKUT) genes FGF8, FGFR2, SLIT3, GDNF and NHS are associated with differentiated cell specific, age-dependent increased peaks. These results suggest that CAKUT is linked to genes functioning in terminal proximal tubule cell fate determination and function, while TRCC oncogenesis is linked to the misexpression of progenitor-specific transcription factors, possibly forcing the dedifferentiation of terminally differentiated proximal tubule cells into a stem-like state.

### Early branching evolution of glioblastoma clonal evolution tracked by single cell mitotic age

Finally, we analyzed an individual tumor sample (CGY2349) in a human glioblastoma scATAC dataset to study whether EpiTrace age analysis could work for cell evolution in oncogenesis (Figure 8a, GSE139136; GSE163655; GSE163656 ^77^). In this tumor, copy number variation (CNV) analysis showed that MDM4 amplification dominates the malignant clones, which additionally have either EGFR or PDGFRA amplification, resulting in increased chromatin accessibility around these genes (Figure 8b-e). With EpiTrace, we identified a pre-malignant cluster (7) that is younger than all malignant clones (4/6/5/0/3) but shows accelerated aging/mitosis count compared to the “normal clones” (1/9) (Figure 8b, f), has lower MDM4 amplification (Figure 8c), and is without either EGFR or PDGFRA amplification (Figure 8d-e).

**Figure 8.**
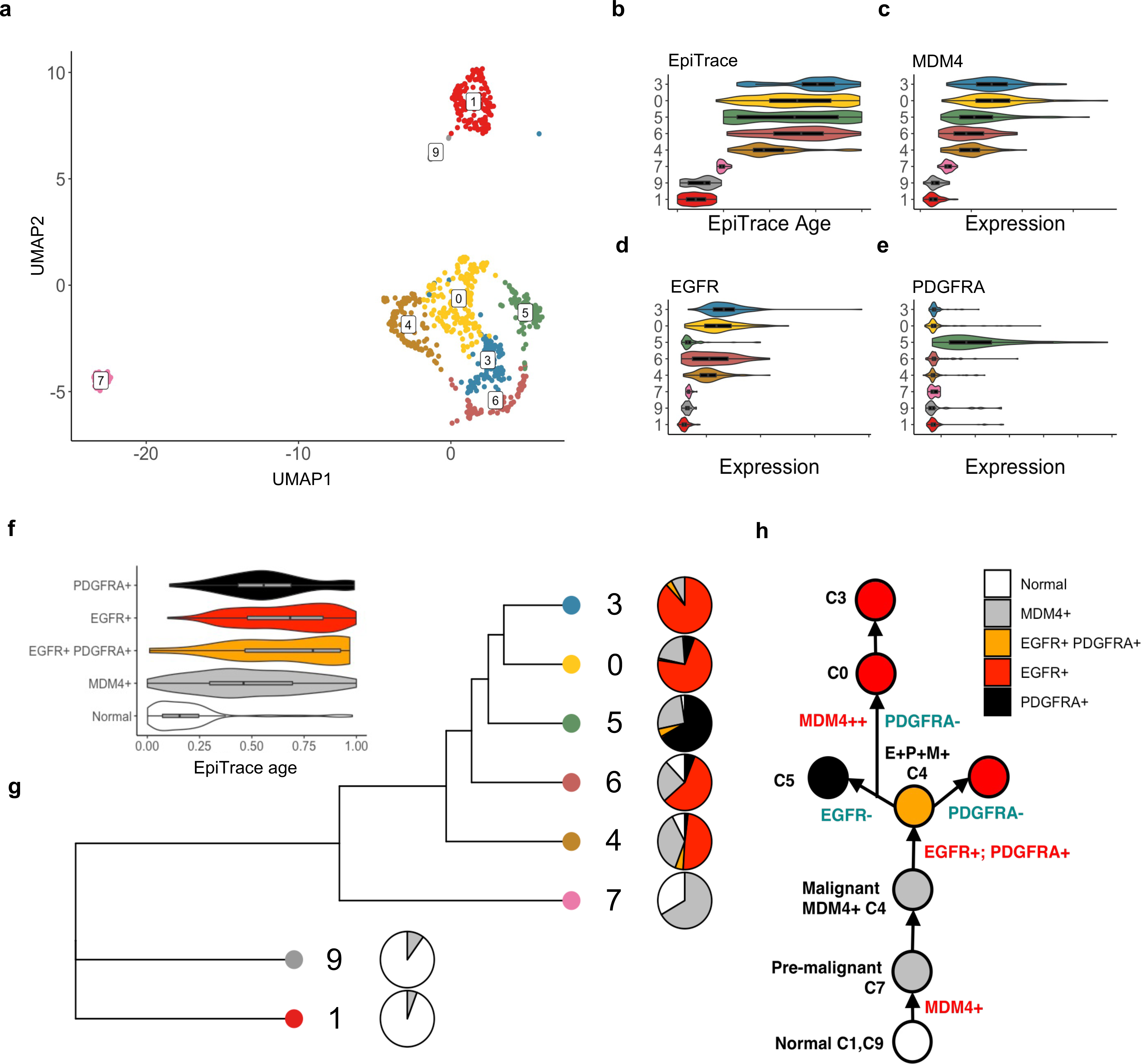
Early-on branching evolution of glioblastoma clonal evolution tracked by single cell mitotic age. **(a)** UMAP of scATAC data from GBM sample CGY4349. Cell clusters are labeled (Numbers). Per-cell-cluster EpiTrace mitotic clock age **(b)** and MDM4 **(c)**, EGFR **(d)**, PDGFRA **(e)** promoter ChrAcc. Note that MDM4 amplification is further gained in younger clones 0/3. **(f)** EpiTrace age of cells with normal karyotype, MDM4+, MDM4+/EGFR+/PDGFRA+, MDM4+/EGFR+, and MDM4+/PDGFRA+. **(g)** EpiTrace phylogeny, clonal composition (pie charts on the right), showing that the original pre-malignant cell cluster (7) evolved into an initial malignant clonal expansion with MDM4 amplification, followed by a malignant MDM4+ population that was double-positive with PDGFRA and EGFR amplifications. In later stages, tumor clones split the oncogenic amplification and grow into either PDGFRA+ (5) or EGFR+ (6/0/3) populations. **(h)** Schematic evolution trajectory.

Interestingly, some MDM4+ cells had both EGFR and PDGFRA amplification (Supplementary Figure 32). EpiTrace age analysis revealed that the MDM4^+^ only cells are ancestral to triple-positive, EGFR^+^/PDGFRA^+^ cells, followed by loss of either EGFR or PDGFRA in the progeny (Figure 8f). This is further supported by EpiTrace phylogeny analysis (Figure 8g). Branched evolution of MDM4^+^/EGFR^+^ and MDM4^+^/PDGFRA^+^ cells was initiated at the beginning of malignant transformation (Supplementary Figure 32). Together, these results characterized the evolutionary trajectory of malignancy from the MDM4^+^ pre-malignant clone to the earliest malignant cell population with amplification of MDM4, PDGFRA and EGFR in a catastrophic genomic instability event, which bifurcated into heterogeneous clones with either PDGFRA or EGFR addiction (Figure 8h). EpiTrace age analysis revealed the pre-malignant state of this tumor and suggested branching evolution of this tumor to indicate that heterogeneous cancer clones arise early in malignancy transformation.

It was previously known that telomere crisis and mitotic missegregation can cause catastrophic events in a single mitosis, most importantly chromothripsis^78, 79^, chromoplexy^80^, and kataegis^81^. Multiple structural variations over the genome can occur simultaneously during such events, resulting in a synchronous, punctuated burst of chromosomal copy number aberration^78, 82^. Most importantly, by timing the occurrence time of these mutational events, it was identified that such events occur early during oncogenesis^83, 84^. PDGFRA and EGFR amplifications were reported to exist in different single cell clones that coexist in a mosaic manner in GBM tumors^85^. While most reports suggest that these mutations are mutually exclusive in single GBM-derived cell lines or tumor sphere cultures^86^, these clones coexist within the same tumor and share common somatic mutations, such as deletion of PTEN and CDKN2A^85, 87^, indicating that they were derived from the same ancestral clone. scRNA-seq^88, 89^ suggests that PDGFRA+/EGFR+ double-positive cells exist in GBM. Single-positive, PDGFRA+ or EGFR+ descendent clones could emerge from double-positive parental clones without specific selection^87^. These observations are similar to our observation with EpiTrace. In our analysis, although we sampled only a fraction of the tumor, the similar cell age estimated for MDM4^+^/EGFR^+^ and MDM4^+^/PDGFRA^+^ clones suggested that neither of these clones gained selective advantage during tumor growth. Instead, they are under neutral evolution. Further experiments with higher resolution clonal tracing, putatively with a genetic marker, are necessary to confirm this observation.

## Discussion

We formulated a model of clock-like chromatin accessibility change during cell mitotic aging, leading to the discovery of a universal epigenomic hallmark during cellular development: chromatin accessibility across clock-like loci. The heterogeneity of chromatin accessibility across clock-like loci is reduced at each mitosis, resulting in a converged, homogeneous activity pattern. We showed that chromatin accessibility changes act upstream of clock-like DNA methylation changes. Counting the fraction of opened clock-like loci of each cell gives a simple, phenotypic measure of cell mitotic age. We leveraged this measure to build a tool, EpiTrace, to predict cellular mitotic age. Furthermore, we showed that the similarity across clock-like loci chromatin accessibility between single cell clusters can serve as an accurate distance measure for phylogenetic analysis.

The DNA methylation shift in ClockDML is widely accepted as a hallmark of aging. However, the molecular mechanism generating age-dependent DNA methylation is yet unknown. Our data indicated that sample age predicted by chromatin accessibility and DNA methylation were significantly correlated, suggesting that they are possibly under the control of a similar biological process.

Chromatin accessibility changes accompany development and cell fate transition. However, we noticed that chromatin accessibility on clock-like loci is phenotypically neutral – that is, irrelevant to cell phenotype – based on several lines of evidence. First, EpiTrace age measured on the same set of clock-like loci generated from one tissue lineage (for example, the ClockDML from human PBMC) works for different lineages. Second, the EpiTrace age of a single cell correlates with its accumulative mitosis number instead of developmental maturity (for example, in the case of neuronal development). Third, clock-like loci derived from one species could be used to predict single cell age in another species. The exact molecular mechanism controlling how clock-like differential DNA methylation occurs on clock-like loci (to generate ClockDML), and how clock-like chromatin accessibility emerges on these loci, is an interesting question awaiting future investigation.

It is unexpected for the authors that the phylogenetic tree built with chromatin accessibility on clock-like loci (EpiTrace phylogeny) for single cell clusters of the same developmental lineage is highly accurate. EpiTrace inferred age is comparable to pseudotime-inferred cell “developmental time” but with higher resolution and less variation (Supplementary Figure 33). In fact, we noticed that such phylogenetic trees sometimes outperform those built with the highly variable peaks from scATAC data in terms of accuracy. Despite the fact that ClockDML (and clock-like loci) are highly enriched in cis-regulatory regions, there is no functional enrichment of them in specific developmental pathways or specific types of genomic elements (in addition to active cis-regulatory elements). Furthermore, this phenomenon is also phenotypically neutral. These results not only suggest a consistent birth sequence of cell types within the lineage but also indicate that senescence is a defined molecular process across cell types.

The current study is not without limitations. We noticed that the quality of inferred mitotic age by the current EpiTrace algorithm is dependent on sequencing depth (Supplementary Figures 34-36). EpiTrace could be less accurate when working on cells with low sequencing depth. Additionally, EpiTrace could be inaccurate when the starting cell population is highly imbalanced (Supplementary Figure 23). This phenomenon is related to single cell population heterogeneity across development, the nature of chromatin accessibility shifts during mitotic aging, the statistical model underlying our algorithm, and the limitations of the sequencing technique. Finally, estimation accuracy and the computational efficiency of EpiTrace heavily rely on the enrichment of clock-like loci in the initial reference loci set. In this view, the full set of scATAC peaks or solo-WCGW sites, albeit they may contain clock-like loci, are not sufficiently enriched (Supplementary Figure 37). As a result, inferring cell age from these reference loci is not as accurate as using the ClockDML set we provided for the algorithm (Supplementary Figure 38). Furthermore, using these loci as references is extremely computationally inefficient and renders single cell dataset analysis virtually impossible. Future improvement of the algorithm might require in-depth study of molecular mechanisms driving age-dependent chromatin accessibility changes on clock-like loci, improvements in the algorithm to adapt with low-quality and highly imbalanced datasets, and improvements in computational efficiency.

In conclusion, we discovered mitosis-associated, age-dependent chromatin accessibility on clock-like loci, which usually harbors ClockDML, and developed computational methods to track single cell age. By comparison studies, we showed that the chromatin accessibility-based mitosis age measure complements somatic mutation, RNA velocity and stemness predictions to predict the cell evolution trajectory with improved precision and power. We expect EpiTrace to be a useful tool for single cell studies for delineation of cellular hierarchies and organismal aging.

## Methods

### Human and animal biospecimens

This study was conducted in accordance with the local legislation, measures of the Declaration of Helsinki, and the Ethics Protocols of Human Genetic Resource Preservation Center of Hubei Province, China (Hubei Biobank). For the human study, blood samples from healthy donors (total n = 71 including: training n = 24; validation n = 47) used in this study were pretreated and preserved by the Hubei Biobank, approved by the Institutional Ethical Review Board (approval number: 2017038-1 and 2021125). Informed consent was obtained from the donors and their guardians. For the mouse study, 74 C57BL/6 mice were used to collect tail tissue (including: training n = 37; validation n = 37), and pregnant C57BL/6 (n = 4) mice were used to collect primary embryonic fibroblasts at Zhongnan Hospital of Wuhan University, which was approved by the Institutional Ethical Review Board (approval number: ZN2022246).

### Datasets used in this study

Datasets used were listed in Supplementary Dataset 2. Quality control basic statistics of the datasets from which we could obtain raw data were listed in Supplementary Dataset 2.

### Cell culture

Isolation of primary mouse embryonic fibroblasts (MEFs) was performed as previously described^90^. Briefly, E13-14 embryos were dissected from the uteri of pregnant C57BL/6 mice, separated from their yolk sac, and homogenized with scissors in 0.25% trypsin-EDTA. Homogenized embryos were aspirated in DMEM supplemented with 10% FBS and 1% penicillin and streptomycin. Primary MEFs were maintained in the same medium. HEK293-dCas9-p300 cells were a gift from Dr. Yi Rao, Peking University, Beijing, China. Immortalized MEF cells were a gift from Dr. Hui Jiang, National Institute of Biological Sciences, Beijing, China. All immortalized cells were maintained in DMEM supplemented with 10% FBS and 1% penicillin and streptomycin.

### FACS sorting of cultured cells into different cell cycle phases

For cell sorting, live MEFs at different passages were stained with Hoechst 33342 (5 μg/mL, Invitrogen, H1399) at 37°C for 20 min in the dark. After trypsin digestion, MEFs were resuspended in PBS with 1% FBS and collected in a sterile tube with a cell-strainer cap (Falcon, 352235). Then, the MEFs were subjected to FACS sorting. MEFs are first gated for whole cells and cell debris (FSC and SSC), then for single cells (Horizon V450-A and Horizon V450-H), and lastly for cells in G1, S, or G2/M phase according to DNA content. The BD FACSAria III Sorter was used for cell sorting at the Flow Cytometry Core Facility of the Medical Research Institute, Wuhan University.

### Single guide RNA (sgRNA) perturbation

sgRNA targeting ClockDML G8 set loci was designed manually (Supplementary Dataset 3). The sgRNAs were synthesized and packaged into lentivirus by GenScript. HEK293-dCas9-p300 cells were transduced with the lentivirus sgRNA library at a low MOI (∼0.4). Puromycin was added to cells 2 days after transduction. Cells were collected for bulk bisulfite sequencing and RNA sequencing. DNA and RNA were extracted from the cells using an AllPrep DNA/RNA dual-prep kit (Qiagen Cat. # 80204). The relative sgRNA expression was quality-controlled by sequencing the PCR product of the sgRNA barcode region.

### Single-stranded bisulfite capture sequencing

Genome-wide CpG bisulfite capture sequencing was performed as in Xiao *et al.*^40^ and He *et al.*^91^ using the Tequila-7N protocol. In brief, genomic DNA was extracted from PBMCs, bisulfite-converted, poly-dT-tailed with terminal transferase, ligated with a poly-dA-extruded 3’ adaptor, linearly amplified for 12 cycles with a primer that complements the 3’ adaptor, and ligated with a 5’ adaptor with 7 N-nucleotide overhangs. The resulting adaptor-ligated single-stranded inserts were amplified with Illumina P5/P7 sequencing adaptor primers and captured with the EpiGiant probe set (Nimblegen, Cat. #07138881001) following the manufacturer’s protocol. Postcapture libraries were retrieved, amplified, quantified, and sequenced on an Illumina NovaSeq with PE150 format to 50 M reads.

### MArchPCR bisulfite capture sequencing

Mouse ClockDML CpG bisulfite sequencing was performed with the MArchPCR protocol. In brief, nested gene-specific primers targeting mouse MM285 ClockDML^49^ were designed by a machine-learning-assisted automated pipeline (to be described elsewhere in another publication). Genomic DNA was extracted from PBMCs, bisulfite-converted, poly-dT-tailed with terminal transferase, ligated with a poly-dA-extruded 3’ adaptor, and amplified by a set of 5’ gene-specific primer 1 (GSP1) targeting mouse MM285 loci and a 3’ common primer targeting the 3’ adaptor. A second round of amplification was performed with a set of 5’ gene-specific primer 2 (GSP2) targeting several base pairs 3’ of the GSP1 landing site and the same 3’ common primer. The resulting DNA library was then purified and reamplified with Illumina P5/P7 sequencing adaptor primers. Libraries were retrieved, amplified, quantified, and sequenced on an Illumina NovaSeq with PE150 format to 3 M reads.

### SHARE-seq on single cells

SHARE-seq was performed essentially following the original published protocol^92^ (https://www.protocols.io/workspaces/shareseq) with slight modifications to work on cryo-preserved cells on MGI sequencers. Adaptor and barcode oligos were synthesized (Supplementary Dataset 4) from Sangon, Shanghai, China. Tn5 protein from Novoprotein (Cat. #: M045-01B) was assembled with customized mosaic end adaptor oligo in house by mixing annealed adaptor oligo and the Tn5 protein at 1.5:1 molar ratio and incubated at room temperature for 30 mins. Cryopreserved cells were thawed according to the 10x Genomics protocol with buffer exchange, fixed with FA at a 1% w/v final concentration for 10 min, and quenched by glycine. Nuclei were isolated by using NIB buffer (10 mM Tris pH 7.5, 1 mM NaCl, 3 mM MgCl2, 0.1% Tween 20, 0.1% NP40, 0.01% digitonin, 0.75% w/v BSA) and washed with NIW buffer (NIB without NP40 and digitonin) twice. Tn5 tagmentation was performed with 25 pmol assembled Tn5 protein against 20,000 nuclei in tagmentation buffer (1x Tango buffer, 0.2% NP40, 32% DMF, supplemented with 0.008% digitonin, 0.08% Tween 20, 1.7% v/v NexGene RNase Inhibitor, and a full Merck protease inhibitor cocktail tablet per 500 µl) for 30 min at 37°C with shaking (500 rpm) in a rotating incubator. Reverse transcription, hybridization with barcode oligos, ligation of the oligos, crosslinking-pull-down, scATAC library amplification, cDNA amplification, re-transposition with the cDNA, and scRNA library amplification were performed as described in the SHARE-seqV2 protocol, with our modified oligo set. Sample multiplexing was performed by splitting the round-1 oligo barcode usage. In all experiments, 4-6 cell samples were multiplexed for each SHARE-seq preparation. DNA nanoballs were prepared with the MGI nanoball MDA kit following the producer’s protocol. Sequencing was performed on an MGI2000 sequencer with a modified cycle setting and customized primers. Sequencing setup was: 101 bp from Tn5 transposed 5’ end using TN5-read1-link1/TN5-read1-link2 primers, 8+8 bp (with 30 bp dark cycle) using APP-A barcode primer 2, MDA reaction, 99 bp from Tn5 transposed 3’ end using Tn5-read2 primer, and 8 bp using R1-oligo-R2-LK-RP primer. Sequencing depth was targeted to 500 M reads for a full SHARE-seq scATAC or scRNA library.

### Preprocessing of sequencing data

All bisulfite sequencing (methylation) data were processed exactly as described in He *et al.*^91^. Per-CpG DNA methylation frequency (beta) was computed with PileOMeth (v0.1.13-3-gca82747, https://github.com/bgruening/PileOMeth). All bulk ATAC and Cut-And-Run sequencing data were processed exactly as described in He *et al.*^91^, with the hg19 reference genome. The 10× scATAC sequencing data were processed by CellRanger-ATAC (v1.2.0) with the hg38 reference genome. 10x scMultiomic sequencing data were processed by CellRanger-ARC (v2.0.1) with the hg38 reference genome. 10x scRNA data were aligned to the hg38 reference transcriptome by kallisto (v0.46.1, https://github.com/pachterlab/kallisto) and converted to a splice/unsplice count matrix. SHARE-seq data were processed separately for the scRNA and scATAC datasets. SHARE-seq ATAC sequencing data were preprocessed by zUMI (2.9.7b) to barcode-corrected ubam, converted to fastq by samtools, and mapped to the GRCh38-mm10 hybrid genome (10x Genomics) or mm10 genome by bwa (0.7.17-r1188). Cell barcodes were extracted by sinto using the ‘nametotag’ command, and fragments were extracted by the ‘fragment’ command. SHARE-seq scRNA data were processed by StarSolo (version 2.7.10a_alpha_220818) using a sjdb with a 50 bp overhang, and the key parameters “-- soloCBposition 0_0 0_7 0_8_0_15 0_115_0_122 \ --soloUMIposition 0_18_0_25 -- soloCellFilter CellRanger2.2 8000 0.99 10 \ --soloCBmatchWLtype 1MM \ -- outFilterMultimapNmax 1 \ --outFilterScoreMinOverLread 0.1 \ -- outFilterMatchNminOverLread 0.1”. The output velocyto result was then used in downstream processing. Fragment files from these preprocessed ATAC data were combined and analyzed in ArchR (v1.0.1)^6^. Quality control, doublet removal, dimensionality reduction (LSI), clustering and peak finding were all performed in ArchR according to its reference. Annotation of cells from their original publications/dataset was performed without adjustment. Cells were not manually excluded from the analysis, except in three cases: 1. Aneuploid 3PN cells in the human embryonic development dataset; 2. PT3 cluster in the kidney scATAC dataset, which is donor specific; 3. CD34-500 Erythroid lineage in the *in vitro* CD34 differentiation dataset, of which the *EpiTrace* age estimation is aberrant due to imbalanced cell composition. Lift-over between reference genome sets was executed with easyLift (v0.2.1, https://github.com/caleblareau/easyLift). scRNA data were passed to Seurat^93^ (v4) for basic quality control, normalization, scaling, dimensionality reduction (PCA), and clustering. The Seurat object was converted to anndata format with SeuratObjects (v4.0.4, https://github.com/mojaveazure/seurat-object) prior to RNA velocity and CytoTRACE analysis. For ATAC and CUT&Tag data processing, the reads were aligned to hg19 reference genome using bwa and de-duplicated by sambamba (v0.5.4), prior to peak calling by MACS2 (2.2.7.1).

### Age-dependent clock-like DML

Age dependent, clock-like DMLs (chronology) were discovered in the training cohort, with correlating per-CpG beta values against sample donor age using an adapted code from scAge (Trapp *et al.* 2021^94^, and personal communication with A.Trapp). The cutoff of the absolute value of Pearson’s correlation between sample age and beta was set to 0.7. Known clock-like mitosis-associated DML from Yang *et al.*^28^ (mitosis) and development-associated DML from Zhou *et al.*^38^ (solo-WCGW) were also included for comparison. Enrichment of clock-like DML on cis-regulatory elements from human single cell ATAC^39^, hematopoiesis cell ATAC^42^, bladder cancer single cell ATAC^40^, and placenta single cell ATAC (Gong *et al.*, unpublished) datasets was performed with Fisher’s exact test in R. Clock-like DML were clustered according to their beta and sample age using *hclust* in R (4.1.3), and unsupervised classification was performed by *cutree*. The mean beta of each group of Clock-like DML was correlated with age. The group (G8) of clock-like DML with the best negative correlation between mean beta and sample age is shown in Figure 1b.

### Predicting sample age by DNAm

Age prediction by DNAm beta values was performed by either GLM trained according to Horvath *et al.*^25^ or a modified scAge code (TimeSeq model)^94, 95^. The training cohort did not include donors aged 0-18 years; thus, no log-transformation was performed for younger donors. For the GLM, the loci were filtered for sufficient coverage >30x. We performed PCA on the CpG DNAm matrix and selected only PC1 for age prediction. For the TimeSeq model, the loci were not filtered.

### Measuring the diversity of chromatin accessibility in the ClockDML region

ATAC peaks overlapping ClockDML were pulled with GenomicRanges *findOverlaps,* and peakwise counts, intersample peak means, medians, standard deviations and coefficients of variation (CVs) were calculated in the R (v4.1.3) package sparseMatrixStats (v1.6.0). The entropy of ClockDML chromatin accessibility was measured by first normalizing per-peak read counts to total counts over all peaks of interest to have a vector of probability of observing a read in peak x (p(x)) and computing the Shannon entropy as: Entropy = −1×sum(p(x)×log(p(x)).

### Measuring the general accessibility of the ClockDML region

EpiTrace measures the total accessibility of given reference (ClockDML, or clock-like loci) region. ATAC peaks overlapping ClockDML were pulled with GenomicRanges *findOverlaps,* and peakwise counts were optionally censor-normalized according to Qiu *et al*.^96^. ClockAcc, defined as the number of total opened DMLs, was measured for each cell for peaks with nonzero read coverage. The algorithm then perform cellLcell-similarity-based reaction-diffusion smoothing to leverage information from other cells to denoise ClockAcc. To do this, cellLcell similarity was first calculated. The inter-single-cell dispersion of each peak was calculated and used to select the top 5% of variable peaks. The cellLcell similarity (correlation) matrix was computed as the correlation of the top 5% of variable peaks between each single cell using WGCNA::cor (v1.70-3). The correlation matrix was further thresholded by its mean, with correlation values lower than the mean being set to zero. Then, the correlation vector for each cell was normalized by dividing the total sum of correlation, denoting the cell-cell similarity index. Assuming the mitotic age acts on the phenotype of cell (manifested as a cellLcell-similarity matrix) to result in smoothened ClockAcc, the mitotic age of cells could be solved by nonnegative least squares. Nonnegative least square decomposition of the cellLcell-similarity matrix and smoothed ClockAcc was computed by the Lawson-Hanson algorithm using nnls::nnls (v1.4). Reaction-diffusion regression of the NNLS result was performed using an HMM-like approach: in each iteration, the NNLS result is updated to the weighted sum of the current NNLS result and the cross product of the NNLS result with the cellLcell similarity matrix until the difference between the updated NNLS result and the current NNLS result is lower than the preset threshold. The cross product of the final NNLS result with the cellLcell similarity matrix results in a smoothed, regressed measure of ClockAcc. The overall algorithm was first proposed in CytoTRACE^3^. The resulting smoothed total chromatin accessibility was renormalized between 0 and 1 to give a rank of samples, Such rank denotes the relative birthday (mitotic age) of the cell within the population. For bulk sequencing, the rank is reversed as (1-rank).

### Iterative algorithm to measure the general accessibility of the ClockDML region in single cell ATAC data

Single cell ATAC data are under-sequenced and sparse. Hence, reads falling on known reference clock-like loci (or ClockDML) might be few for individual cells. We use correlation to pick additional genomic loci whose chromatin accessibility strongly correlates with the initial estimated cell age, then use them as additional “reference clock-like loci” to boost the algorithm performance. Briefly, after the renormalization of smoothed total chromatin accessibility, the rank is not reversed. For iteration, read counts of all ATAC peaks were correlated against single cell age estimation (the rank). The correlation coefficient was scaled, and peaks with a sufficiently high correlation coefficient vs age (normally Z>2.5 or 3) were selected. A new “reference clock-like loci” set was built as the union of high-correlation peaks and the previous reference clock-like loci. In the next round of iteration, this new “reference clock-like loci” set replaces the previous reference for computation. The iteration was performed for designated times or until convergence of age estimation. Final single cell *EpiTrace* age estimation was given by the rank of smoothed total chromatin accessibility over clock-like loci in the final iteration. No reversal of ranking was performed for single cell datasets.

### Phylogenetic tree construction with chromatin accessibility in the ClockDML region

A mitosis birth sequence-based phylogenetic tree was inferred with chromatin accessibility on ClockDML in Seurat with the function BuildClusterTree and rooted using package ape (v5.6-1) with the cell cluster that has minimal mean *EpiTrace* age serving as an outgroup.

### Association of chromatin accessibility with estimated cell age

The read counts of all ATAC peaks were correlated against sample age estimation with the R (v4.1.3) package WGCNA and BiocParallel (v1.28.3). Correlation coefficients were scaled by R. Age-dependent chromatin accessibility trends of loci with significant correlations were analyzed with tradeSeq (v1.8.0). To generate a heatmap for visualization, chromatin accessibility of the same locus is averaged for cells of similar age bin (0-100) and phenotype.

### UMAP projection and cell transition trajectory for embryonic ATAC data

The ATAC data were collected from public databases (PRJNA494280^45^ and PRJNA394846^46^) and in-house placenta ATAC data. After alignment, peaks were called for each data point and merged to become a union ATAC peak set. The union peak set was then used as the input feature set. A read count matrix (peak x cell, 253,541 x 44) was constructed and normalized by Signac::RunTFIDF(). A total of 250,290 variable features were identified by Signac::FindTopFeatures (min.cutoff = 50). Dimensional reduction was performed with Signac::RunSVD() and UMAP projection by Seurat::RunUMAP (reduction = ‘lsi’, dims = 3:6). A pseudotime trajectory was constructed using slingshot, and an arrow was drawn according to the smoothed trajectory.

### Cross-mapping of ClockDML between species

When genome synteny lift-over chains are available (human-mouse, human-zebrafish, mouse-human, mouse-zebrafish), lift-over between genomes for the ClockDML was performed with UCSC LiftOver executable with the respective chain file. For fly human mapping, without available lift-over chains, ortholog pairs were identified bioinformatically using the FlyBase database. Human orthologous genes with ClockDML present in their promoter were identified. Then, the corresponding promoters in the fly ortholog genes were taken as “human-guided” fly clock-like loci.

### Comparing 8CLC-development cluster hierarchy inference results with known results

We first built a “real cluster hierarchy” based on RNA expression of known 4CL (TET1/NCOA3/FGFR1/HYAL4/WNT3/DNMT3L/LEFTY1/DHCR24), 8CLC (DPPA3/ZNF280A/TPRX1/ZSCAN4/DUXA) marker genes, extracellular matrix markers (FN1/COL1A1/COL3A1/EMP1/CDH11), and the cluster-specific differentially expressed genes. The clusters were individually ranked according to EpiTrace age and scVelo pseudotime, and each rank was subsequently correlated with the “real cluster hierarchy”.

### Mitochondrial SNV (mGATK) analysis

mGATK analyses were performed following Lareau *et al.*^8^ to reproduce the clones described in the original publication.

### Clonal age inference and expansion ratio estimation

For each mtscATAC cell myeloid-biased clone from CD34_800 experiment, the clone was first classified by whether it harbored only progenitor (prog/prog_my) cells (Prog-only), intermediate (my_1) cells (Int-only), or both. The difference in the total numbers of terminal cells (my2, my3, and my4) in this clone on latter timepoints (D14/D20) from that on initial timepoint (D8) was taken as the “expanded cell number”. The mean EpiTrace age of cells on initial timepoint (D8) was taken as the “initial age of this clone”. Finally, the log-expanded cell number was regressed against the initial age for each clone.

### RNA velocity and CytoTRACE analysis

Spliced and unspliced scRNA data were exported from R (v4.1.3) with SeuratObjects to h5ad format and imported into Python (v3.9.7) to be processed by CellRank (v1.5.1, https://github.com/theislab/cellrank)^69^. The scVelo and CellRank CytoTRACE kernels were used for estimating RNA velocity and CytoTRACE score, respectively. scVelo (v0.2.5, https://github.com/theislab/scvelo)^97^ with Scanpy (1.9.3) was used to produce velocity embeddings in UMAP space.

### Multiomic derivation of cell evolution trajectory with kernel combining cell age and RNA velocity

A custom kernel of CellRank built upon *EpiTrace* age was used to project *EpiTrace* age into CellRank. A combined kernel of RNA velocity and *EpiTrace* was built with 0.75×velocity + 0.25×age, which was used in the downstream GPCCA estimator in CellRank. Lineages were calculated by GPCCA, with cell clusters with the least age designated root and cell clusters with the maximal age designated terminal. Velocity genes were computed by the GPCCA estimator for a given lineage.

### Transcription factor motif differential analysis

TFBS motif differential activity was determined by Signac (v1.5.0, https://github.com/timoast/signac)^98^ using the same *EpiTrace* object built during *EpiTrace* analysis.

### Copy number variation analysis

scATAC copy number variation was called with CopySCAT (v0.3.0, https://github.com/spcdot/CopyscAT)^77^ and annotated with ChIPseeker (v1.30.3)^99^.

### Long-read NanoNOME-seq data analysis

Processed methylated C calling data were downloaded from NCBI (GSE183760^56^). Methylation calls were performed to call a base “methylated” with log_lik_ratio >1 or “unmethylated” with log_lik_ratio < −1. All bases with log_lik_ratio between −1 and 1 were called “unknown”. Nucleosome-sized (147 bp) flanks were taken from each ClockDML site, and the CpG and GpC methylation states were called for each CpG and GpC site within this region. A region is defined as “opened” if the fraction of “methylated” GpC exceeds 20%. For pseudotime analysis, methylation-per-base information of the reads from the same Cas9-targeted region was used to construct a matrix, which was then used by TSCAN (1.34.0) to compute a pseudotime evolution sequence of the reads.

### scCOOL NOME-seq data analysis

Sequencing data from human embryonic cell scCOOL-seq experiments^55^ were mapped to the hg19 reference genome using bwa-meth. Per-read CpG and GpC coclassified epireads were extracted using Pile-O-Meth (0.1.13-3-gca82747 using HTSlib version 1.2.1). Reads overlapping the ESC mitosis-related EpiTOC^28^ loci were extracted. For each stage of cells, the mean CpG and GpC methylation levels on these reads were calculated and summarized.

### BSPCR data processing and correlation to the human ClockDML aging coefficient

The BSPCR data from PRJNA490128^57^ were downloaded from NCBI and mapped to the GRCm38.75 genome with bwa-meth. The CpG methylation levels were extracted with Pile-O-Meth. The sequenced CpG loci were mapped back to the human genome using the mm10-hg19 chain using LiftOver. Transactivation-mediated DNAm shifts in the mouse loci were then compared to the human age-dependent DNAm shift coefficient.

### Pseudotime trajectory inference (slingshot, VIA, Monocle, Palantir)

Slingshot, VIA, Monocle2, Monocle3, and Palantir computations of the mtscATAC dataset were performed exactly as the software instructions mentioned.

### Gene set enrichment (GO)

Genes from the clustering result of the anti-PD1 scATAC dataset (C1, C2, and C3) were used for GO enrichment analysis using the R package clusterProfiler^100^ (4.4.4) with the function ‘enrichGÒ. Pathways with adjusted P-values < 0.05 were considered significant.

### Region overlap enrichment

Overlap enrichment between regions were tested by Fisher’s exact test. For random permutation, background random loci were permuted for indicated (100 or 1,000) times using regioneR (1.28.0).

## Code and data availability

*EpiTrace* and accompanying testing data are available at https://github.com/MagpiePKU/EpiTrace. Tutorials and usage manuals for *EpiTrace* are provided on publicly accessible website (https://epitrace.readthedocs.io). Codes to reproduce the analysis, along with the original and processed data for reproducibility purpose, are provided on OSF (https://osf.io/8xd2p, doi:10.17605/OSF.IO/8XD2P). A full list of downloaded and generated data used in this study, including their access IDs, references, usage in this paper, and/or download links, is provided in Supplementary Dataset 2. Accession codes for the publicly available datasets in this study are: (NCBI): PRJNA494280; PRJNA394846; GSE178969; GSE190130; GSE178324; GSE178966; GSE142745; GSE129785; GSE162170; GSE166547; GSE139136; GSE163655; GSE163656; GSE74912; GSE89895; GSE179606; GSE65360; GSE163579; GSE137115; GSE152423; GSE164978; GSE100272; GSE183760; PRJNA522707; GSE102395; GSE103590; GSE121862. (CNGB): CNP0001454. Chemical induction of 8-cell-like state data is downloaded from https://figshare.com/s/ff707bf8242f7b3ed8f5, https://figshare.com/s/760d3ff54f1214a50cc2 and https://figshare.com/s/9c01c3b58d34b80de230. mtscATAC data is downloaded from https://github.com/caleblareau/mtscATACpaper_reproducibility. anti-PD1 treated cancer biopsy data is downloaded from https://github.com/GreenleafLab/MPAL-Single-Cell-2019. Cortical scMultiome data is downloaded from https://github.com/GreenleafLab/brainchromatin. In-house generated datasets are uploaded to the OMIX database in CNGB, China: human data: OMIX005823; and mouse/in-vitro data: OMIX005824. No restrictions would be applied on using the in-house generated mouse and *in vitro* datasets. Due to local legal requirements, the in-house generated human dataset would only be available in processed data format and requires a case-by-case application through the China Human Genetic Resources Management Office.

## Author contributions

Conceive of idea: YX, WJ, XW, YZ; Algorithm development: WJ, YZ; Biological specimen collection: YX, LJ, GW, KQ, XW; Data collection: JF, FC; Cell culture experiment: LJ, MY; Sequencing experiment: YX, WJ; Bioinformatic analysis: WJ, JF, FC, YZ; Biology and clinical analysis: YX, WJ, YZ; Writing the manuscript: YX, WJ, LJ, KQ, YZ; Supervised the study: XW, YZ.

## Acknowledgments

The non-binary scAge DNAm-based age prediction code was kindly provided by Dr. Alex Trapp. The authors would like to thank all the authors sharing publicly available datasets used in this study. The excellent technical assistance of Ms. Yayun Fang and Ms. Danni Shan at Wuhan University is gratefully acknowledged. The authors would like to thank Drs. Yi Rao and Hui Jiang for reagents. The authors would like to thank Drs. Yangyang Liu, Junqing Ye, Xuepeng Chen, Ge Gao, Jianrong Yang, and Zhenglin Du for discussion. We would also like to thank Dr. Xiaoling Li at the National Institute of Environmental Health Sciences (NIEHS) for the helpful discussion and critical reading. Part of the analysis was performed on the High Performance Computing Platform of the Center for Life Science (Peking University). This study is supported by grants from the National Natural Science Foundation of China (82273065), the Research Fund of Zhongnan Hospital of Wuhan University (SWYBK01-02, YKYXM20210105), and the Fundamental Research Funds for the Central Universities (2042022dx0003). The funders played no role in the study design, data collection and analysis, decision to publish, or preparation of the manuscript.

