## Supplementary Information for "Tracking single cell evolution via clock-like chromatin accessibility"

Supplementary Notes: Pages 2-8

Supplementary Figures: Pages 9-64

List of Supplementary Datasets: Page 65

References for Supplementary Information: Page 66

### Supplementary Notes

#### Chromatin accessibility on ClockDML correlates with sample age in bulk ATAC-seq data

We measured chromatin accessibility on ClockDML within a well characterized developmental hierarchy – hematopoiesis – with bulk ATAC-seq data from fluorescence activated cell sorting (FACS) -sorted blood cells[1]. Interestingly, cross-ClockDML chromatin accessibility profile segregates samples according to their developmental stage (Supplementary Figure 2a). Along the axis of differentiation from hematopoietic stem cell (HSC) towards terminally differentiated blood cells, we found decrease of inter-sample per-loci chromatin accessibility variation (Supplementary Figure 2b), mean chromatin accessibility (Supplementary Figure 2c), and within-sample cross-loci chromatin accessibility entropy (Supplementary Figure 2d). As a result, per-sample cross-peak coefficient of variation increases along the axis of differentiation (Supplementary Figure 2e). Such phenomenon might correspond to overall chaotic nature of transcription in undifferentiated cells[2], as the percentage of reads overlapping ClockDML within the whole ATAC-seq library gradually increases over differentiation[1, 3, 4] (Supplementary Figure 2f), suggesting a canalized epigenome with limited openings in terminally differentiated cells. Together, these results indicate that ClockDML are more accessible in cells of lower mitotic age compared to cells of higher mitotic age. Furthermore, the overall accessibility pattern is more random in lower mitotic age (Figure 1a and Supplementary Figure 2).

#### Establishment of algorithm to measure total variable chromatin accessibility on ClockDML

We firstly validated whether ClockDML chromatin accessibility is correlated with cell mitotic age. To this end, we developed a computational algorithm, EpiTrace, to measure the total fraction of opened ClockDML (ClockAcc) in each sample and compare/rank them across samples. To reduce technical background noise from less variable ClockDML regions, the algorithm selects the most variable ATAC peaks across all ClockDML within the dataset to produce a smoother output (smoothened measure of variable total ClockAcc, smoothedClockAcc).

smoothedClockAcc negatively correlates with developmental stage in the hematopoiesis bulk ATAC dataset[1] (Supplementary Figure 4a). In another test case, we tested induced pluripotent stem cells (iPSC) which shows age resetting during cell fate reprogramming. In concordance with previous studies with DNAm-based age estimation, smoothedClockAcc is higher in iPSC compared to their ancestor lymphoblastoid cells or differentiated cardiomyocytes (Supplementary Figure 4b), indicating a reset-ed cell age during iPSC induction.

We predict a relative cell mitotic age, EpiTrace Age, within these samples by computing cell-x-cell similarity matrix, performs iterative diffusion-regression cycle for ClockAcc to reduce the variation of ClockAcc within similarity blocks of cells, and finally ranking the resulted regularized values across all samples to produce a normalized cell age index between 0 and 1. As mentioned earlier, the total ClockAcc is negatively correlated with cell/sample developmental stage or mitotic age, and we used the reverse of rank for total ClockAcc (reverse EpiTrace age) as an indicator of mitotic age in bulk samples. In the hematopoiesis dataset, cells in developmental hierarchy (Supplementary Figure 5a) were accurately positioned according to reverse EpiTrace age (Supplementary Figure 5b). Furthermore, phylogenetic trees built with peaks overlapping ClockDML agrees with known developmental hierarchies (Supplementary Figure 5c). In iPSC induction dataset[3], the iPSC shows lower reverse EpiTrace age compared to its ancestor or its progenitor (Supplementary Figure 5d). In an additional bulk ATAC-seq dataset from PBMC and tumor-infiltrated CD8 cells[4], naive CD8 cells showed lowest reverse EpiTrace age, whilst tumor-infiltrated, non-naive CD8 cells showed highest reverse EpiTrace age (Supplementary Figure 5e). Together, these results suggested that the total fraction of opened ClockDML in bulk ATAC-seq data serves as an internal timer along cell division.

**Theoretical considerations in adopting the EpiTrace algorithm to single cell ATAC**

In ideal conditions, bulk ATAC sequencing of a single type of cell resembles a sampling process across all opening chromatin regions with replacement, with the probability of sampling over a given chromatin region correlates with ATAC peak height, because the same region from different individual cells could be sampled again and again. In contrast, single cell ATAC sequencing with limited reads resembles a sampling process without replacement, because a cell only has a limited number of copies of the same region, and it is highly unlikely that a single region is sampled twice under a very low total read depth. Heterogeneity of chromatin accessibility across the genome in young cells would result in sparse coverage across a wide range of genomic regions. In other words, whilst saturated sampling could rescue peaks with flat-and-wide shape in bulk ATAC-seq dataset, these regions would be likely to drop off in scATAC dataset. In contrast, peaks with high maximal height would be more likely to be successfully sampled in scATAC dataset. Therefore, ClockAcc should be positively correlated with cell mitotic age in scATAC-seq (Supplementary Figure 3). As a result, adopting EpiTrace to single cell ATAC data required removal of the reversal in ranking procedure.

Additionally, we consider the following three points: firstly, chromatin accessibility associated with age is unnecessary accompanied by DNA methylation changes, as data supported that chromatin accessibility is epistatically upstream of DNA methylation changes; secondly, despite that we determined chromatin accessibility on ClockDML is phenotypic neutral, it is unnecessary for all regions exhibiting age-associated chromatin accessibility are general for all cell types – that said, lineage-specific genomic loci with age-associated chromatin accessibility might exist, and furthermore these loci are unlikely to fall on ClockDML; thirdly and lastly, because of the sparseness nature of single cell ATAC sequencing, sequencing coverage across ClockDML might be sub-optimal. To enhance age determination accuracy, we perform iterations to update the "clock-like loci" list after age determination: correlation of all scATAC peaks against the predicted EpiTrace age, extract peaks with high correlation coefficient with cell age, and include them in the new "clock-like loci" set for a next round of analysis. Such iteration greatly improves performance of EpiTrace in single cell dataset.

**Details of EpiTrace algorithm**

EpiTrace was designed to estimate the total fraction of opened aging loci (i.e. "reference clock-like loci"). In brief, it takes in a set of reference clock-like loci and estimates the opened fraction of reference clock-like loci as cell age.

If iteration option is activated, the program then calculates the correlation between cell age and peak chromatin accessibility. The top correlated peaks were taken as new "putative reference clock-like loci", and the estimation procedure was repeated. This process is iteratively repeated until the estimated cell age converges or reaches a preselected limit.

In detail, the algorithm uses the following steps:

1. Prepare a matrix Mat where columns are single samples (hereinafter called cells) and rows are the reference peaks. Here, the initial reference peak set ('aging loci') is predetermined by prior knowledge, such as ClockDML in human/mouse, or ClockDML-homolog region in other animals. Other reference peak sets, such as putative G-quadruplex loci (pGQS), could be used, given user has a good biological knowledge of their nature.
2. Peaks without any reads were removed.
3. (optional) log1P-transform and perform census transformation or normalization of Mat. (We find that this step is not always necessary in most datasets)
4. Peaks that were expressed universally in the samples and had high variance were selected. First, the peaks that were expressed in >5% of cells were selected. Then, the dispersion coefficient is calculated as Vars/Means of each peak. Select the top 3,000 (default) peaks with the largest dispersion.
5. Perform the matrix x matrix correlation to calculate similarities between the cells (from the viewpoint of aging loci). This results in a correlation coefficient matrix:

$$CorM = corr(Mat)$$

Here in the matrix CorM, rows and columns denote the reference and target cells.
6. Calculate the average correlation coefficient as the mean (CorM)
7. Remove the diagonal elements (cell-to-itself autocorrelation) in CorM.

8. Elements in CorM that are lower than the mean (CorM) are set to zero. Therefore, the remaining non-zero CorM elements denote highly similar cell pairs.

9. Correlation coefficients are censor-normalized for each cell:

$$CenM = CorM/[rowSums(CorM)]$$

10. Remove those cells without any significant correlation to other cells (these are likely to be abnormal cells).

11. The correlation of each peak to the total accessible loci counts of each cell was calculated.

12. Select the top 200 (default) peaks that show the best correlation in step 12.

13. Calculate v, the per-cell average of reads falling on the peaks as in step 13. v is an intermediate tool variable that represents total accessible loci counts (in the reference loci set).

14. Perform diffusion smoothing (using an HMM) along the cell-to-cell similarity (the coefficients in CenM) for v until it converges. In other words, information from other cells is borrowed to make the noisy estimation of v more accurate.

15. According to our model, v is correlated with cell age. Rank v as an index of cell age x.

16. Since we initiated on a limited starting set of reference peaks (by overlapping scATAC peaks with ClockDML, or pGQS), it is possible that we do not include a complete set of aging loci in our analysis. Given our initial estimation of x, $x_0$, as the rank in step 16, we could then compute the correlation coefficient between peak reads and $x_0$ on all single cells (or samples). Peaks with correlation coefficients that ranked top among all peaks (with a Z score by the user's definition) are then considered new potential 'aging loci'.

17. Repeat the calculation from step 1 to obtain a new estimation of x, $x_1$.

18. Calculate the difference between $x_1$ and $x_0$.

19. If the difference is still larger than expectation (by user's definition), then repeat step 16 until the difference is less than what is expected or the iteration number is too large (by user's definition).

The inclusion of iteration is mainly required for single-cell ATAC datasets. Basically, we use iteration to include more potential clock-like loci as reference, to mitigate the under-sampling problem in single-cell ATAC-seq data. We would like to note that there is no ground-truth cell/sample age information provided to the algorithm. Instead, the algorithm simply leverages the fact that heterogeneity of chromatin accessibility on clock-like loci gradually reduces over cell replication. In fact, we noted that when using EpiTrace algorithm for bulk ATAC datasets, iteration swiftly converges and usually ends within 1-3 cycles, probably because such data are sequenced to saturation.

### Supplementary Figures

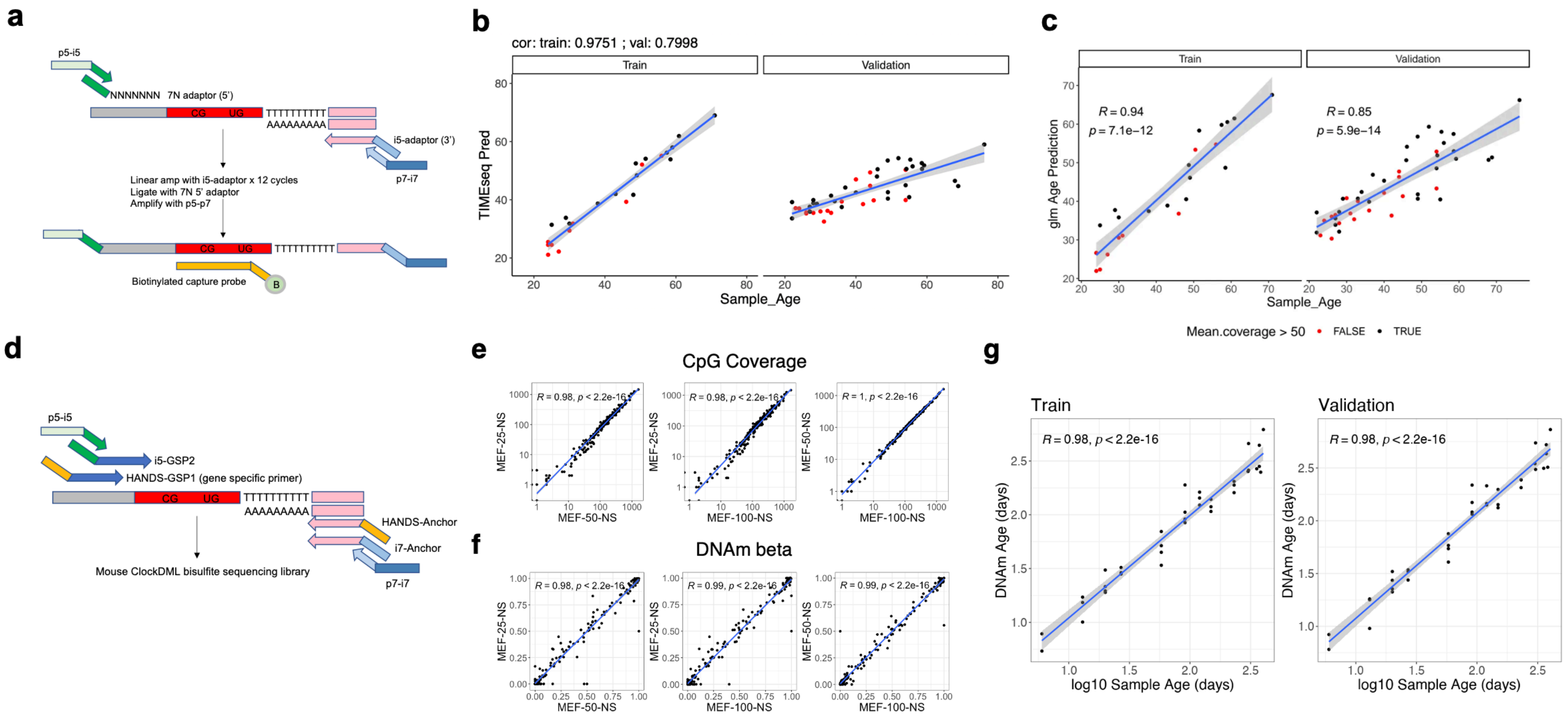


**Supplementary Figure 1. Using the DNA methylation level in the ClockDML set to predict donor age in human and mouse samples.**

**(a)** Schematic of the bisulfite sequencing protocol targeting human ClockDML. Bisulfite-converted DNA fragments (with methylated CpG as “CG” and unmethylated CpG converted to “UG”) were 3'-tailed with poly-dT with terminal deoxynucleotide transferase under the control of a tail-length-controller adaptor (pink) with poly-dA extrusion. Linear amplification with adaptor-specific primers was performed for 12 cycles. The linear-amp products were then ligated to a 5' adaptor with 7 random nucleotide overhangs and amplified to produce a double-stranded library. The library was captured by biotin-labeled probes targeting human ClockDML sites for sequencing. **(b)** TIMEseq prediction of sample age in the training set and validation set samples. **(c)** GLM-based prediction of sample age in the training set and validation set samples. **(d)** Schematic of the MArchPCR (multiplex, anchored PCR) bisulfite sequencing protocol targeting mouse ClockDML. Bisulfite-converted DNA fragments were poly-A tailed and ligated to a 3' adaptor as in (a) and then amplified with a common i7-anchor primer targeting the 3' adaptor and a mix of “GSP1” primers targeting the 5' of ClockDML. These primers are modified to have a common sequence (HANDS) in their 5' tail to reduce primer dimer products. Another round of PCR amplification was performed with another set of “GSP2” primers targeting a few base pairs 3' to the “GSP1” targeted sites and 3' common primers, resulting in a DNA library ready for sequencing. **(e)** Performance stability of CpG coverage with different amounts of starting materials (25 ng, 50 ng, 100 ng of genomic DNA). **(f)** Performance stability of CpG DNA methylation level measurement with different amounts of starting materials. **(g)** GLM-based prediction of mouse sample age in the training and validation sets

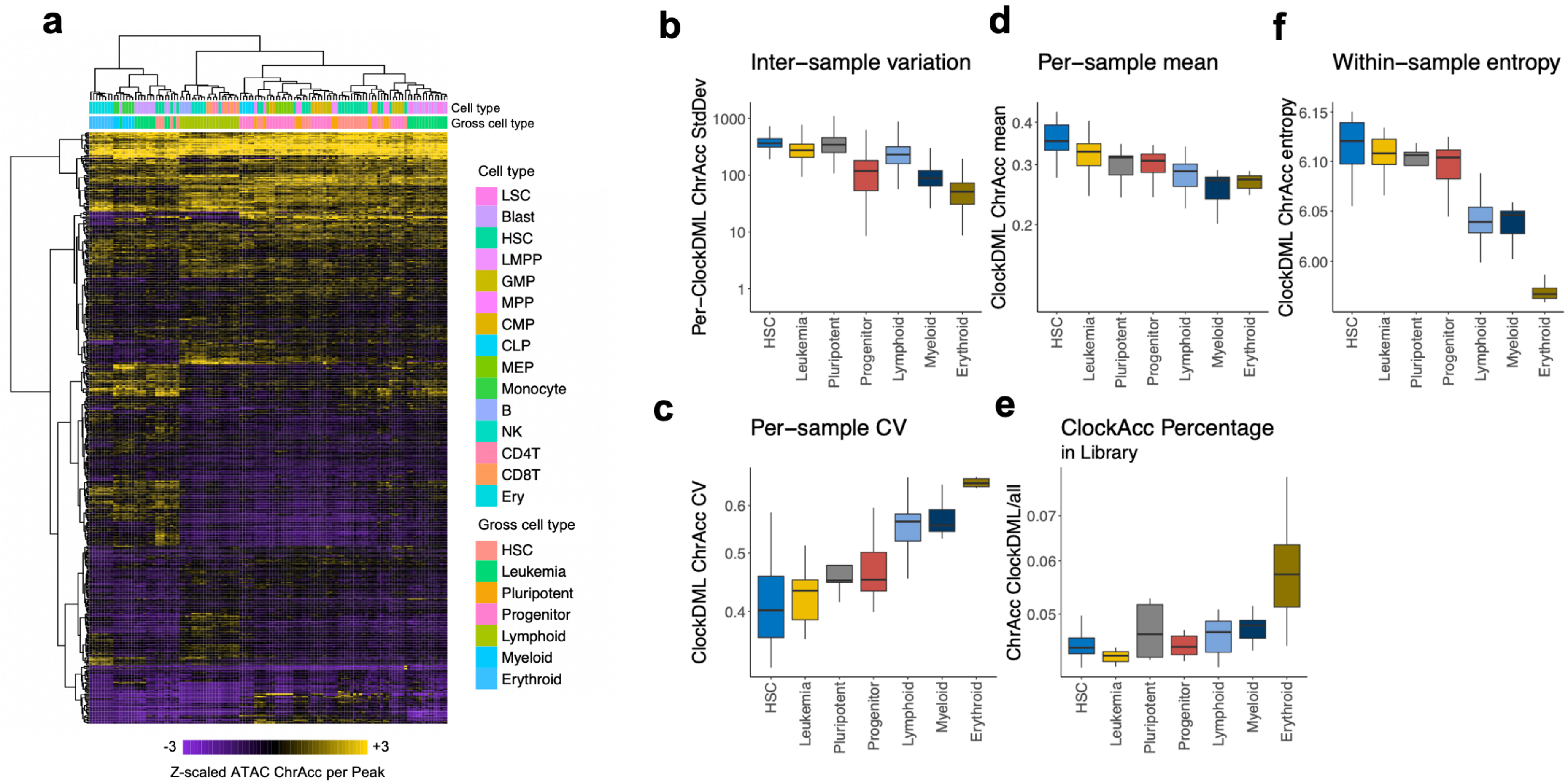


**Supplementary Figure 2. The ChrAcc pattern on ClockDML gradually converges during cell development.**

Bulk ATAC-seq of FACS-sorted blood cells overlapping the ClockDML region **(a)** was used to compute the intersample variation of ClockDML ChrAcc **(b)**, per-sample normalized ClockDML ChrAcc **(d)**, and per-sample ClockDML ChrAcc coefficient of variation **(c)**. The chromatin accessibility pattern is more chaotic in progenitor cells and gradually converges to a fixed pattern in terminal cells, as evidenced by the per-sample cross-peak entropy **(f)** and the increase in the percentage of reads originating from ClockDML-associated peaks **(e)**. Data: GSE74912.

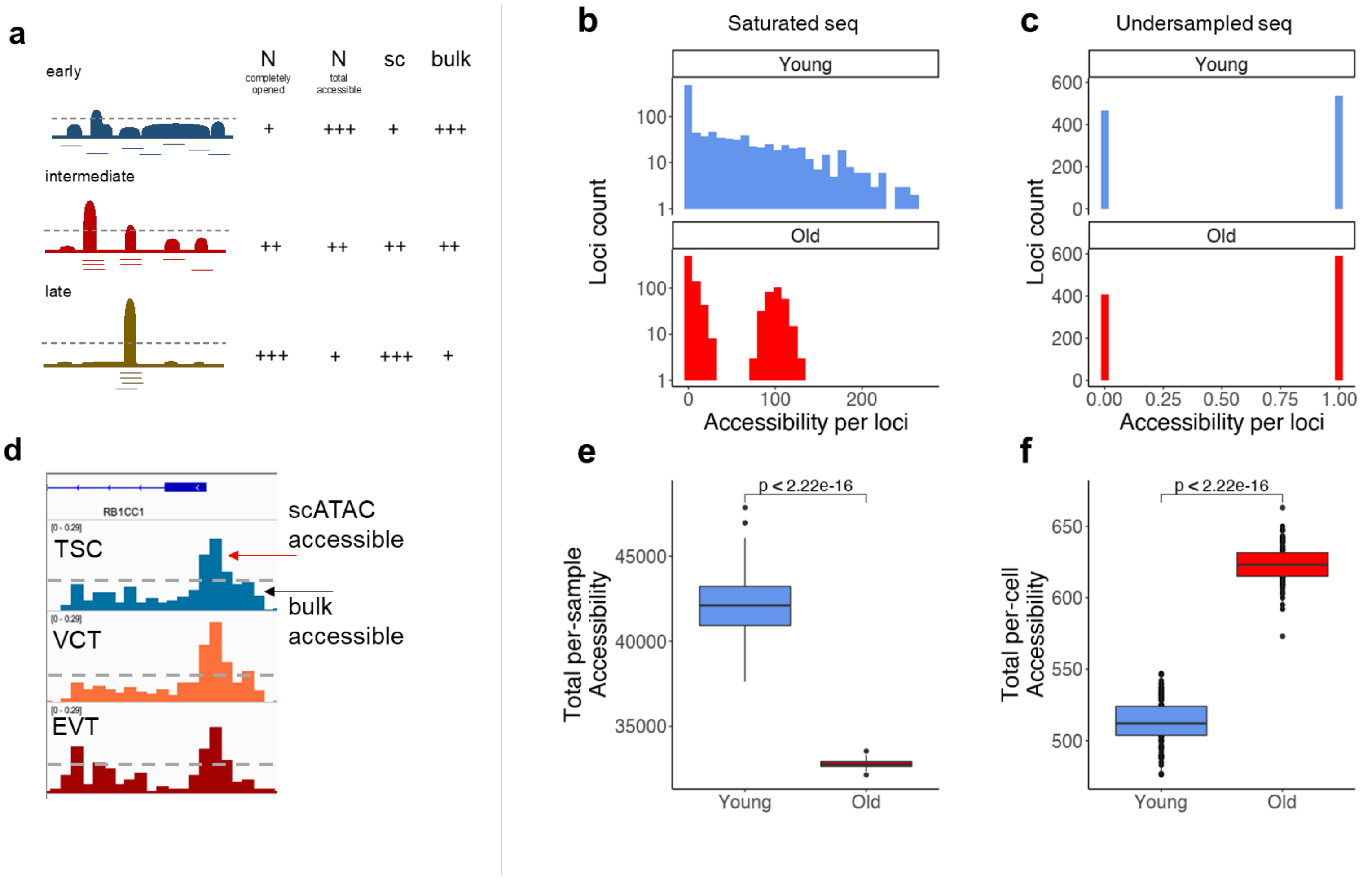


**Supplementary Figure 3. Undersampling in single cell sequencing results in the opposite of the observable total ChrAcc on ClockDML peaks compared to bulk ATAC sequencing.**

**(a)** Schematic of chromatin accessibility on ClockDML in early, intermediate or late-born cells. Top: aggregated chromatin accessibility across single cells from a similar population; bottom: single cell chromatin accessibility reads (each row represents reads from a single cell). Although accessible loci are more abundant in early cells, the completely opened loci are enriched in terminal cells. This results from the heterogeneity of single cell chromatin accessibility, which in turn suggests heterogeneity in a "pure" single cell population. **(b)** Aging loci were modeled such that in the "young" state, chromatin accessibility showed a low mean value but high variance, and in the "old" state, chromatin accessibility displayed a bimodal distribution, each with lower variance, in the low- and highly accessible states. **(c)** In downsampled ATAC-seq, such as single cell experiments, the genome is severely undersampled. The maximal sequenced fragment number from any locus is limited (one in this case). **(d)** Pseudobulk scATAC signals on a ClockDML around RB1CC1 from human placental trophoblast stem cells (TSCs), villous cytotrophoblasts (VCTs) and extravillous trophoblasts (EVTs). Completely opened loci are schematically shown above the gray threshold line. Data from Gong et al., unpublished. **(e)** In bulk ATAC experiments, total per-sample chromatin accessibility on aging loci decreased in old samples. The simulation is iterated 100 times. **(f)** In undersampled experiments such as single cell ATAC, total per-cell chromatin accessibility on aging loci increased in old samples. The simulation is iterated 100 times. Group statistics: t-test. Correlations: Pearson's.

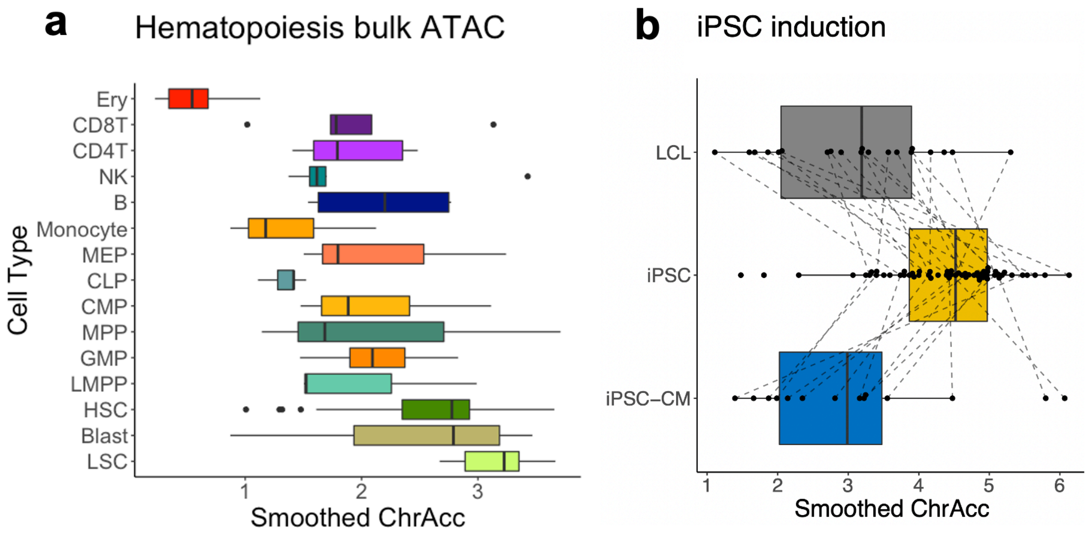


**Supplementary Figure 4. ClockDML ChrAcc in bulk samples.**

**(a)** Smoothened ChrAcc on FACS-sorted HSCs. Data: GSE74912. **(b)** Smoothened ChrAcc on lymphoblastoids, iPSCs, and iPSC-cardiomyocytes. Data: GSE89895.

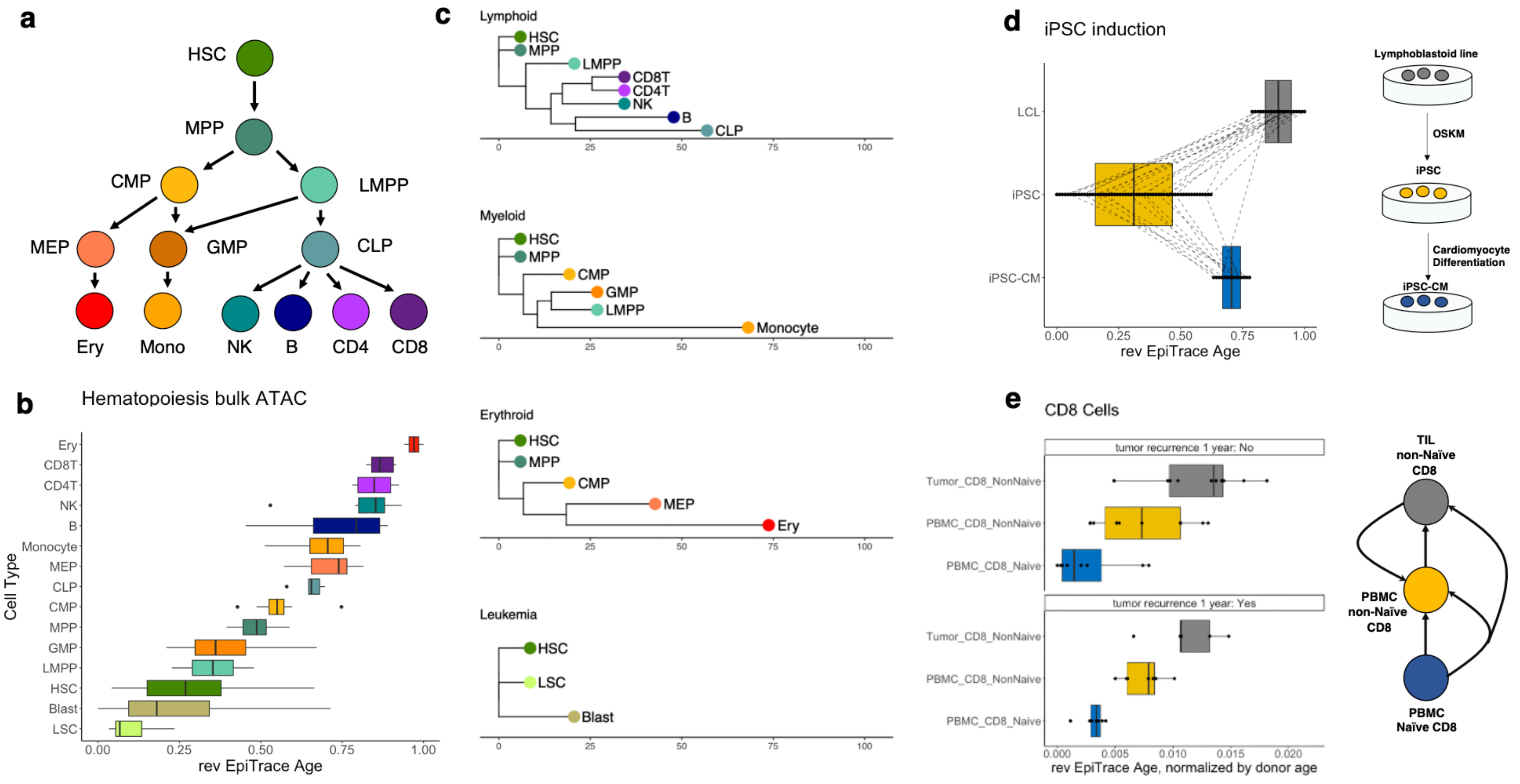


**Supplementary Figure 5. Bulk ATAC cell age estimation using reverse EpiTrace age.**

**(a)** Schematic diagram of hematopoiesis. **(b)** Sample age (x-axis, reverse EpiTrace age) of bulk sample ATAC-seq data from FACS-sorted normal and leukemic cells. **(c)** EpiTrace phylogeny of hematopoietic sub-lineages. **(d)** Reverse EpiTrace age of bulk ATAC-seq of lymphoblastoid cells (LCL), derived iPSCs (iPSC), and iPSC-derived cardiomyocytes (iPSC-CM). Samples from the same donor across different time points are grouped (lines). **(e)** Reverse EpiTrace age of FACS-sorted, naïve or non-naïve CD8 cells from PBMCs or tumor tissue. Data: GSE74912

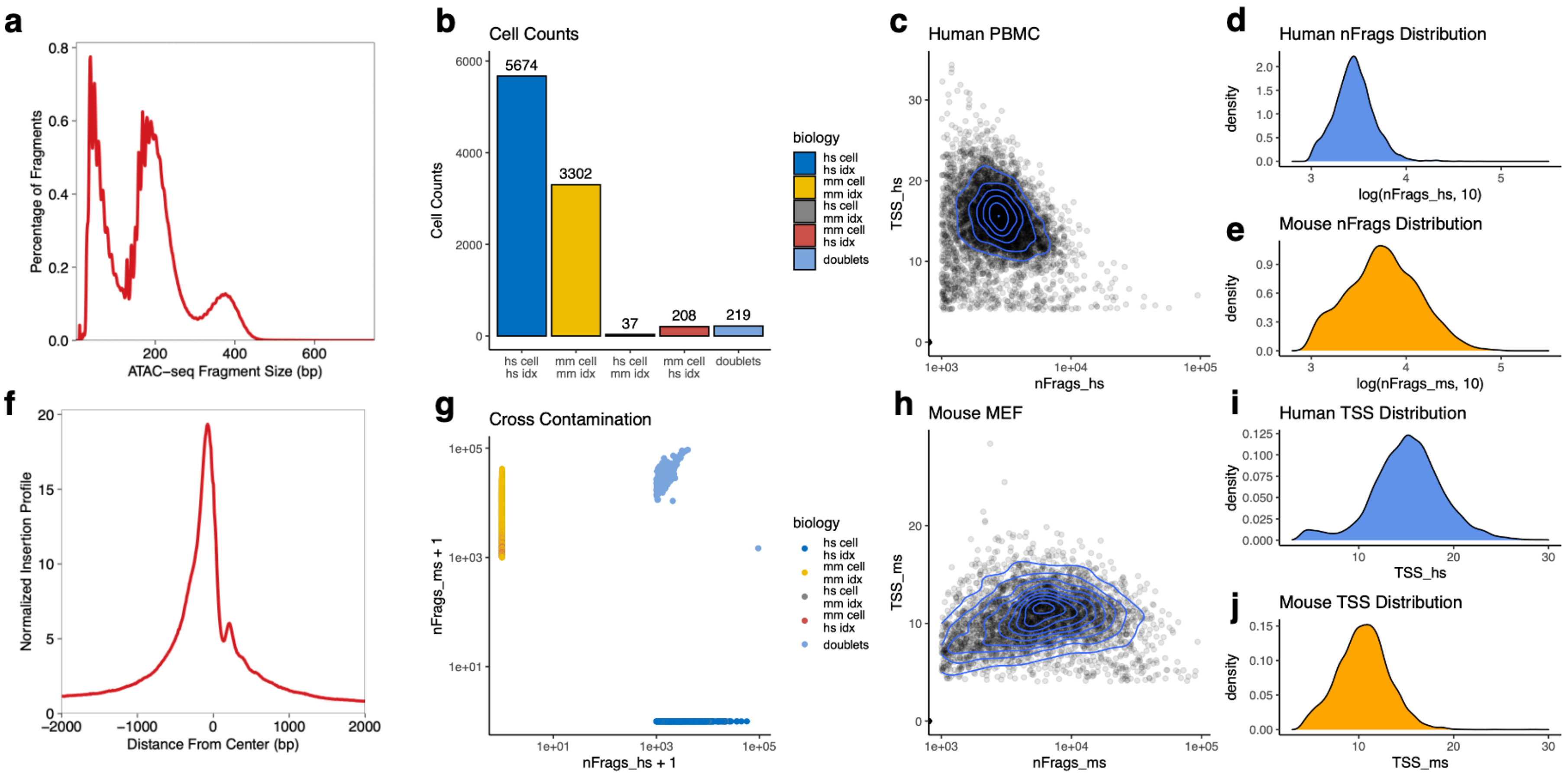


**Supplementary Figure 6. Assay validation of SHARE-seq using human/mouse mixed samples**.

Human PBMCs from one donor and mouse immortalized MEF cells were used for the SHARE-seq assay. During the first round of single cell barcode hybridization, human and mouse cells were applied to different barcodes, such that the first-round single cell barcodes served as a species-specific sample index. Single cells were separated by the combination of three round single cell barcodes. The sequenced reads were mapped to a customized hg38+mm10 hybrid genome using STARsolo (RNA) or a customized pipeline with zUMI/bwa/sinto. Mapped fragments were analyzed by Seurat (RNA) or ArchR (ATAC). **(a)** Overall fragment size distribution of ATAC reads. **(b)** Number of single cells classified by whether their genome mapping result was strictly human (hs cell), strictly mouse (ms cell), and whether their sample index was human (hs idx) or mouse (ms idx). Single cells with reads mapping to both the human and mouse genomes were defined as doublets. Mixed-up cells were defined as cells with different origins (sample) compared to their content (mapping result) and consisting of < 5% of all cells. **(c)** Total reads (nFrags) and TSS enrichment score for human cells. **(d)** Distribution of total reads (log) for human cells. **(e)** Distribution of total reads (log) for mouse cells. **(f)** Overall ATAC-seq coverage relative to TSS. **(g)** Single cell reads that mapped to the human (X-axis) or mouse (Y-axis) genome, log-transformed. **(h)** Total reads (nFrags) and TSS enrichment score for mouse cells. **(i)** Distribution of TSS enrichment scores for human cells. **(j)** Distribution of TSS enrichment scores for mouse cells.

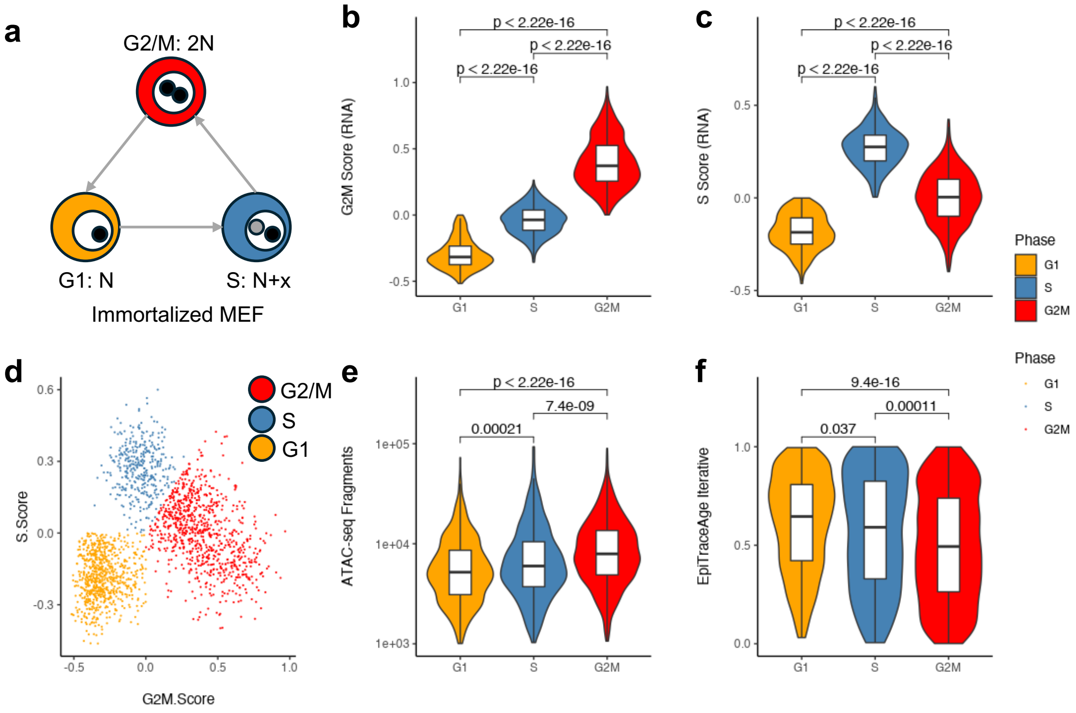


**Supplementary Figure 7. Cell cycle effect on EpiTrace age on immortalized mouse MEF cells.**

**(a)** The immortalized mouse MEF cells were subjected to SHARE-seq. **(b)** G2/M-phase-specific gene expression score and **(c)** S-phase-specific gene expression score of classified G1-, S- and G2/M-phase cells. **(d)** Classification of single cells according to the S-score and G2/M-score. **(e)** ATAC fragments from single cells in G1, S, and G2/M phase. **(f)** EpiTrace age of G1-, S-, and G2/M-phase cells.

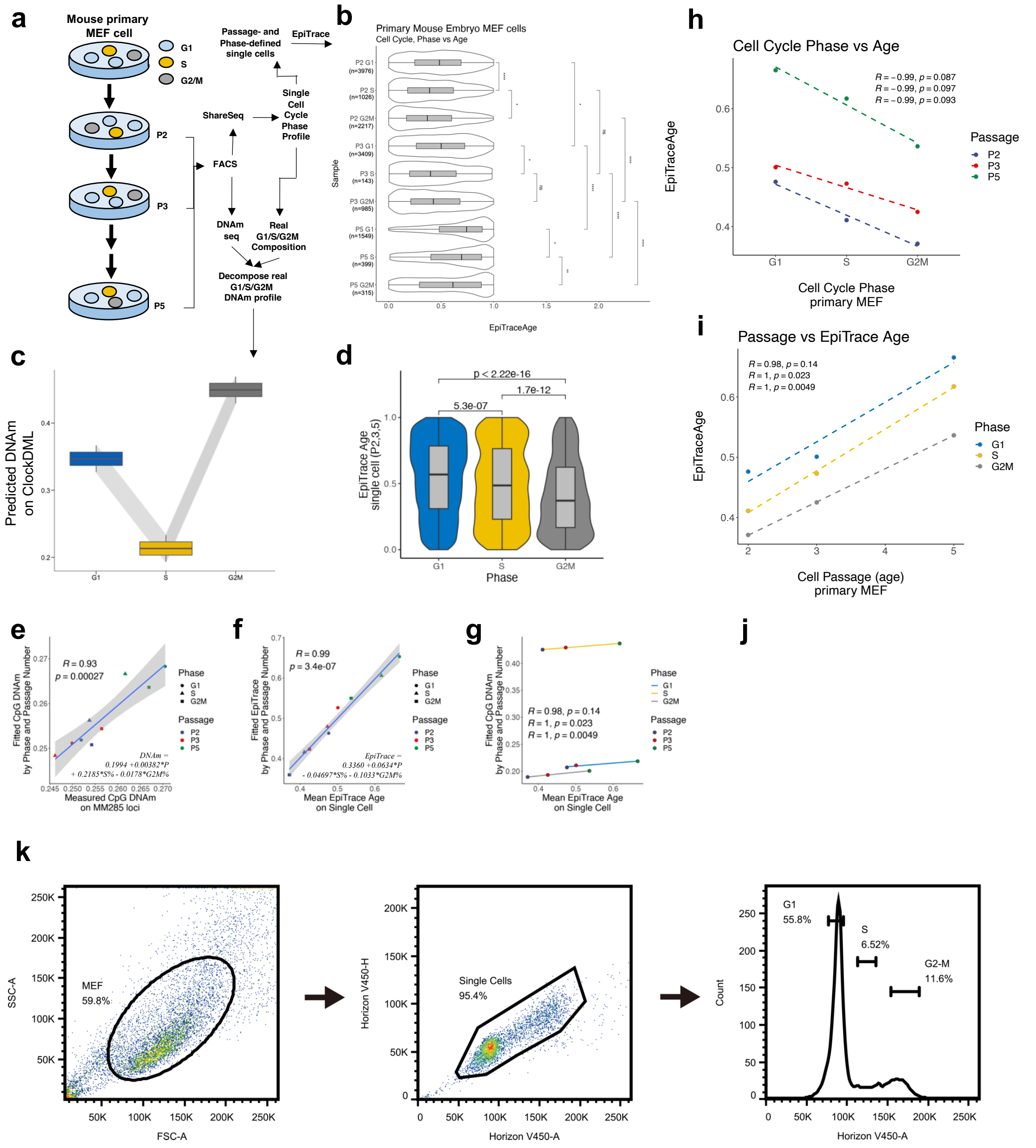


**Supplementary Figure 8. The epigenetic modification cycle during cell replication underlies EpiTrace age dynamics.**

**(a)** Schematic of the experiment: primary mouse embryonic fibroblasts (MEFs) were obtained from a single den of mouse embryos and cultured in vitro. At a defined passage (2, 3, 5), half of the cells underwent FACS sorting to split them to enrich for cells in G1, S or G2/M phase according to DNA content. Half of the sorted cells were then subjected to multiomics profiling with SHARE-seq. The other half of the cells underwent anchored PCR to amplify mouse-specific ClockDML loci (MM285) for bisulfite sequencing. The real G1/S/G2M percentage of sorted samples was determined by the scRNA profile. Single cells with a scRNA expression profile matching the FACS profile were included in the EpiTrace analysis. **(b)** EpiTrace age inferred from the single cell ATAC sequencing data, split by passage (P2/3/5) and phase. **(c)** Predicted DNAm on ClockDML by cell cycle phase (also see **e**). Active DNA methylation occurred during G1->S phase, and new “naïve” DNA was incorporated into the genome during the S->G2/M transition. **(d)** New “naïve” histone inclusion into the genome during G1->S->G2/M progression resulted in a decrease in EpiTrace age. **(e)** Linear fitting of ClockDML DNAm against the sample cell cycle profile (S-phase percentage, G2/M-phase percentage) and mitosis number indicates that both cell mitosis age and cell cycle profile contributed to ClockDML DNAm linearly. **(f)** Linear fitting of the mean EpiTrace age of single cells against the cell cycle profile (S-phase percentage, G2/M-phase percentage) and mitosis number indicates that both cell mitosis age and cell cycle profile contributed to EpiTrace age linearly. **(g)** Correlation between EpiTrace age and ClockDML DNAm, split by cell cycle phase (G1/S/G2M). **(h)** For each mitosis, the EpiTrace age of the same phase incrementally increased. **(i)** Correlation of EpiTrace age against mitosis generation, split by each cell cycle phase. **(j)** Schematic model of the cell cycle and the respective timing of DNA/histone exchange. **(k)** Example FACS sorting result. MEFs are first gated for whole cells and cell debris (FSC and SSC), then for single cells (Horizon V450-A and Horizon V450-H), and lastly for cells in G1, S, or G2/M phase according to DNA content. P-values: Student’s t-test for grouped comparison and Pearson’s R for linear regression.

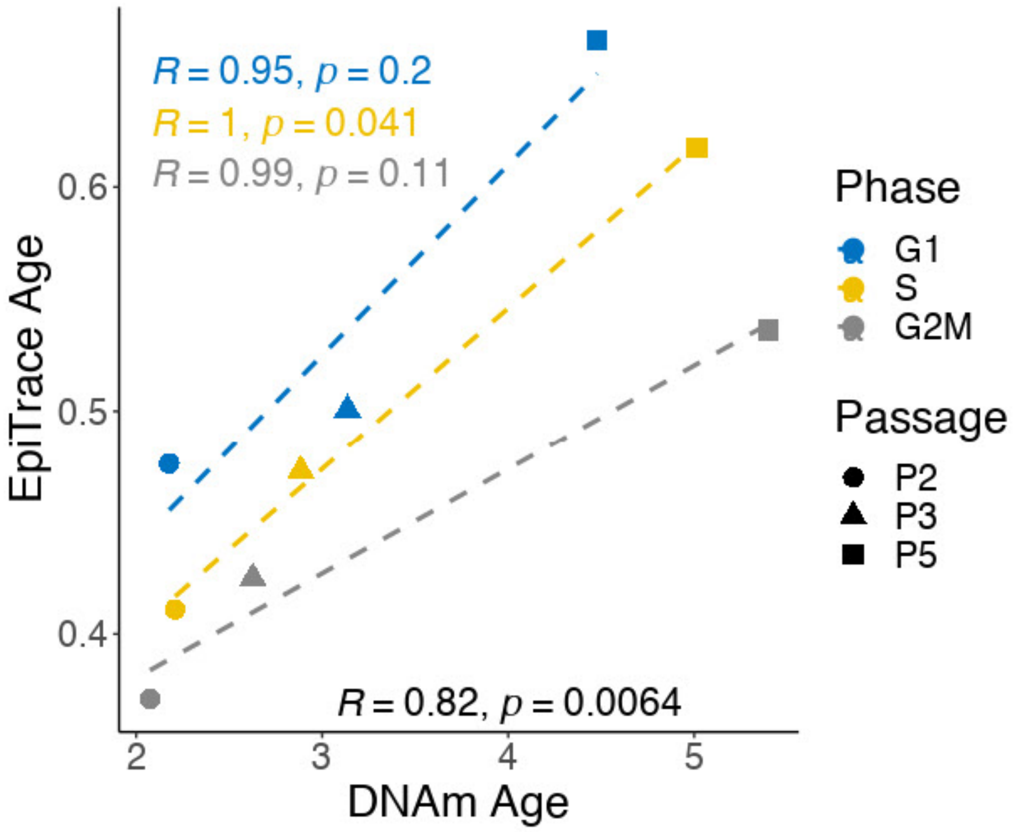


**Supplementary Figure 9. Correlation between DNAm-estimated age and EpiTrace-estimated age of the same pMEF samples.**

Cells from G1, G2/M, and S-phase are individually grouped for linear regression. Overall linear regression across the cell cycle phase is given in the lower right corner.

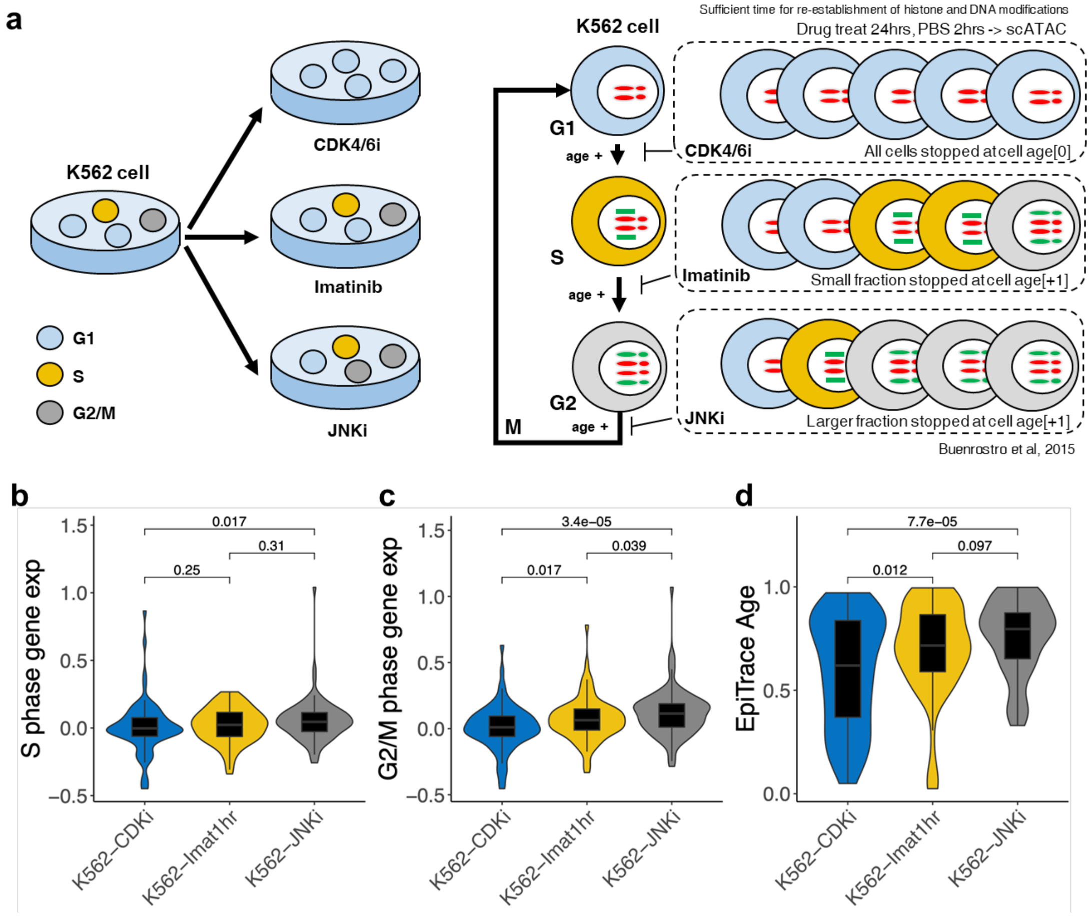


**Supplementary Figure 10. Prolonged blockage of cell cycle progression overcomes the effect of naïve histone inclusion on EpiTrace age.**

**(a)** Schematic of the pharmacological cell cycle inhibition experiment[5] in the CLL cell line (K562). Cells that undergo cycling are stopped at G1 (CDK4/6 inhibitor, CDKi), S (imatinib, Imat1 hr), or G2 (JNK inhibitor, JNKi) before scATAC sequencing. The potential increase in cell age is shown in the figure. **(b)** scATAC inferred S phase cell cycle gene expression of the treated cell groups. P-values are from the Wilcoxon test. **(c)** scATAC inferred G2/M phase cell cycle gene expression of the treated cell groups. P-values are from the Wilcoxon test. **(d)** EpiTrace age of treated cell groups. P-values are from the Wilcoxon test.

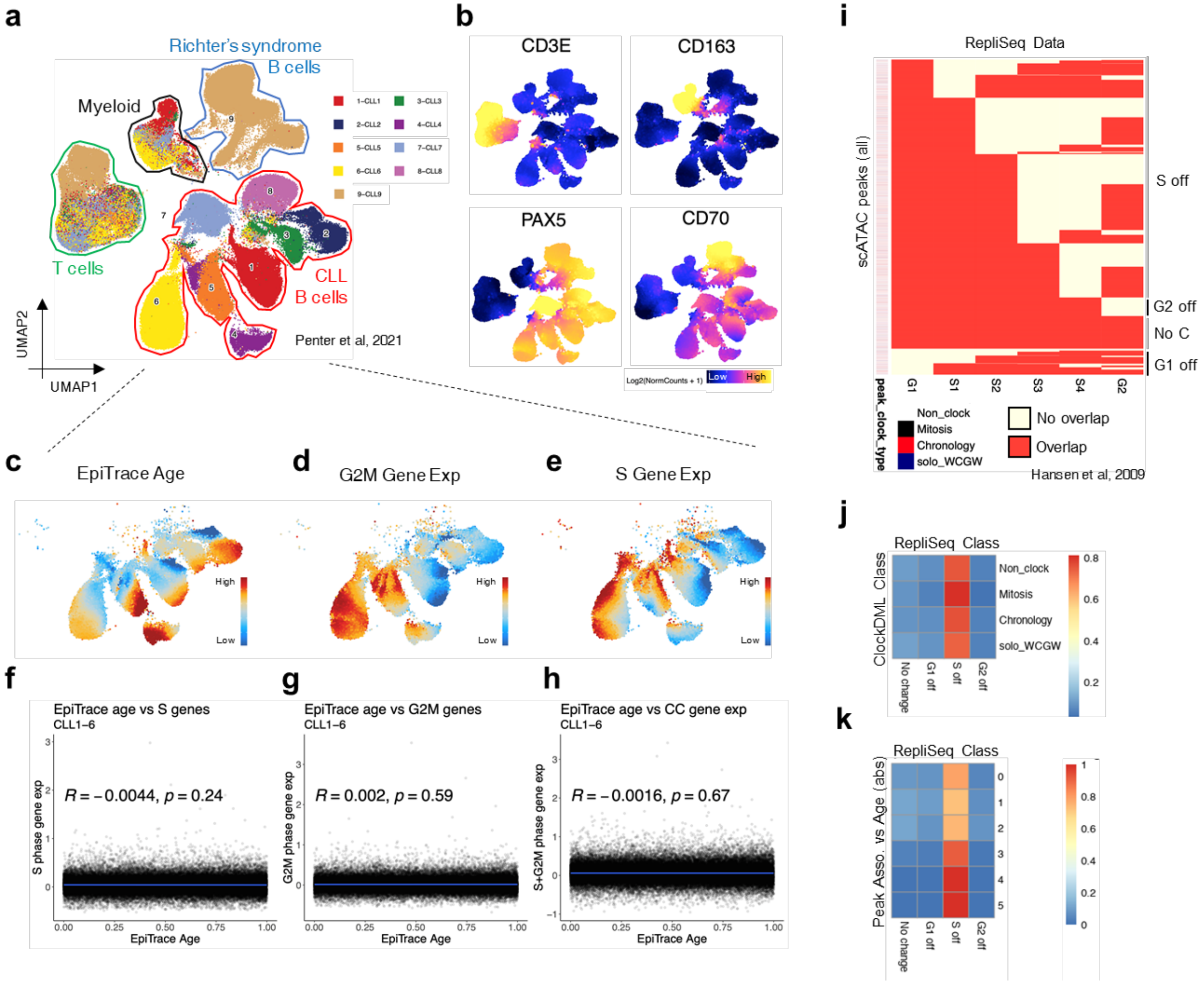


**Supplementary Figure 11. The global shift in EpiTrace age is not mainly driven by the cell cycle in human cells in vivo.**

**(a)** UMAP of scATAC cells in vivo collected blood cells from 9 CLL patients[6], labeled by their cell types and colored by donor. **(b)** Marker gene expression for each cell type (T: CD3E; Myeloid: CD163; B: PAX5; Richter's syndrome transformed B: CD70). **(c)** EpiTrace age of B cells from CLL1-CLL6. **(d)** G2/M phase cell cycle gene expression. **(e)** S phase cell cycle gene expression. **(f)** No correlation between EpiTrace age and S phase gene expression. **(g)** No correlation between EpiTrace age and G2/M phase gene expression. **(h)** No correlation between EpiTrace age and the expression of all cell cycle-related genes. **(i)** scATAC peaks overlapping with Repli-seq identified DNAseI-hypersensitive regions at each cell cycle phase[7], annotated by whether they are with a ClockDML (peak clock type: left annotation). Peaks are classified by their behavior in Repli-seq as "G1 off", "S off", "G2 off", or "No change". **(j)** The prevalence of ClockDML overlapping with each Repli-seq domain class. There was no significant enrichment of peaks with ClockDML (in contrast to other peaks) with any Repli-seq domain class. **(k)** The prevalence of scATAC peaks, grouped by their association with EpiTrace age, overlapping with each Repli-seq domain class. There was no significant difference in the EpiTrace age association distribution across the Repli-seq domain class (the majority of peaks were S-off).

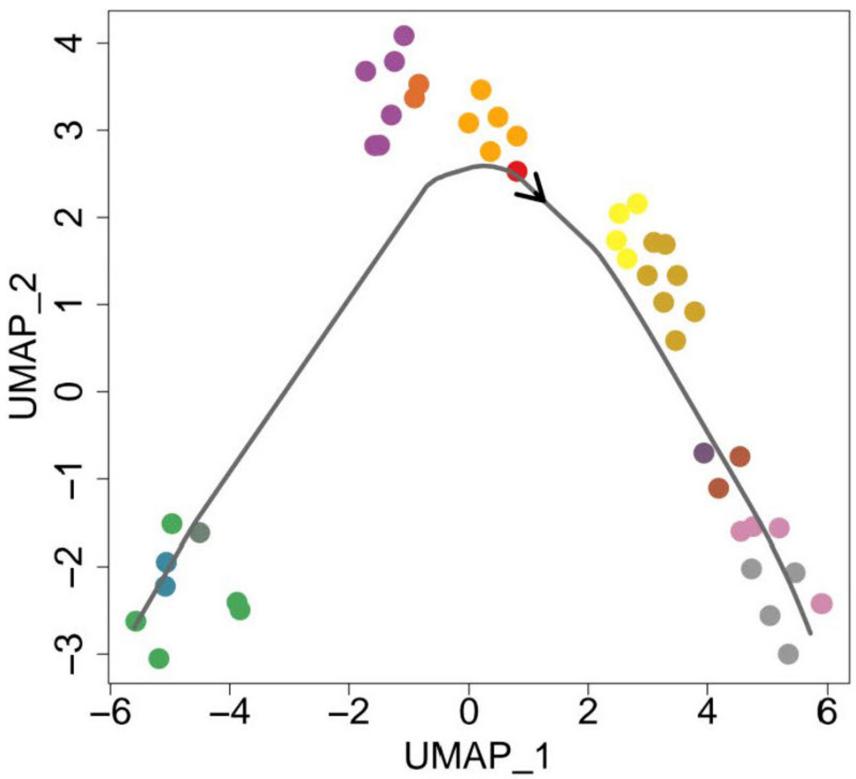


**Supplementary Figure 12. Raw data of slingshot-based prediction of human embryonic development trajectory.**

Color codes of the cells are similar to those in Figure 1e.

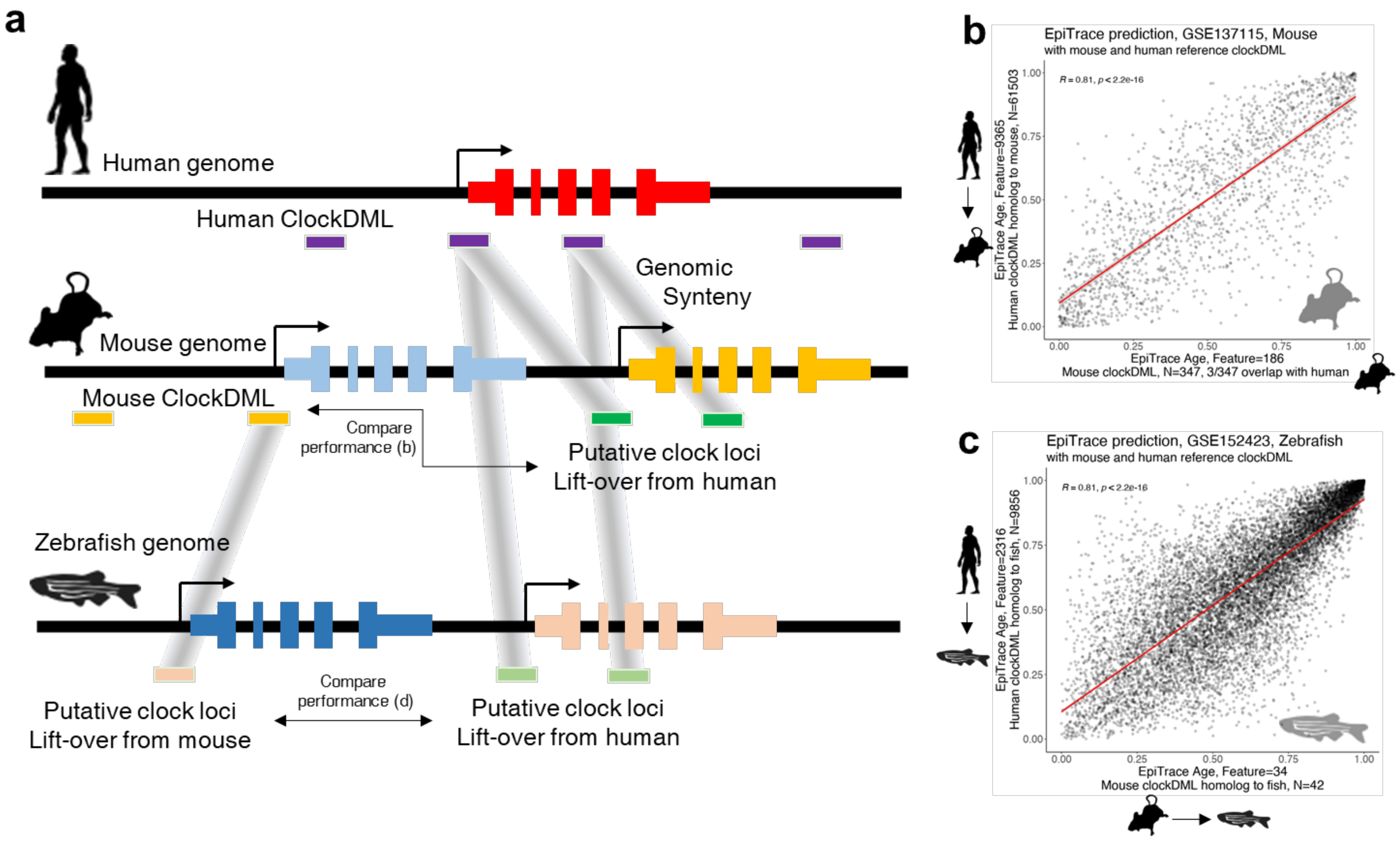


**Supplementary Figure 13. Additional validation of the accuracy of cross-species inference.**

**(a)** Schematic of the definition of putative clock genomic regions in the nonhuman vertebrate genome. Clock genomic loci could be either defined by genome-wide DNA methylation analysis (mouse) or genomic synteny (human mapped to mouse, and human/mouse mapped to zebrafish). **(b)** Comparing EpiTrace prediction of mouse single cell age with mouse ClockDML (x) or lift-over human ClockDML (y) as reference clock loci. R and P-values are from Pearson's correlation. **(c)** Comparing EpiTrace prediction of zebrafish single cell age with lift-over mouse (x) or human (y) ClockDML as reference clock loci. R and P-values are from Pearson's correlation. Mouse data: GSE137115. Zebrafish data: GSE152423.

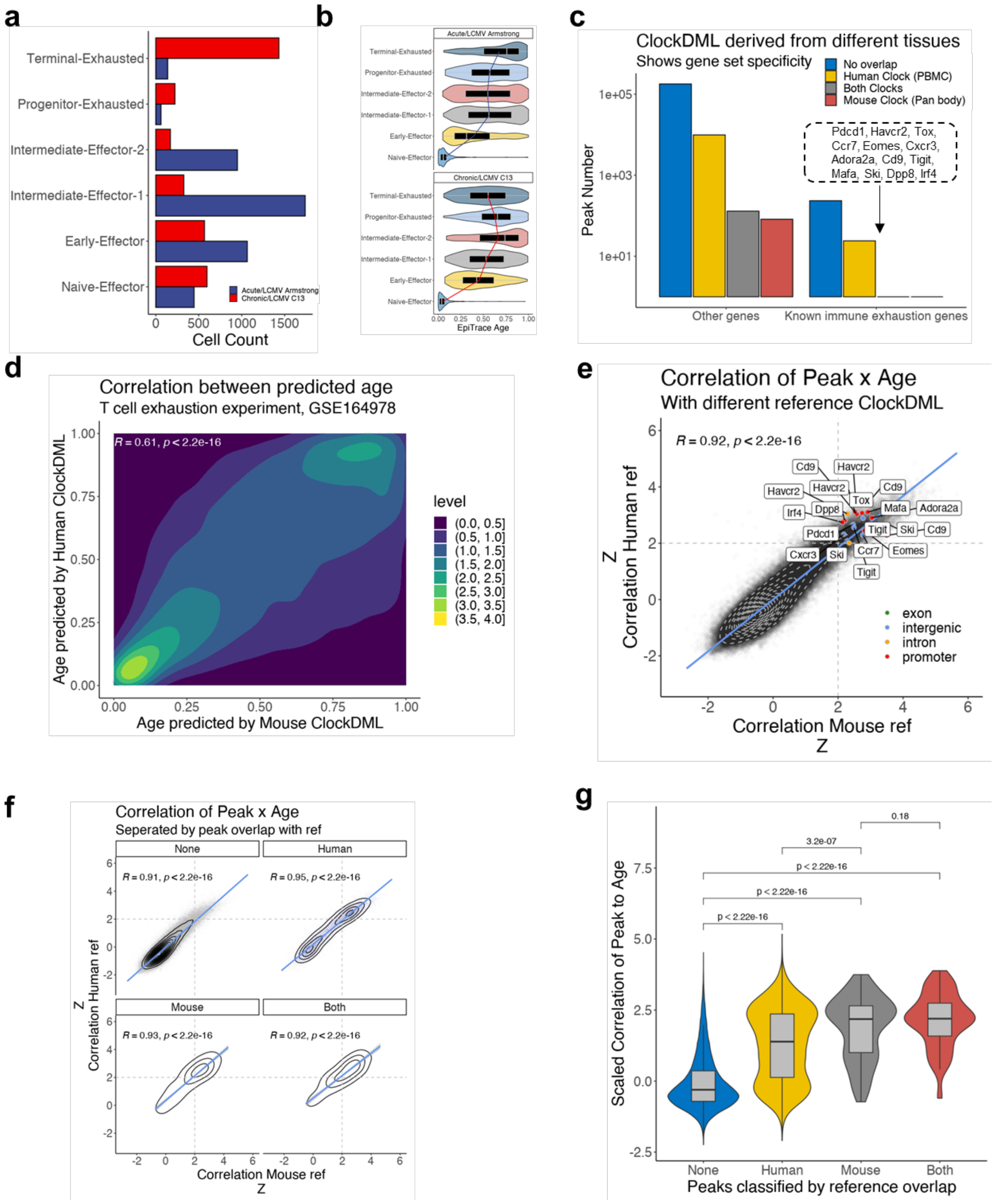


**Supplementary Figure 14. Prediction of mouse blood cell mitosis age using human blood-cell-derived ClockDML sites and mouse non-blood-derived ClockDML sets.**

**(a)** LCMV-responding T cells in the mouse cell dataset GSE164978, including naïve, intermediate, and terminally differentiated effector CD8+ T cells as well as dysfunctional progenitor and terminally exhausted CD8+ T cells. **(b)** EpiTrace age predicted cell age in acute or chronic LCMV strain infection. **(c)** scATAC peaks classified by overlap with ClockDML on non-immune exhaustion (other genes) or known/expected immune exhaustion genes (names shown in the figure). The Panbody mouse ClockDML set does not include any overlap on these candidate immune exhaustion genes. **(d)** Correlation between single cell ages predicted by human PBMC-trained ClockDML or mouse panbody ClockDML. **(e)** Association between scATAC peaks and cell age by two predictions. Labeled points are enhancer peaks for known immune exhaustion genes. **(f)** Association of peaks to EpiTrace ages (predicted by human or mouse ClockDML), classified by their overlap to ClockDML sets. **(g)** Association of peaks, classified by their overlap to ClockDML, to EpiTrace age (predicted by human ClockDML). ClockDML harboring a genomic region shows a greater association with cell age. Correlation: Pearson's. Grouped comparison significance was determined by t-test.

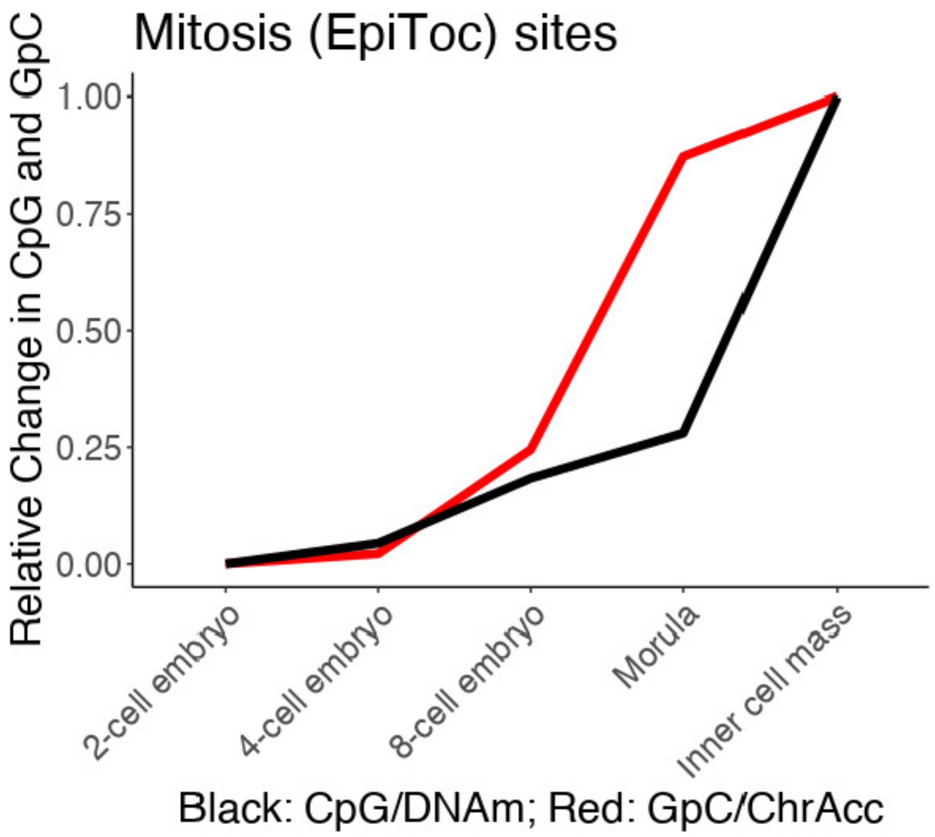


**Supplementary Figure 15. A chromatin accessibility shift precedes a DNAm shift in human embryonic development.**

Single cell scCOOL-seq of human embryo development was performed. Aggregated GpC methylation changes (red) and CpG methylation changes (black) are shown for EpiTOC ClockDML. Data: GSE100272.

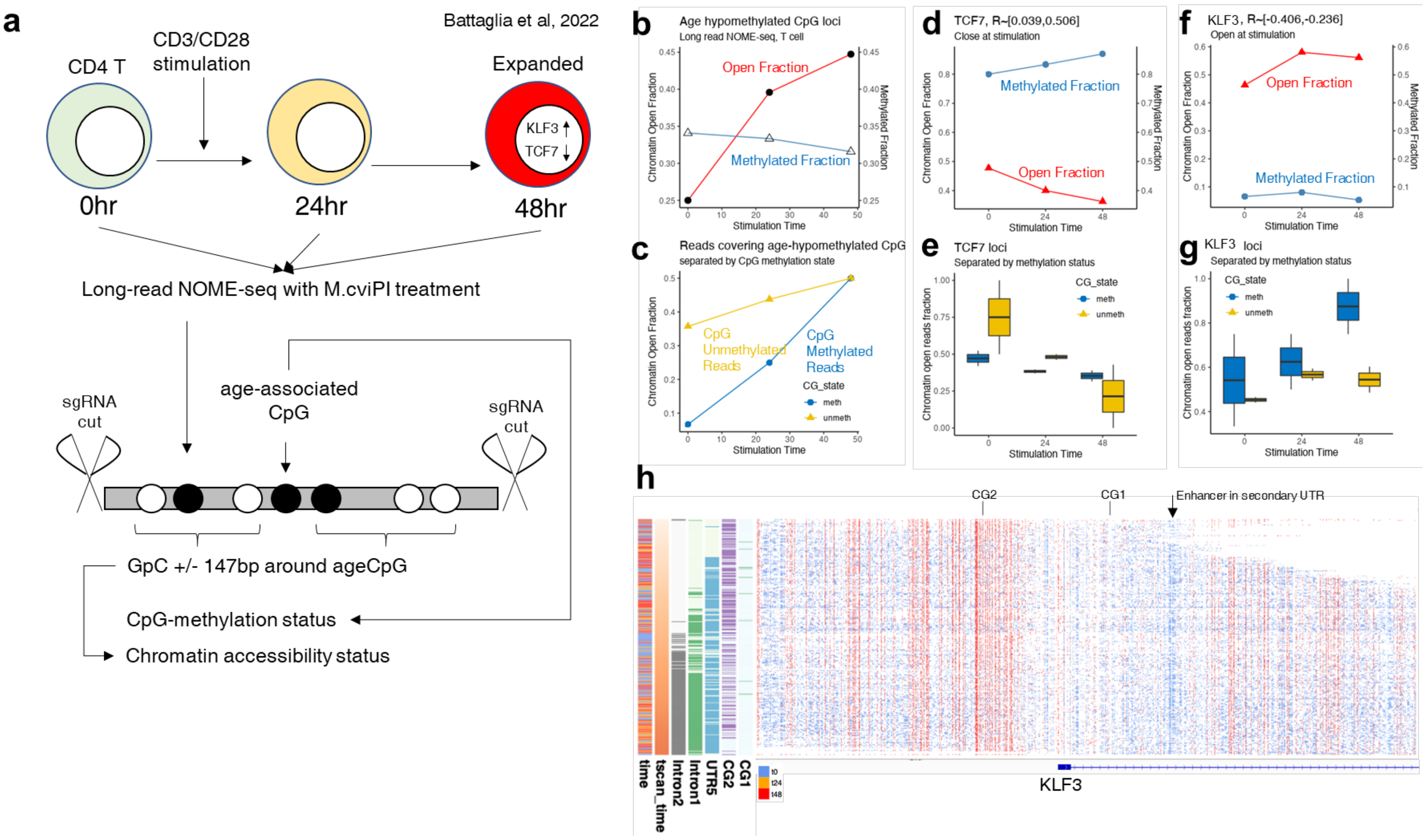
a
Battaglia et al, 2022
CD4 T
CD3/CD28
stimulation
Expanded
KLF3
TCF7
0hr
24hr
48hr
Long-read NOME-seq with M.cviPI treatment
sgRNA
cut
age-associated
CpG
sgRNA
cut
GpC +/- 147bp around ageCpG
CpG-methylation status
Chromatin accessibility status
b
Age hypomethylated CpG loci
Long read NOME-seq, T cell
Open Fraction
Methylated Fraction
Chromatin Open Fraction
Methylated Fraction
Stimulation Time
c
Reads covering age-hypomethylated CpG
separated by CpG methylation state
CpG
Unmethylated
Reads
CpG
Methylated
Reads
CG_state
meth
unmeth
d
TCF7, R~[0.039,0.506]
Close at stimulation
e
TCF7 loci
Separated by methylation status
Chromatin open reads fraction
f
KLF3, R~[-0.406,-0.236]
Open at stimulation
g
KLF3 loci
Separated by methylation status
h
CG2
CG1
Enhancer in secondary UTR
time
tscan_time
Intron2
Intron1
UTR5
CG2
CG1
t0
t24
t48
KLF3

**Supplementary Figure 16. Age-associated CpG methylation changes are secondary to chromatin accessibility shifts.**

**(a)** Schematic diagram of the analysis. Naïve human CD4 T cells were expanded with CD3/CD28 stimulation, and cells were collected at the indicated time points. Genomic DNA was treated with M.cviPI to methylate accessible (open) GpC loci and subjected to selective nanopore sequencing facilitated by targeted Cas9 digestion. The methylation state of each cytosine on the read was determined. Chromatin openness around the CpG loci was called by +/- 147 bp GpC methylation level >= 20%. Since the targeted sequenced reads encompassed few DMLs with an age association > 0.7 (ClockDML), we analyzed all CpGs that showed an association with age (classified as hypomethylated or hypermethylated). **(b)** Reads encompassing age-hypomethylated CpG sites opened quickly in response to stimulation, while the CpG methylation fraction changed little. **(c)** Accessibility of reads encompassing age-hypomethylated CpG increased in response to stimulation regardless of the specific CpG methylation state. **(d)** Methylated read fraction and open read fraction at the TCF7 locus. CpG correlation with age are all positive in this loci (shown in title). **(e)** Mean accessibility of reads with unmethylated or methylated CpG (grouped by CpG site) on TCF7. The accessibility of unmethylated reads decreased without having undergone CpG methylation. **(f)** Methylated read fraction and open read fraction at the KLF3 locus. CpG correlation with age are all negative in this loci (shown in title). **(g)** Mean accessibility of reads with unmethylated or methylated CpG (grouped by CpG site) on KLF3. The accessibility of methylated reads increased without having undergone CpG demethylation. **(h)** Details of long-read nanoNOME reads arranged by pseudotime (tscan_time) and sampling time (time). Reads are shown as horizontal lines with GpC methylation (blue) or CpG methylation (red). CG2/CG1 and activated enhancer. are shown. Together, these results indicate that chromatin accessibility shifts in response to artificial stimulation precede DNA methylation changes in age-associated CpG loci. Data: GSE183760.

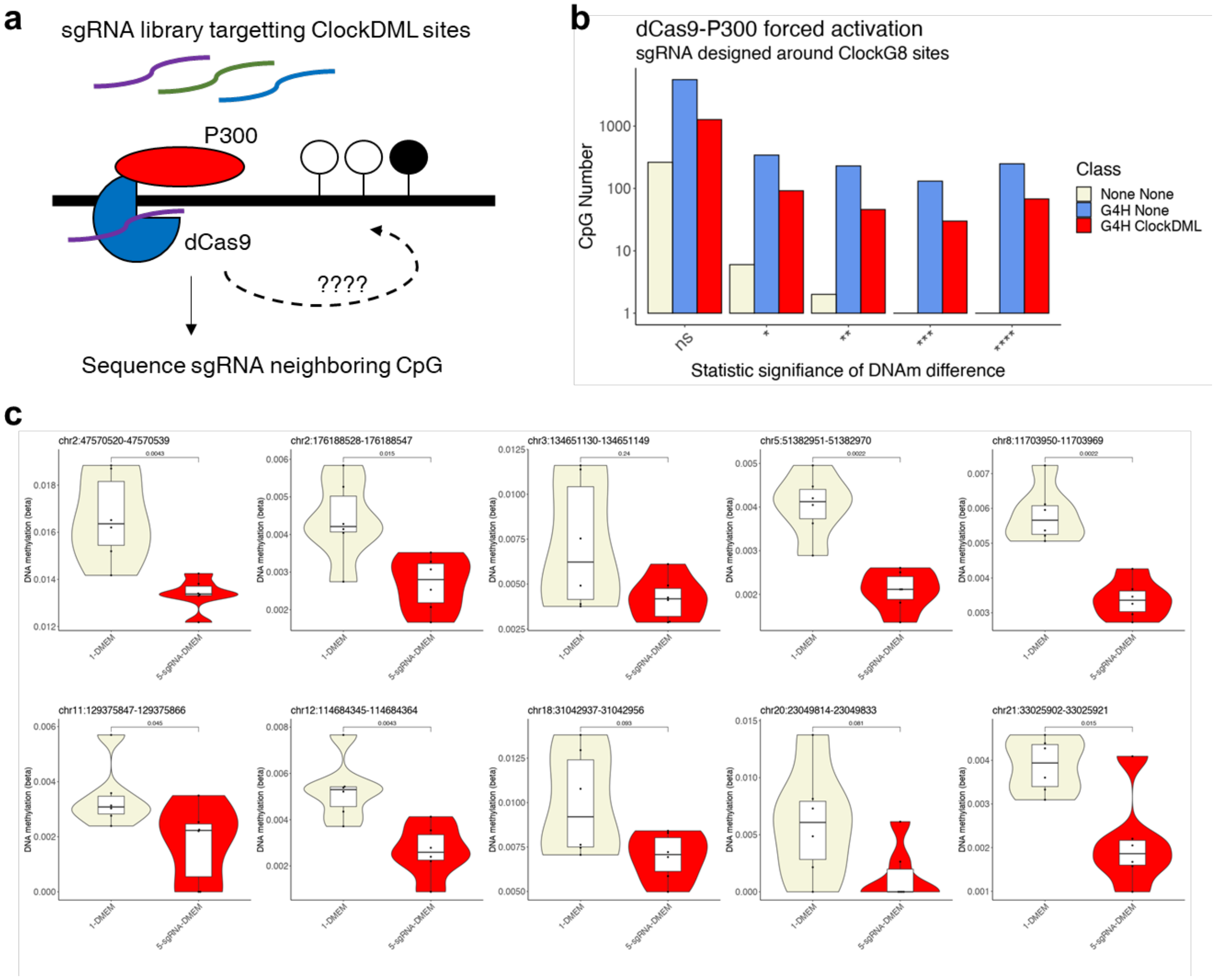
a
sgRNA library targetting ClockDML sites
P300
dCas9
????
Sequence sgRNA neighboring CpG
b
dCas9-P300 forced activation
sgRNA designed around ClockG8 sites
CpG Number
1000
100
10
1
ns
*
**
***
****
Statistic signifiance of DNAm difference
Class
None None
G4H None
G4H ClockDML
c
chr2:47570520-47570539
chr2:176188528-176188547
chr3:134651130-134651149
chr5:51382951-51382970
chr8:11703950-11703969
chr11:129375847-129375866
chr12:114684345-114684364
chr18:31042937-31042956
chr20:23049814-23049833
chr21:33025902-33025921
DNA methylation (beta)
1-DMEM
5-sgRNA-DMEM

**Supplementary Figure 17. Gain of chromatin accessibility around ClockDML G8-class loci drives local DNA hypomethylation.**

**(a)** Schematic of the experiment. **(b)** Number of CpG loci with different statistical significance levels in DNAm between targeted and control cell samples. **(c)** Example CpG loci that show DNA demethylation in the targeted cell samples compared to the control. P-value: t-test. P-values are BH adjusted.

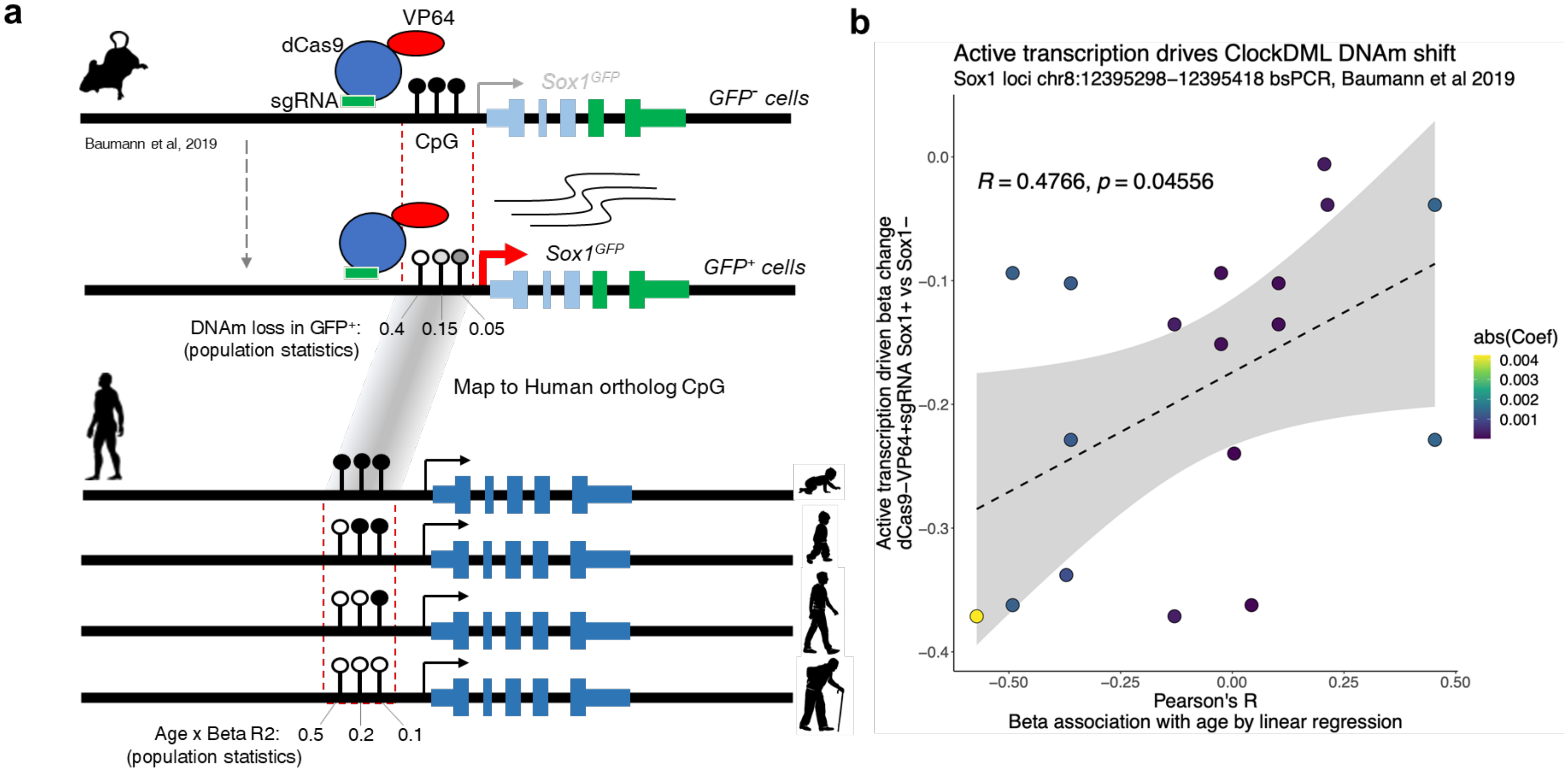


**Supplementary Figure 18. Binding of transcriptional activators in TSS promotes spontaneous DNA methylation loss on clock-like differentially methylated loci.**

**(a)** Schematic diagram of the analysis. Mouse Sox1$^{GFP}$ cells expressing sgRNA-guided dCas9-VP64 binding to the mouse Sox1 promoter (dCas9 cells) were FACS-sorted by GFP expression and sequenced for CpG methylation on the Sox1 promoter. While dCas9-target binding was strong, the prevalence of GFP$^{+}$ cells was low. bsPCR sequencing showed that GFP expression is accompanied by sporadic DNA demethylation. Actively demethylating the loci by dCas9-TET coexpression enhanced Sox1 expression[8]. We mapped the mouse loci to human orthologous CpG loci and calculated the Pearson's correlation between CpG methylation level and human age. The spontaneous demethylation (driven by occupation of DNA by transactivator proteins, possibly by blocking DNMT interaction with the target DNA) tendency in mice was then compared to the age-methylation correlation on these loci. **(b)** Correlation between age-methylation correlation (Pearson's R, x-axis) and transcription-driven demethylation (Sox1$^{+}$ - Sox1$^{-}$, y-axis) for sequenced CpG in the Sox1 promoter. CpGs are colored by their absolute age-methylation correlation coefficient. R and P-values are from Pearson's test. Together, the data indicate that transactivator binding promotes spontaneous ClockDML DNAm shift.

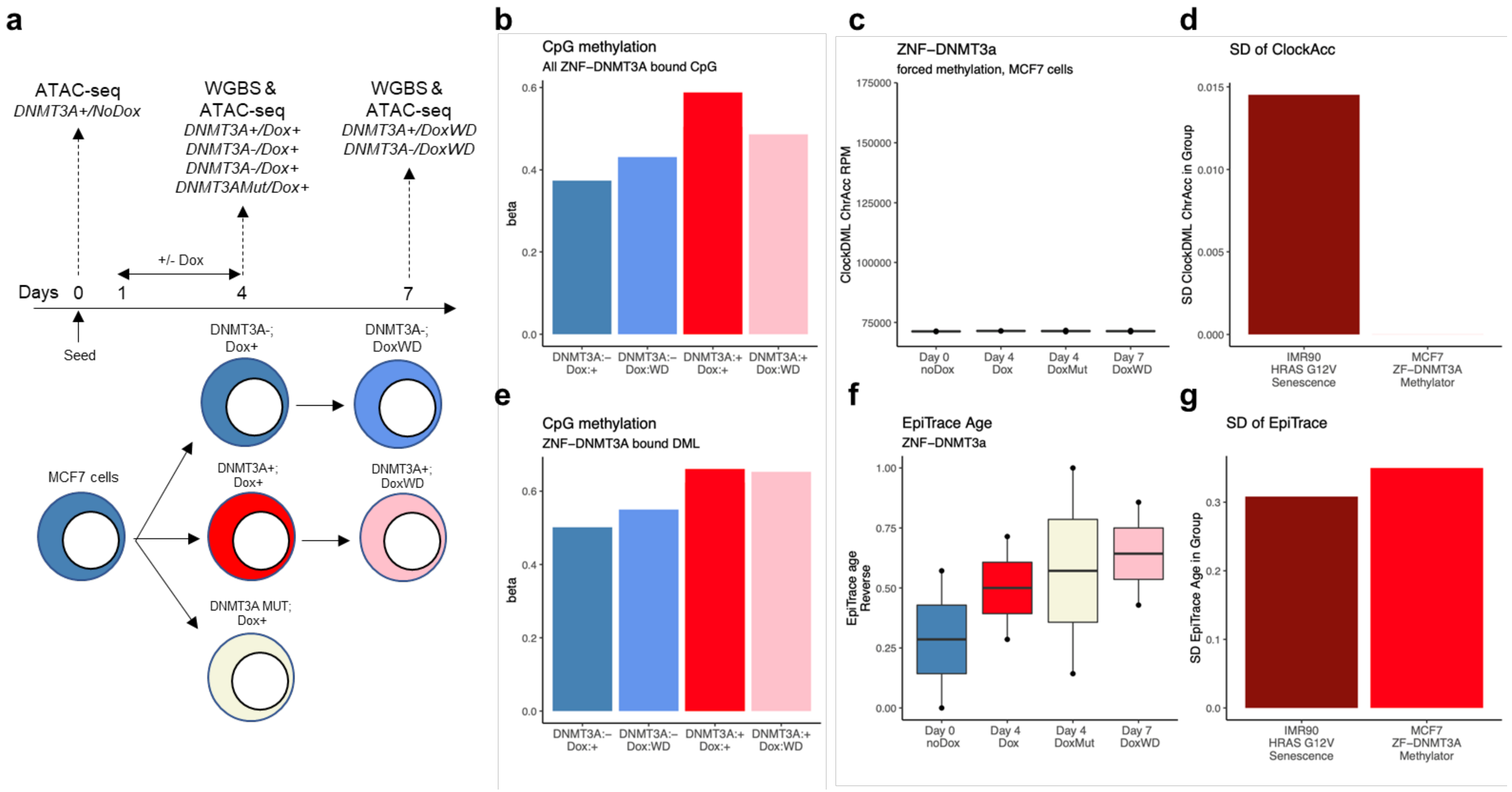


**Supplementary Figure 19. Forced methylation on ClockDML is irreversible but does not lead to EpiTrace age change.**

**(a)** Schematic of the ZNF-DNMT3A methylator expression experiment. Temporary expression of ZNF-DNMT3A or a ZNF-DNMT3A catalytic-dead mutant (DNMT3A-MUT) was induced by doxycycline. **(b)** DOX-induced zinc-finger-tethered DNMT3A (ZNF-DNMT3A) expression (DNMT3A+; Dox+) is accompanied by an increase in CpG methylation (beta) at its target sites compared to controls (DNMT3A-; Dox+/WD). This effect was quickly reversed by doxycycline withdrawal (DNMT3A+;Dox:WD). **(c)** On ClockDML sites, however, the methylation increase is irreversible (compare DNMT3A+; Dox+ with DNMT3A+; Dox:WD). **(d)** Minimal ClockDML ChrAcc change under ZNF-DNMT3A forced methylation (could be compared to ClockDML ChrAcc change during real cell senescence). **(e)** EpiTrace age of the ZNF-DNMT3A activation dataset is correlated to cell culture days (Days) but not expression of ZNF-DNMT3A (Dox) or ZNF-DNMT3A enzyme-dead mutant (DoxMut). **(f)** Standard deviation of ClockDML ChrAcc in the senescence or methylator datasets. **(g)** Standard deviation of EpiTrace age in the senescence or methylator dataset. Data: GSE103590 (senescence) and GSE102395 (methylator).

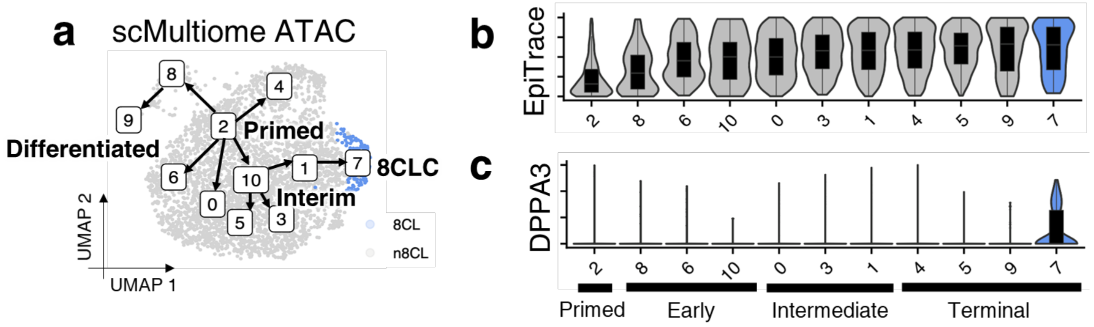


**Supplementary Figure 20. Details of the 8-cell-like induction scMultiome ATAC dataset.**

**(a)** scMultiome ATAC UMAP projection; cell clusters are labeled. Trajectories between clusters are drawn. The 8-cell-like (8CL) cluster is colored blue. **(b)** EpiTrace age prediction of single cells in each cluster, ordered by mean EpiTrace age. **(c)** DPPA3 enhancer activation in each cluster. Clusters are labeled “Primed” (ground-zero), “early”, “intermediate” and “terminal”, according to their position.

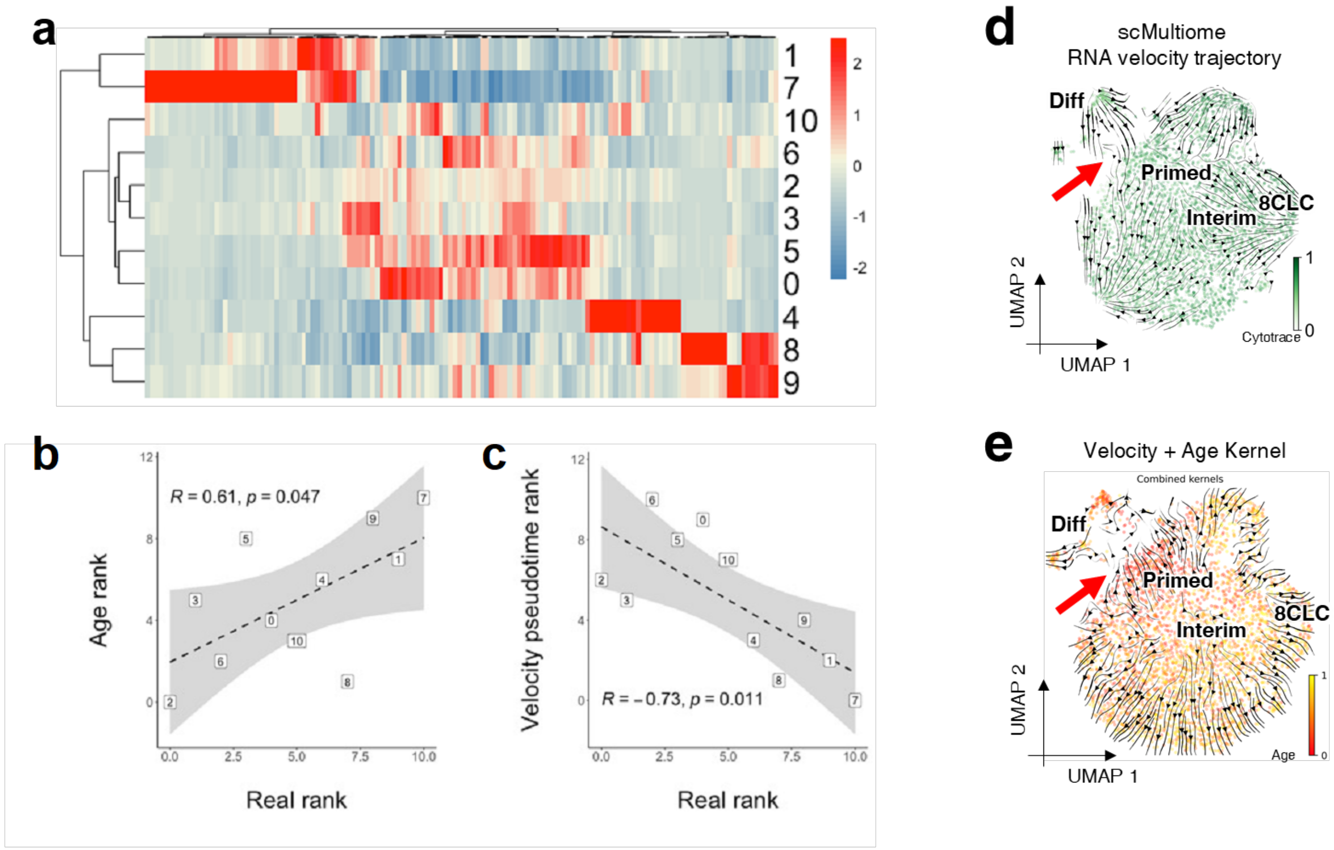


**Supplementary Figure 21. Comparison of RNA velocity-based prediction (PAGA) and EpiTrace prediction of the cell evolution trajectory in the 8CLC induction dataset.**

**(a)** The single cell clusters are hierarchically arranged based on the RNA expression of 4CL and 8CLC marker genes[9] and extracellular matrix marker genes enriched in Clusters 8 and 9, serving as the "real rank" in (b) and (c). The scaled RNA expression level is shown on the right. **(b)**

Correlation of RNA expression hierarchy (x-axis) and cluster hierarchy ranked by EpiTrace age (y-axis). **(c)** Correlation of RNA expression hierarchy (x-axis) and cluster hierarchy ranked by scVelo pseudotime (y-axis). **(d)** Streamline projection of RNA velocity on UMAP for the single cells. Highlighted is a trajectory from differentiated cells toward primed cells (the starting population). **(e)** Streamline projection of Velocity+Age kernel prediction of cell evolution velocity (CellRank) of the same cells, showing the primed cell evolving toward differentiated cells. Correlation statistics (R and P-value): Pearson's.

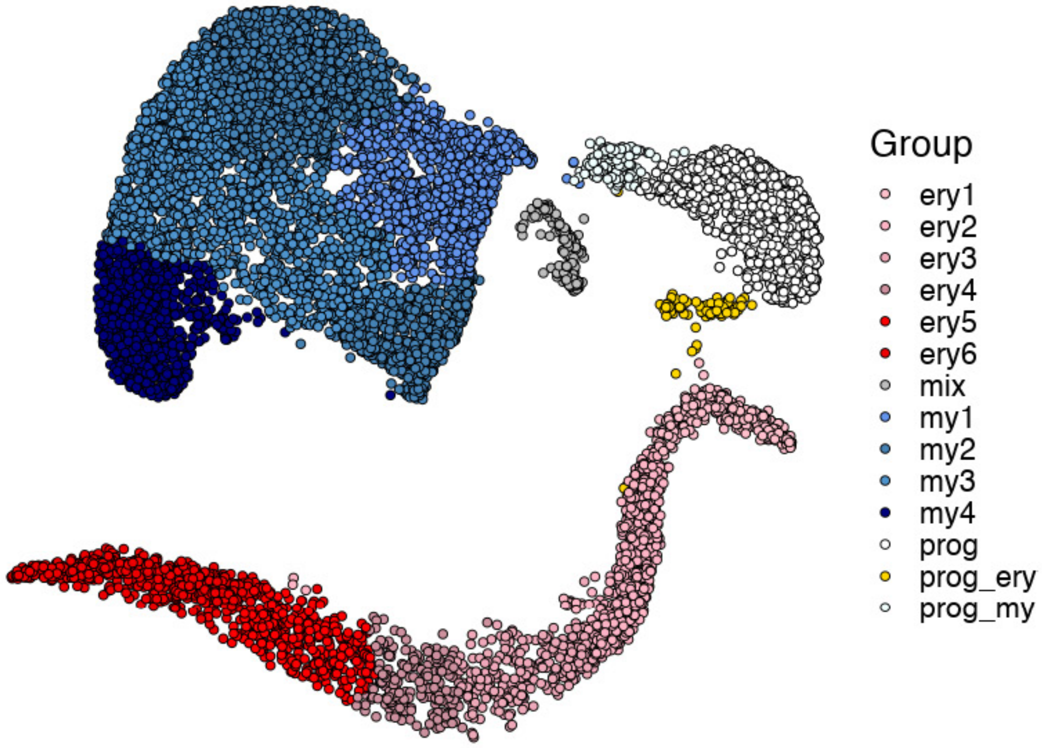


**Supplementary Figure 22. Cell type annotations for the in vitro CD34+ HSC expansion/differentiation experiment.**

Prog: progenitor cells; prog_ery: erythroid lineage progenitor cells; prog_my: myeloid lineage progenitor cells; mix: mixture lineage cells; my1-my4: myeloid cells at different stages of development; ery1-ery6: erythroid lineage cells at different stages of development.

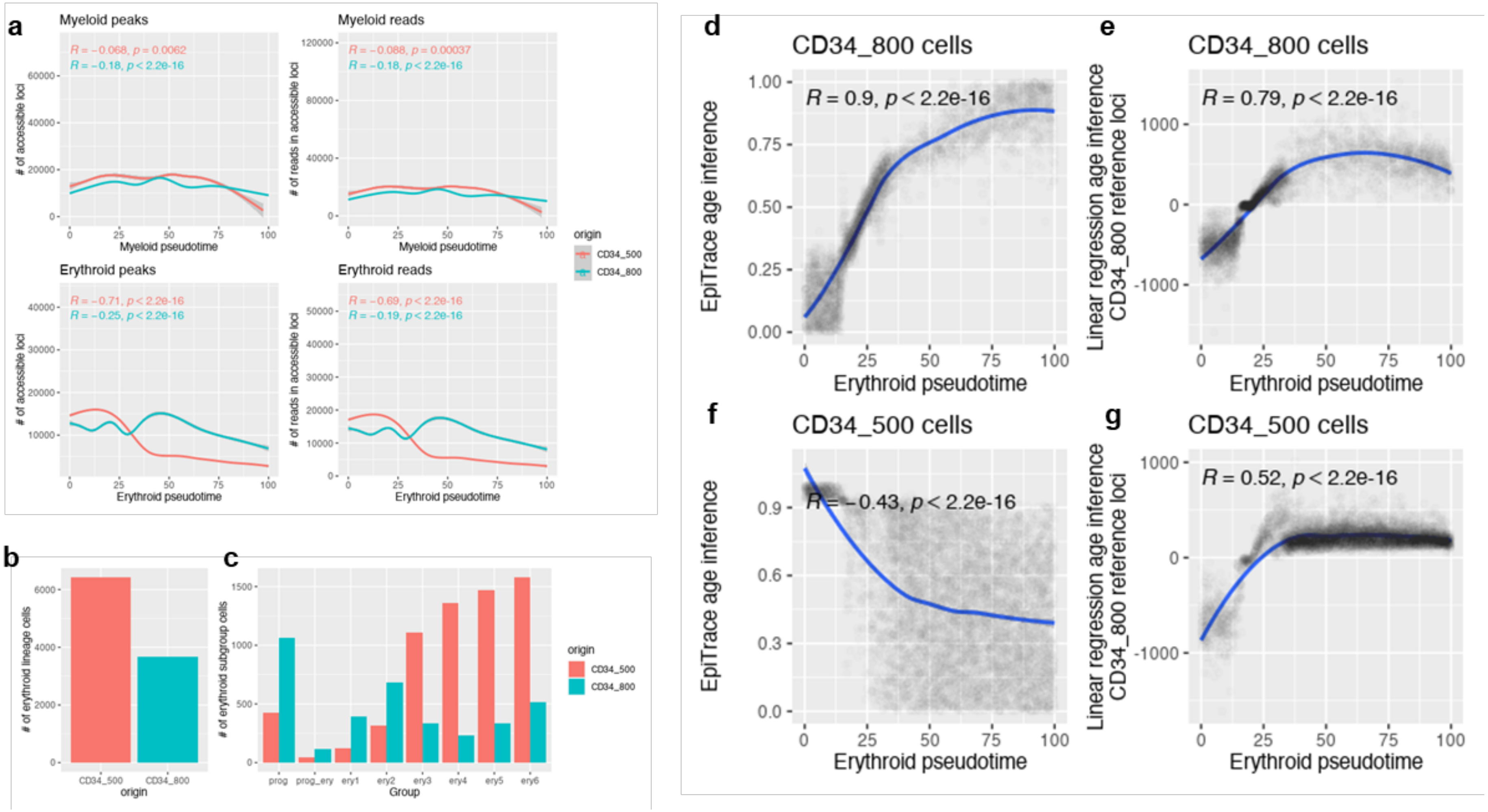


**Supplementary Figure 23. Cell composition bias could lead to inference error in EpiTrace analysis.**

(**a**) Distribution of single cell ATAC peaks or reads-in-peaks (y-axis) for myeloid and erythroid cells with respect to developmental pseudotime (x-axis). **(b)** Erythroid lineage cell number from CD34_500 and CD34_800 experiments. **(c)** Single cell numbers in each cluster from the CD34_500 and CD34_800 experiments. **(d)** Correlation between pseudotime (x-axis) and EpiTrace age (y-axis) for CD34_800 cells. **(e)** Correlation between pseudotime (x-axis) and "correlative loci" linear regression result (y-axis) for CD34_800 cells. **(f)** Correlation between pseudotime (x-axis) and EpiTrace age (y-axis) for CD34_500 cells. **(g)** Correlation between pseudotime (x-axis) and "correlative loci" linear regression result (y-axis) for CD34_500 cells, using the CD34_800 derived reference loci.

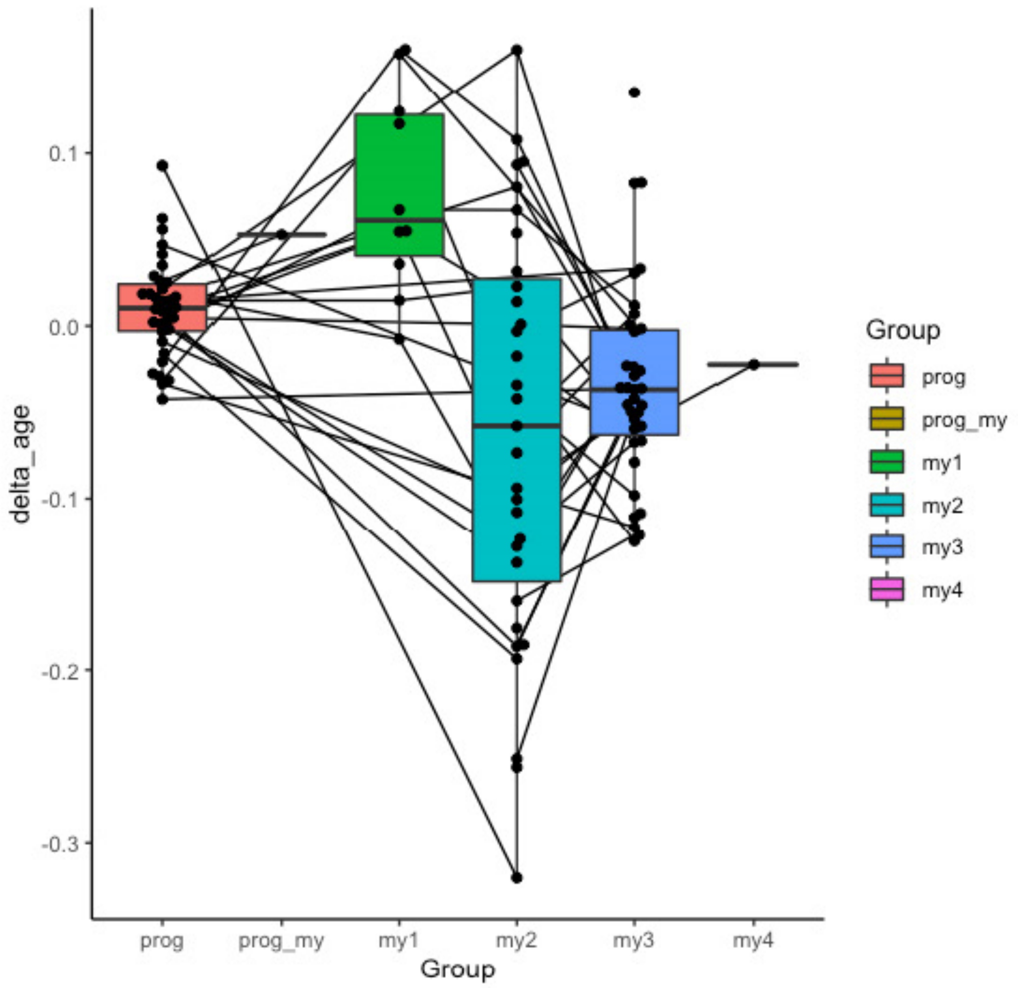


**Supplementary Figure 24. Age shift in myeloid cell clones following induced differentiation.**

Cells are grouped as in Supplementary Figure 22, and per-clone mean EpiTrace ages for cells in that particular differentiation state (cell cluster) are drawn. Lines denote similar clones.

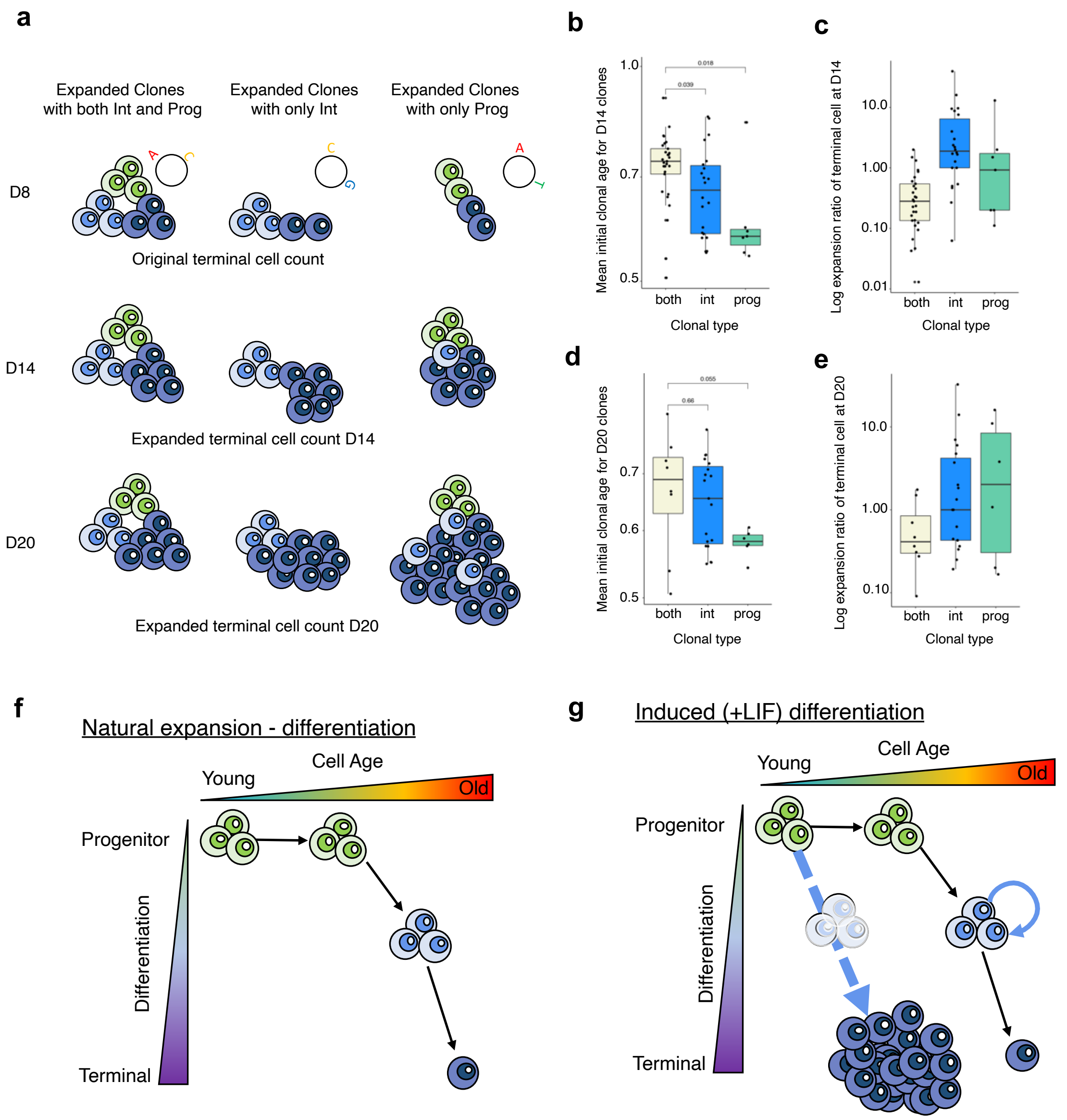


**Supplementary Figure 25. Analysis of the relationship between progenitor cell age and clonal expansion potential.**

**(a)** Schematic of the analysis. Single cells in the CD34+ HSC in vitro expansion/differentiation experiment were first grouped by their mtDNA mutation profile (examples as the mutations on "microcircles") into clones. Clones were further classified by whether progenitor and/or intermediately differentiated cells were present in the clone at the initial timepoint (Day 8/D 8). Clones were then tracked at subsequent timepoints (Day 14/D 14, Day 20/D 20) to measure the increase in terminally differentiated cells. **(b)** Mean initial clonal EpiTrace age for the clones analyzed on day 14. **(c)** Mean initial clonal EpiTrace age for the clones analyzed on day 20. **(d)** Log-expansion ratio of terminal cells in each clone on day 14. **(e)** Log-expansion ratio of terminal cells in each clone on day 20. **(f)** Scenario under natural expansion-differentiation. **(g)** Scenario under forced differentiation. Correlation statistics (R and P-value): Pearson's. Group statistics: t-test.

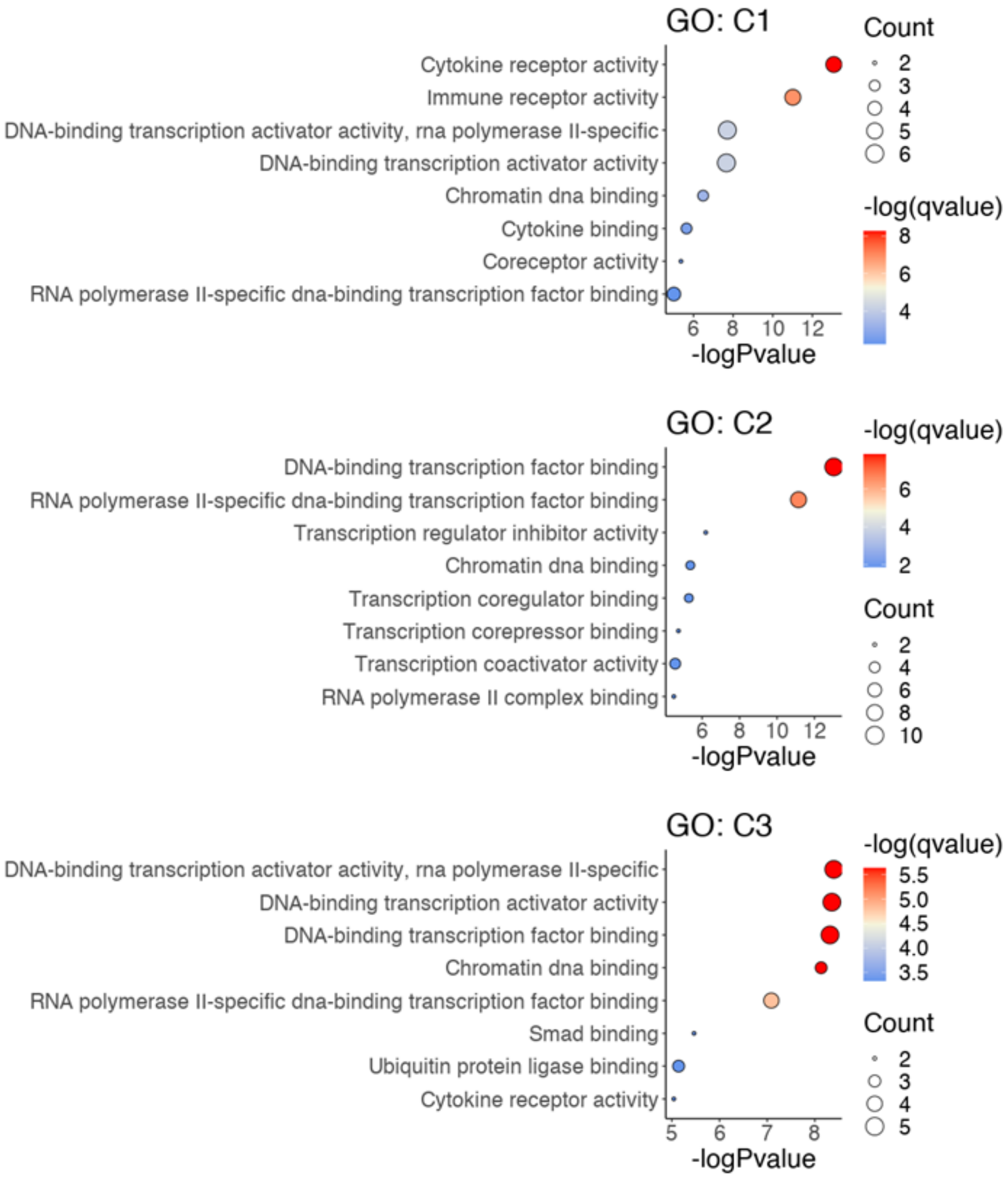


**Supplementary Figure 26. GO-term enrichment of differentially activated peak clusters in intratumoral cytotoxic T cells according to the classification in Fig. 4c.**

Only pathways with adjusted P values < 0.05 are shown. The C1-specific enriched pathway is highlighted by an arrow.

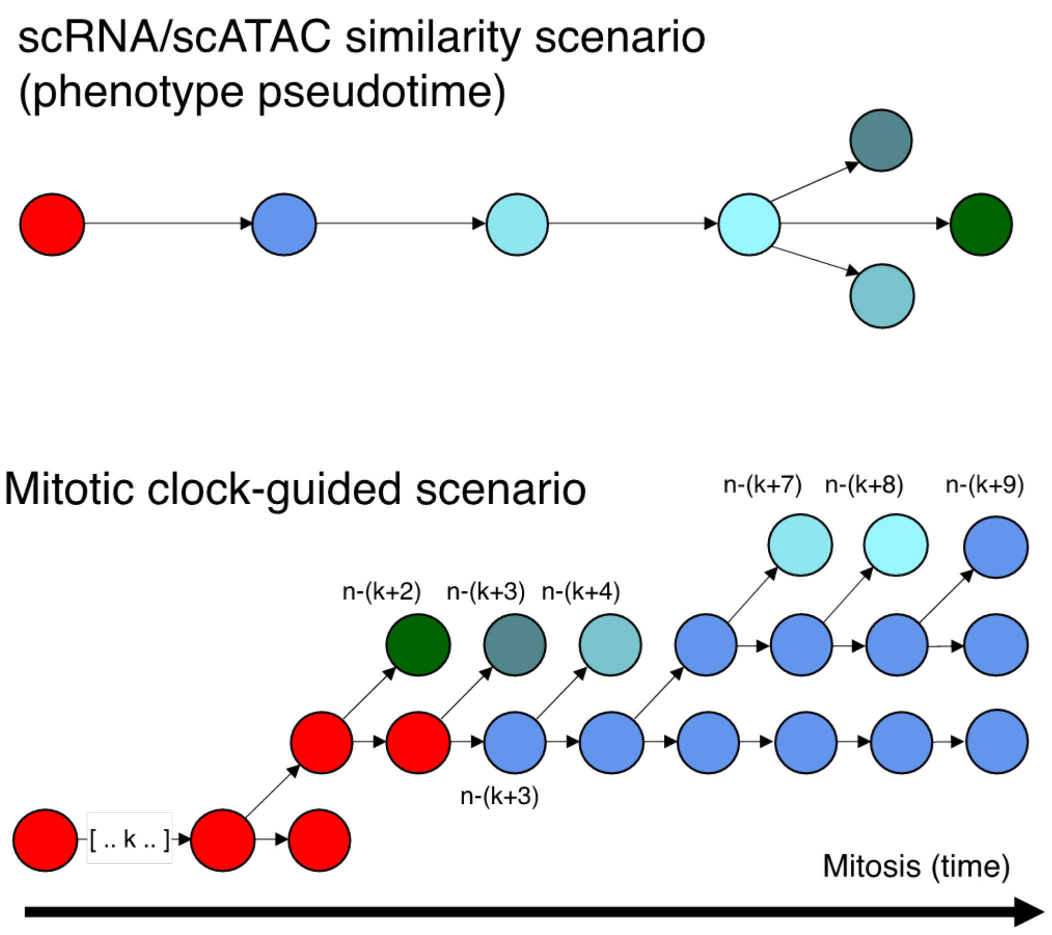


**Supplementary Figure 27. Glutaminergic neuron development model by scRNA/scATAC similarity or combined with mitotic clock (EpiTrace).**

The radial glia (RG) cells divide to differentiate into intermediate progenitor cells (IPCs), which further divide into subplate cells (first emerged, SP) or neurons (N). The earlier derived neurons undergo longer postmitotic development time and are more phenotypically mature. The relative cell division number of each cell is labeled for the mitotic-clock-guided scenario.

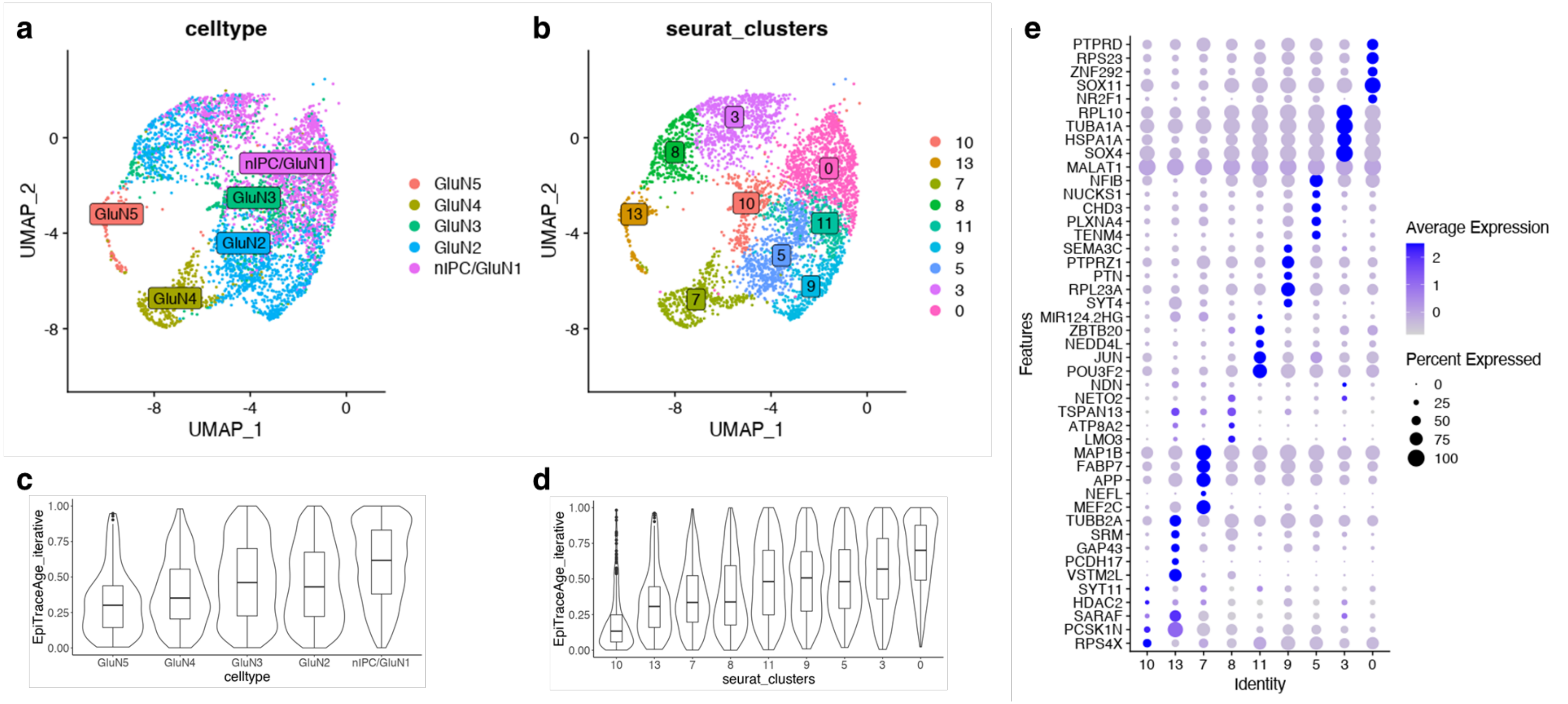


**Supplementary Figure 28. Postmitotic glutaminergic neuron maturation.**

**(a)** UMAP of GluN2, 3, 4, and 5 neurons. **(b)** UMAP of the same neurons clustered by gene expression. **(c)** EpiTrace age of cells in (a), classified as in the original publication by scATAC similarity. **(d)** EpiTrace age of cells in (b), classified by gene expression clusters. **(e)** Differentially expressed genes in each cell cluster, as in (b).

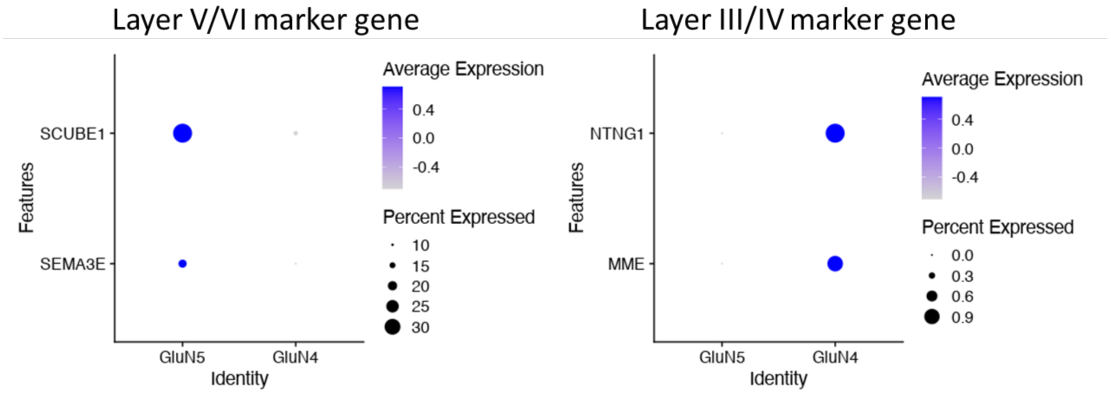


**Supplementary Figure 29. Correlation of single cell cluster age and layer-specific gene expression.**

Expression of Layer V/VI (Left) and Layer III/IV (Right) marker genes in the GluN5 and GluN4 clusters.

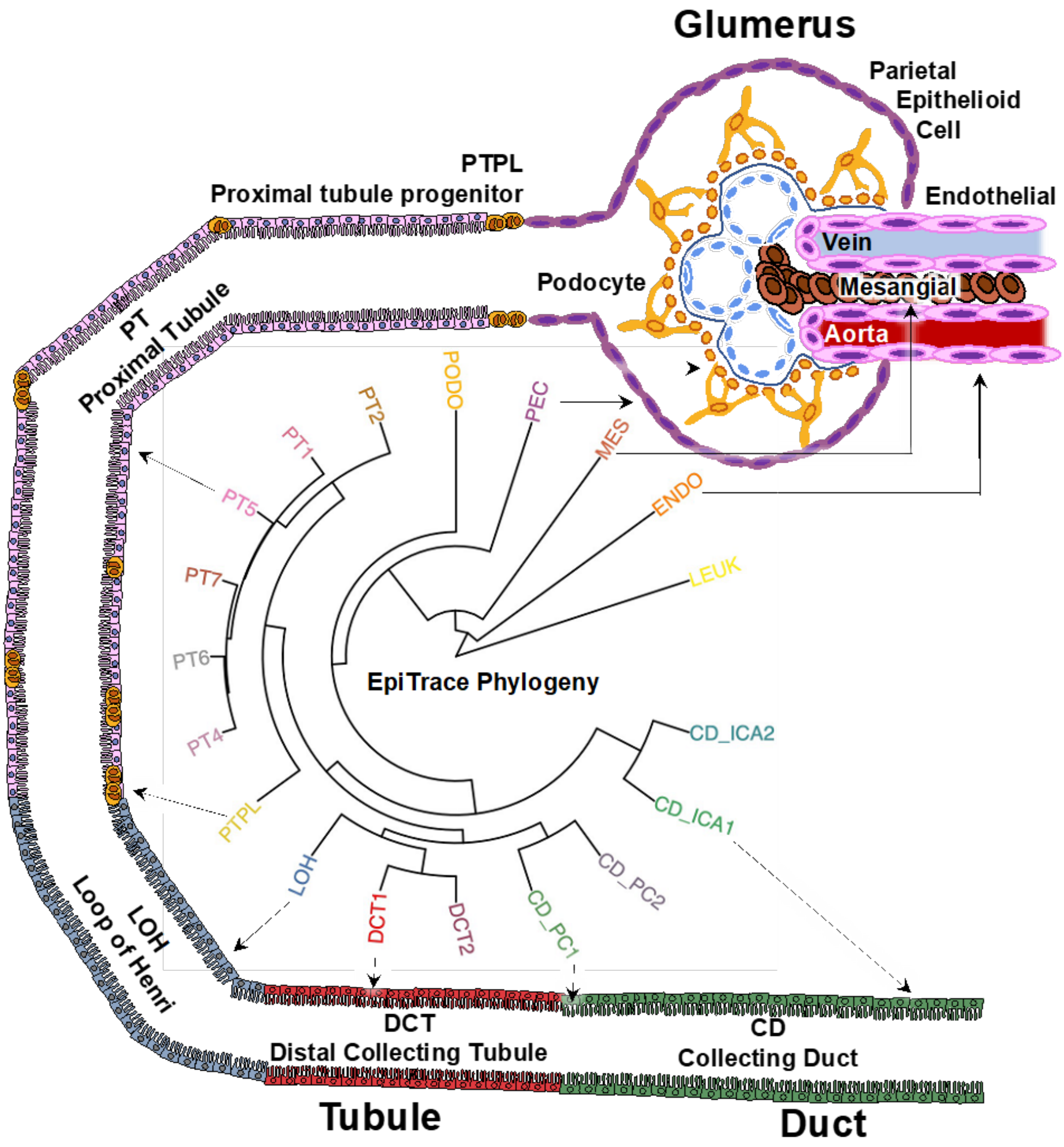


**Supplementary Figure 30. Additional schematic drawing showing the correlation between anatomical positioning and the EpiTrace phylogeny of kidney cells.**

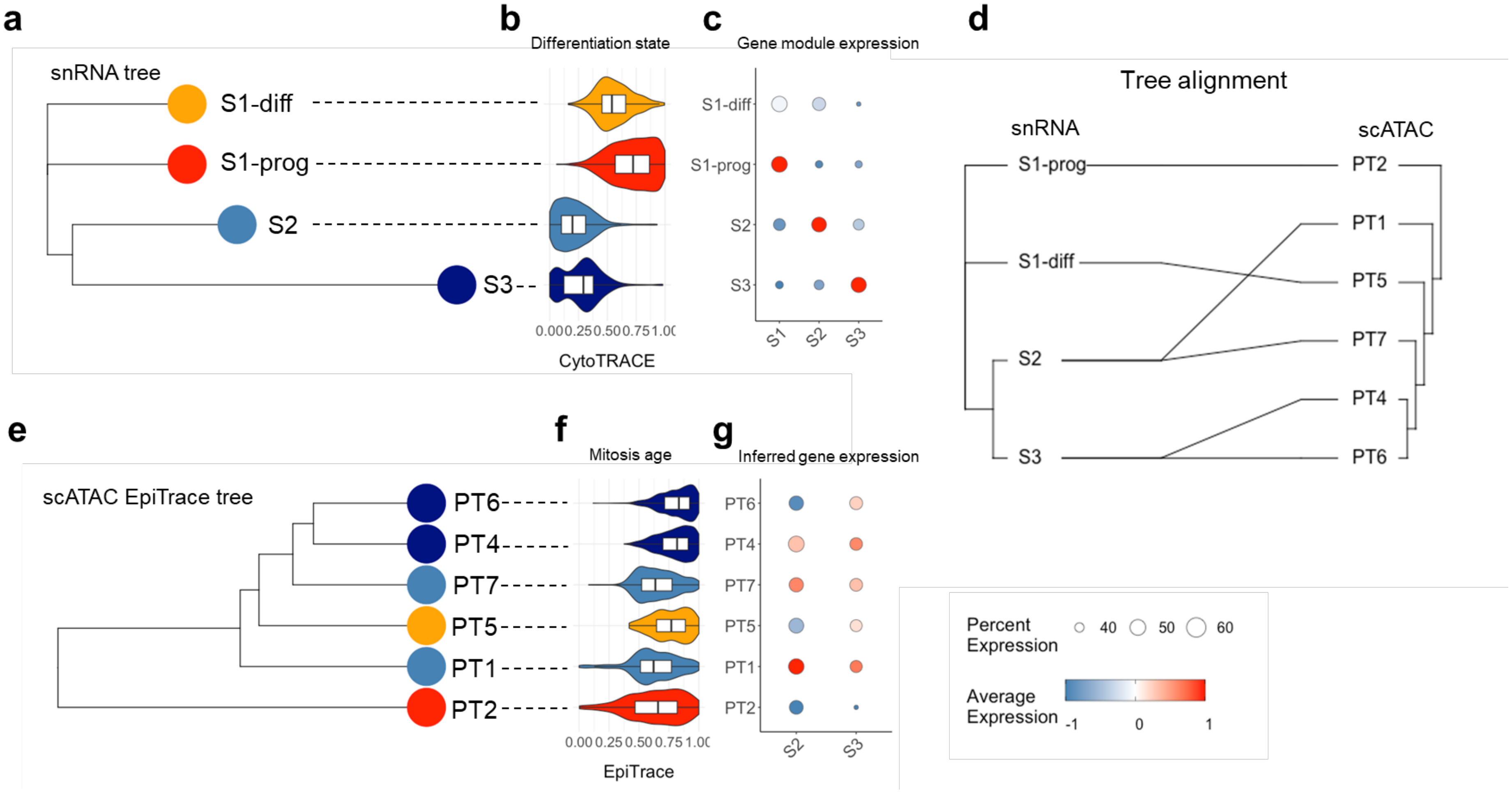


**Supplementary Figure 31. Validation of EpiTrace-inferred proximal tubule cell development phylogeny with snRNA data.**

**(a)** Phylogenetic tree built with snRNA expression data from human kidney PT cells, showing S1-prog, S1-diff, S2, and S3 clusters. **(b)** CytoTRACE score of snRNA PT cells. **(c)** Scaled expression of PT-cell-cluster gene modules in these snRNA cells. **(d)** Matching between phylogenetic trees of snRNA and scATAC results. **(e)** Phylogenetic tree built with scATAC using EpiTrace. **(f)** Mitosis age inferred by EpiTrace. **(g)** Scaled expression of PT cell cluster gene modules in these scATAC cells.

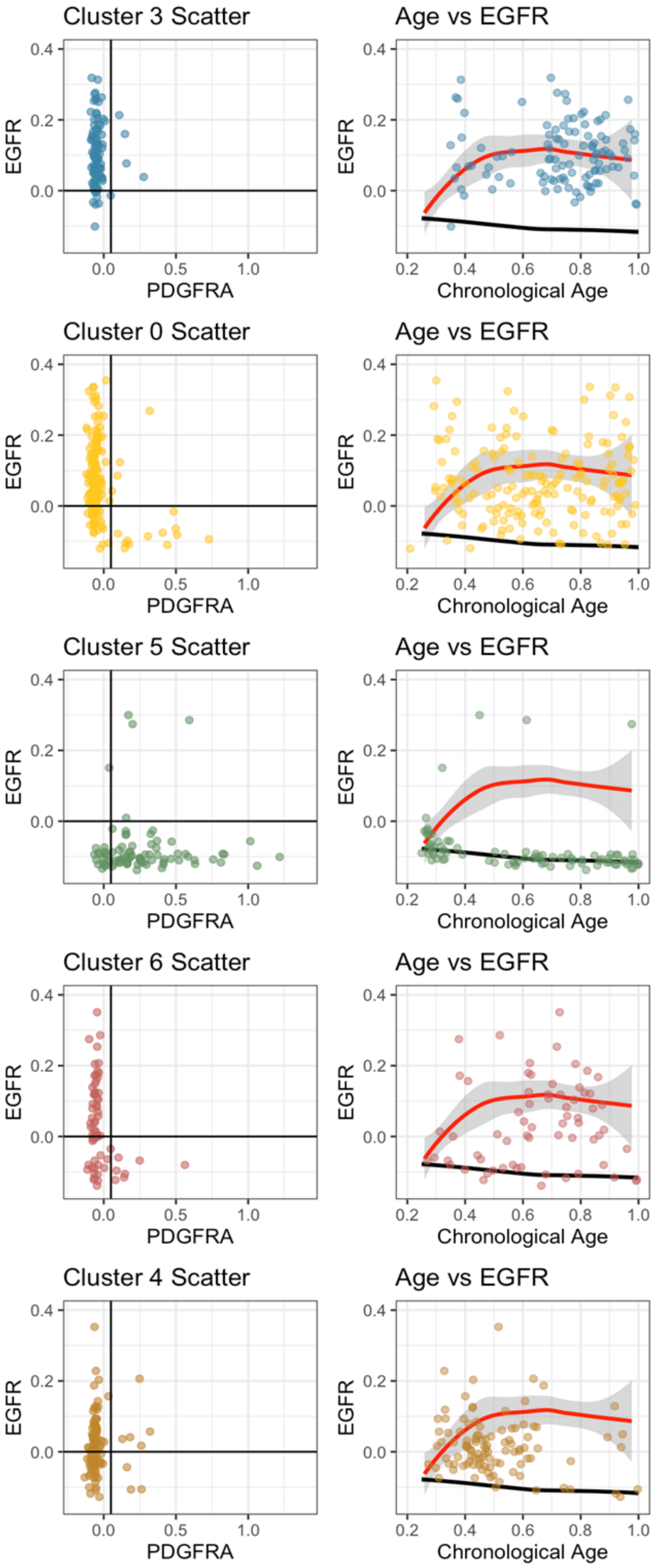
Cluster 3 Scatter
Age vs EGFR
Cluster 0 Scatter
Age vs EGFR
Cluster 5 Scatter
Age vs EGFR
Cluster 6 Scatter
Age vs EGFR
Cluster 4 Scatter
Age vs EGFR
EGFR
PDGFRA
Chronological Age
0.4
0.2
0.0
0.5
1.0
0.6
0.8

**Supplementary Figure 32. Branching evolution of EGFR+ and PDGFRA+ clusters from a double-positive EGFR+/PDGFRA+ cancer clone.**

Scatter plot (left) shows the EGFR and PDGFRA peak heights of each single cell within the designated cluster. Single cell EGFR peak height (y-axis) as a function of EpiTrace age (x-axis) is shown on the right, superimposed with regressed EGFR peak height over time for EGFR+ (red) and PDGFRA+ (black) clones. Branching evolution starts around EpiTrace age 0.3 and results in either EGFR+ or PDGFRA+ clones, as most evident in cluster 6 (consisting of both EGFR+ and PDGFRA+ cells).

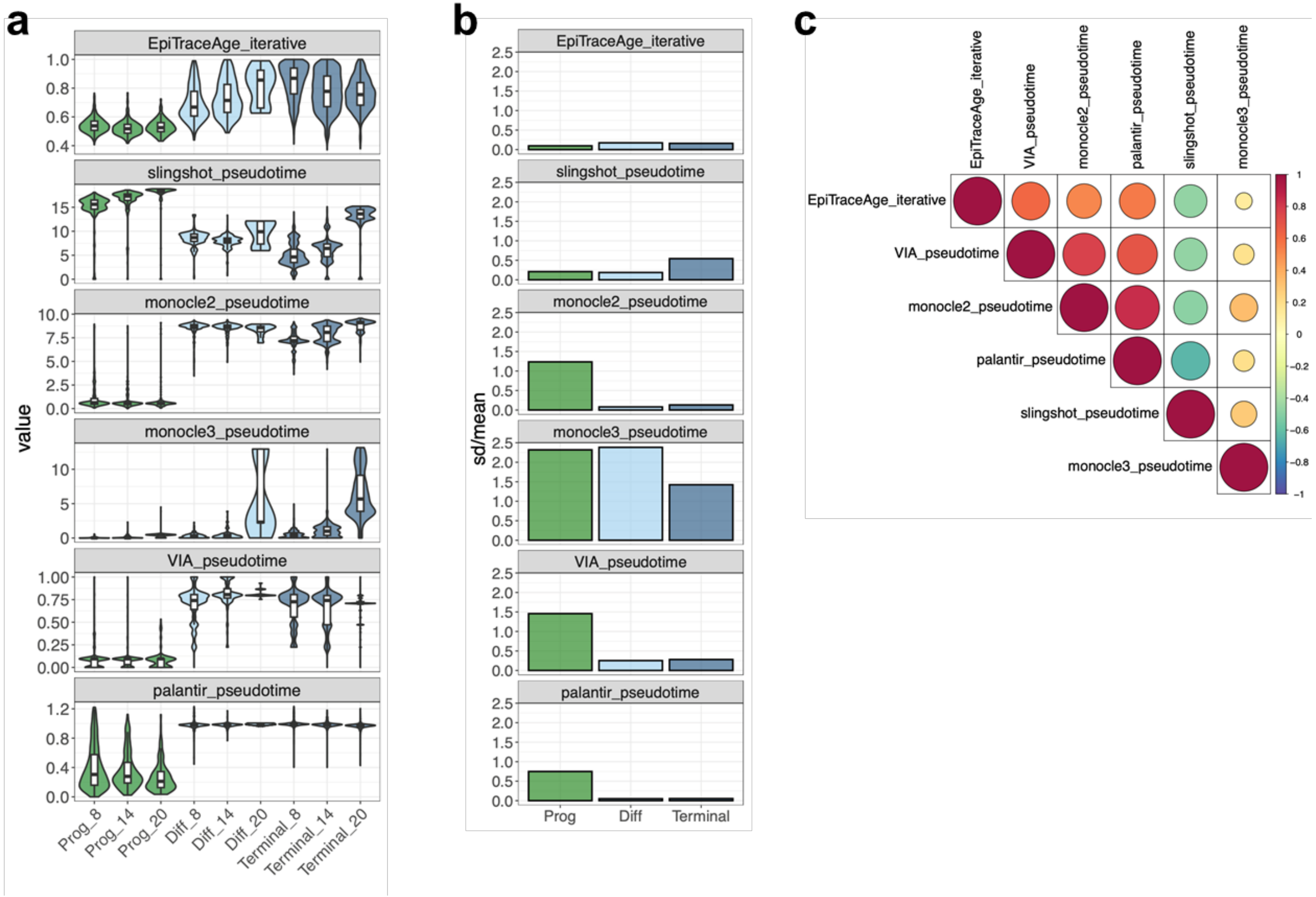


**Supplementary Figure 33. Performance comparison between EpiTrace and pseudotime-based predictors.**

**(a)** Pseudotime prediction with EpiTrace, slingshot, monocle2, monocle3, pyVIA and palantir in the mtscATAC dataset[10]. Pseudotime is grouped by cell type (Prog: progenitor, Diff: differentiated and Terminal: terminally differentiated) and cell culture day (8: Day 8, 14: Day 14 and 20: Day 20). **(b)** Variation in pseudotime (sd/mean) pseudotime prediction tools in (a).

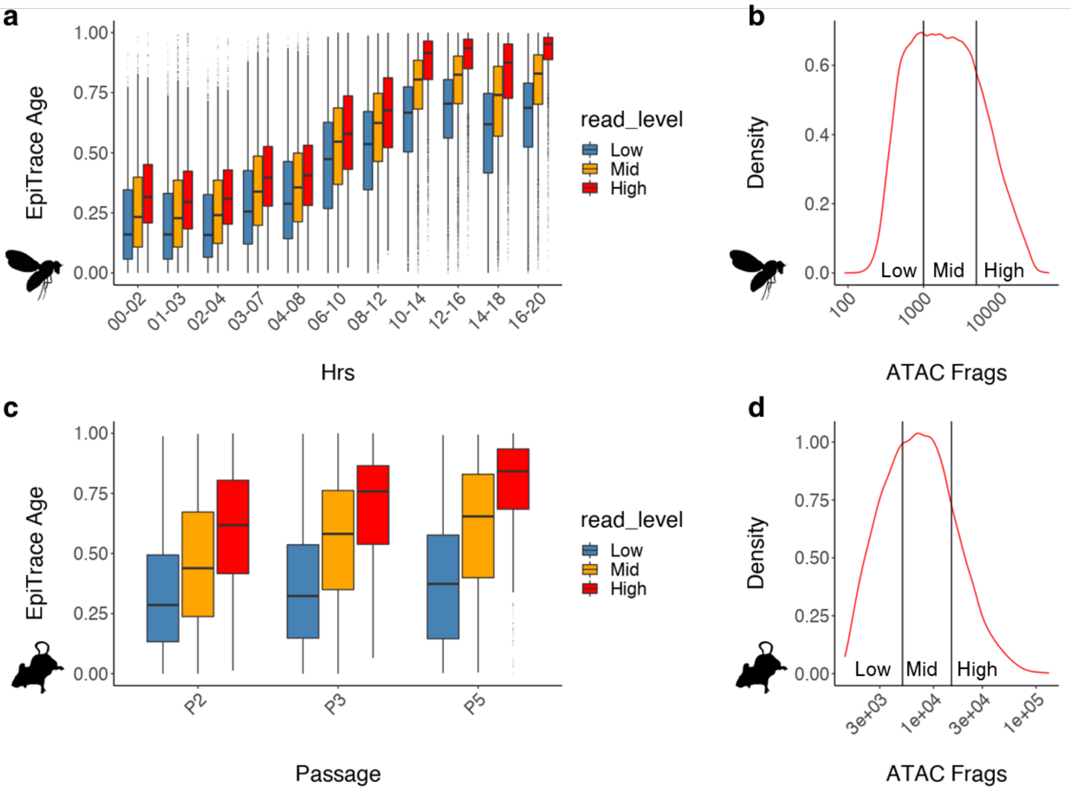


**Supplementary Figure 34. Age prediction from single cells with low, mid- or high sequencing depth.**

**(a)** EpiTrace age of low, mid- and high sequencing depth across Drosophila embryonic development time series (scATAC, GSE190130). **(b)** The classification of low, mid- and high sequencing depth groups in (a). **(c)** EpiTrace age of low, mid- and high sequencing depth across pMEF in vitro passages (pMEF SHARE-seq, in this study). **(d)** The classification of low, mid- and high sequencing depth groups in **(c)**.

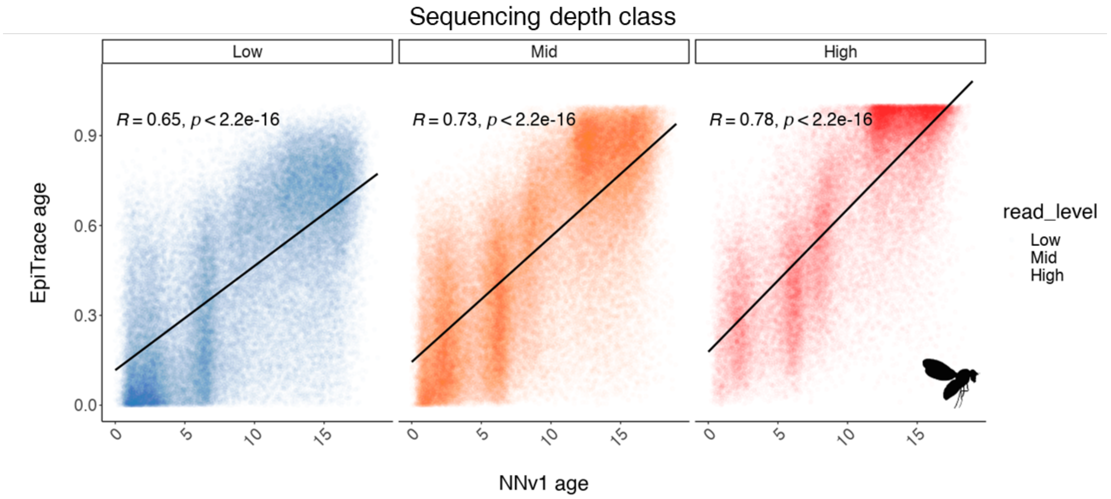


**Supplementary Figure 35. Correlation of EpiTrace age prediction and NNv1-predicted biological age from single cells with low, mid- or high sequencing depth.**

Correlation of NNv1 inferred single cell embryonic age and EpiTrace age in low-, mid- and high-sequencing depth groups in the fly scATAC dataset as in Supplementary Figure 34. Correlations: Pearson's.

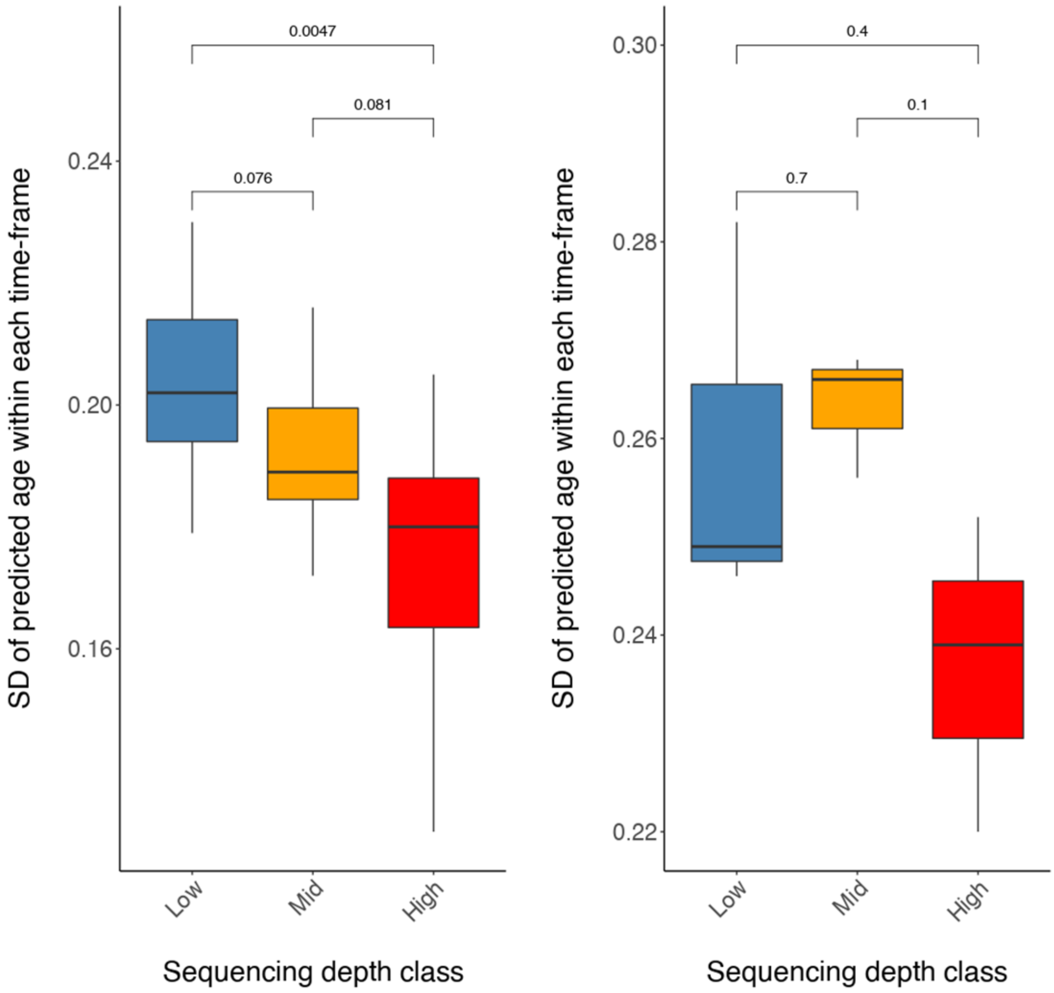


**Supplementary Figure 36. Standard deviation of predicted single cell age within each known timeframe from single cells with low, mid- or high sequencing depth.**

The standard deviation of EpiTrace inferred age in low-, mid- and high-sequencing depth groups for mouse and fly single cells is shown in Supplementary Figure 34. Statistics: t-test.

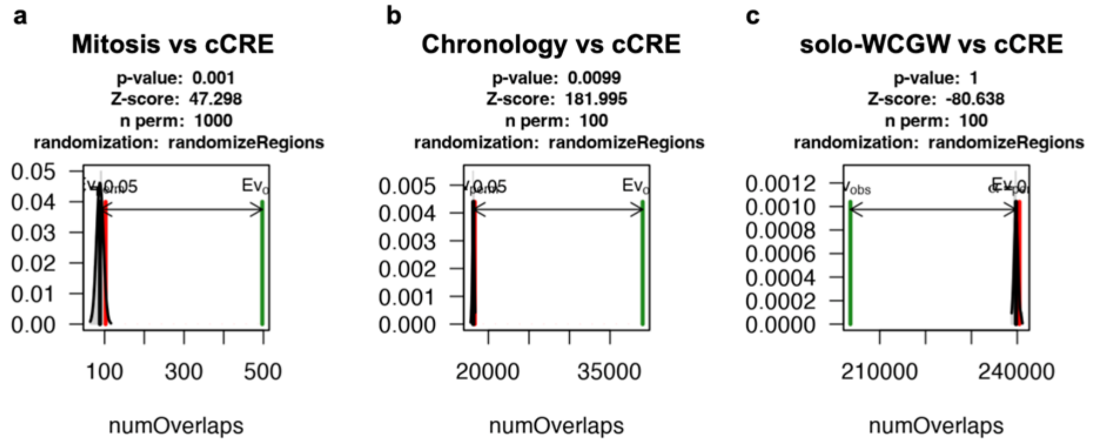


**Supplementary Figure 37. Randomized sampling test for enrichment of mitosis ClockDML, chronology ClockDML, and solo-WCGW ClockDML in canonical cis-regulatory open regions (cCRE).**

Simulated randomized sampling was performed by regioneR::overlapPermTest. cCRE is defined as in[11] the pan-human scATAC atlas.

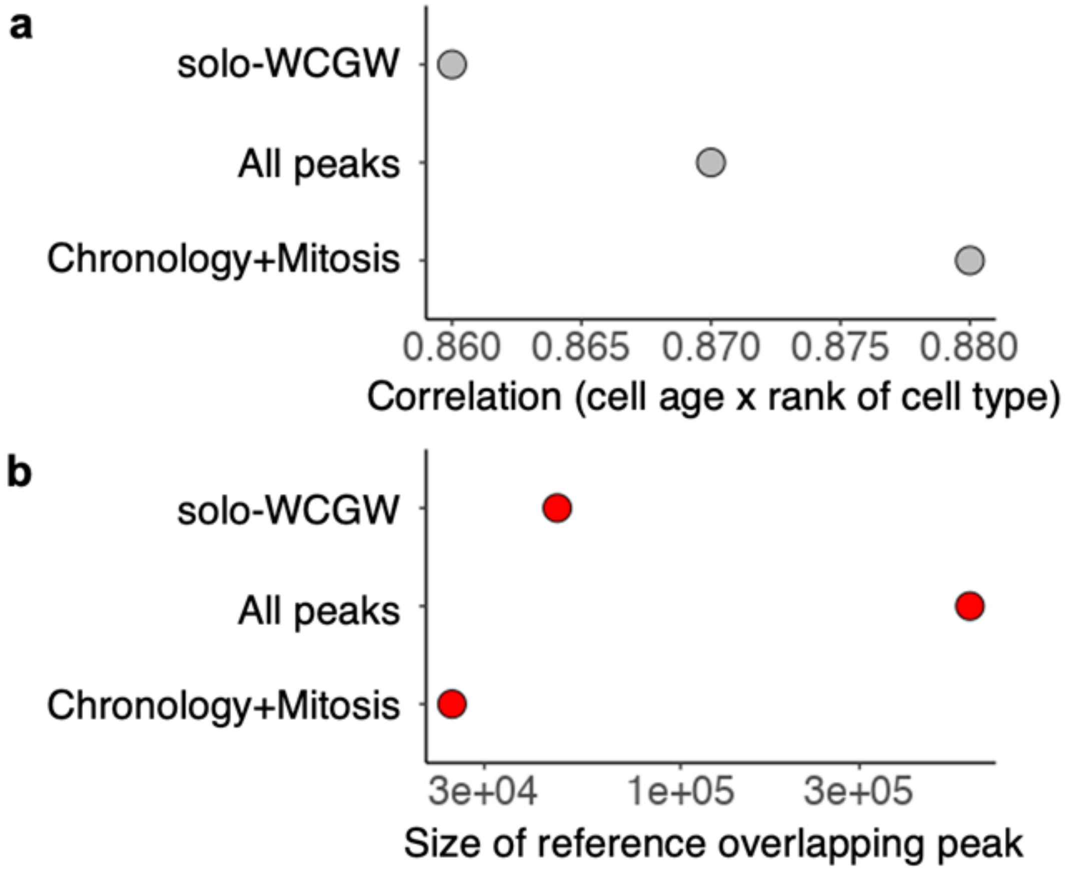


**Supplementary Figure 38. EpiTrace inference using solo-WCGW compared to other reference sites.**

**(a)** Sample age x expected age (cell rank) in the human blood hematopoiesis bulk ATAC-seq dataset. **(b)** The size of reference-overlapping peaks in the analysis in (a)

### List of Supplementary Datasets

**Supplementary Dataset 1.** Human ClockDML identified in this study.

**Supplementary Dataset 2.** Datasets used in this study, and quality control basic statistics for datasets with raw data available.

**Supplementary Dataset 3.** sgRNA designed for ClockDML G8 set.

**Supplementary Dataset 4.** Oligos used for SHARE-seq on MGI platform.
